# Identification of novel anti-bacterial compounds from a chemically highly diverse pathways-based library using phenotypic screens in amoebae host models

**DOI:** 10.1101/497032

**Authors:** Sébastien Kicka, Nabil Hanna, Gianpaolo Chiriano, Christopher Harrison, Hajer Ouertatani Sakouhi, Valentin Trofimov, Agata Kranjc, Hubert Hilbi, Pierre Cosson, Leonardo Scapozza, Thierry Soldati

## Abstract

*Legionella* pneumophila and tubercular *Mycobacteria* are the causative agents of potentially fatal diseases due to their pathogenesis but also to the emergence of antibiotic resistance that limits treatment strategies. The aim of our study is to explore the antimicrobial activity of a small ligand-based chemical library of 1,255 structurally diverse compounds. These compounds were screened in a combination of three assays, two monitoring the intracellular growth of the pathogenic bacteria, *Mycobacterium marinum* and *L. pneumophila*, and an additional anti-virulence “plaque” assay for *M. marinum*. We set up these assays using two amoeba strains, the genetically tractable *Dictyostelium discoideum* and the free-living amoeba *Acanthamoeba castellanii*. In summary, sixty-four compounds showed anti-infective/anti-virulence activity in at least one of the 3 assays. The intracellular assays hit rate varied between 1.7% (n=22) for *M. marinum* and 2.8% (n=35) for *L pneumophila* with 7 compounds in common between both pathogens. In parallel, 1.2 *%* (n= 15) of the tested compounds were able to restore *D. discoideum* growth in presence of *M. marinum* spiked in a lawn of *Klebsiella pneumoniae*. We also validated the generality of the hit compounds identified using the *A. castellanii-M. marinum* anti-infective screen in the powerful *D. discoideum-M. marinum* host-pathogen model. The characterization of anti-infective and antibacterial hits in the latter infection model revealed compounds able to reduce intracellular growth more than 50% at 30 μM. Our studies underline the relevance of using a combination of low-cost and low-complexity assays with full 3R compliance associated with a rationalized focused library of compounds to help identifying new chemical scaffolds and dissect some of their properties prior to run further compounds development steps.

## INTRODUCTION

Nowadays, the burden of antibiotic-resistant bacteria is reaching a critical point. Classical strategies to identify new antibiotics, based on their inhibitory effect on bacterial growth in broth, outside the host animal model or human, which were successful during the 50-60’s to identify the main antibiotic classes used today, are now reaching their limit. Indeed, the dramatic lack of *in vivo* confirmation of promising chemical scaffolds identified *in vitro* and/or against validated molecular targets, due to pharmacokinetic or toxicity problems at later animal and clinical stages, have given a decisive impulse to design new experimental strategies for the screening procedure ‘per se’, as well as for the design of the chemical library (Pethe et al. 2010). In addition, the development of new curative treatments against pathogenic bacteria, coupled to rationalized political choices constitute a major challenge for the future of public health (Carlet, Rambaud, and Pulcini 2014; Perez et al. 2015).

Over the years, millions of compounds have been synthesized or extracted from natural sources worldwide and are now available for biological screens (Diop et al. 2018; Farnsworth et al. 1985). The general concept behind the re-screening or repurposing of compounds with new assay systems is that small molecules have an intrinsic characteristic to interact with different targets with different potency and that an identified chemical scaffold can be developed for a new indication. At the same time, new phenotypic screening methodologies have been established, allowing the detailed study of small molecules interfering with host-pathogen interactions (Wambaugh et al. 2017). These types of assays are amenable to low or medium throughput screens. Taking into account the availability of compounds and the existence of new assays, two strategies can potentially be followed. The first one is based on random screening of the millions of compounds, while the second is based on screening a selection of compounds that form a representative set of the millions of compounds and are enriched for potential hits using a virtual screen approach (Westermaier, Barril, and Scapozza 2015). Random, high throughput screening (HTS) campaigns yields a hit rate of ~1% and are cost intensive, while screening the selected database yields similar hit rates at a lower cost, with a maximized chemical backbone diversity, allowing the use of low to medium throughput screening systems that are more complex (Macarron et al. 2011). Indeed, for the design of such small, highly diverse libraries, chemical information scientists have identified unique scaffolds by analysing the chemical diversity of all the available compounds. Furthermore, microbiology provides information on the pathways and their ligands involved in host-pathogen interactions that allow enriching the highly diverse library with compounds possessing a pharmacophore known to interact with targets of these pathways (Loregian and Palu 2013).

Among pathogens, intracellular bacteria have the ability to colonize host cells by hijacking specific defence mechanisms, including the bactericidal phago-lysosomal degradation pathway (Gopaldass et al. 2012; Dunn et al. 2017; Cardenal-Munoz et al. 2017). In addition, some bacteria exploit host resources to survive longer and proliferate in this hostile environment (Barisch and Soldati 2017). In the present study, we focused on two pathogenic bacteria that currently represent a human threat: *Legionella pneumophila*, the main causative agent of common pneumonia, which can evolve in the severe form of legionellosis known as the legionnaire’s disease (Cunha, Wu, and Raza 2015), and *Mycobacterium marinum*, a close relative to *Mycobacterium tuberculosis* (Mtb) that causes tuberculosis, a major health burden in human populations (Dheda, Barry, and Maartens 2016). The latter two pathogens are considered facultative intracellular species because of their capacity to grow within the host cell and also in the extracellular space or the environment.

Free-living amoebae (FLA) naturally present in soil and water are predatory for many bacteria and fungi that they ingest by phagocytosis (Scheid 2014; Cosson and Soldati 2008). Indeed, amoebae share ecological niches with most bacteria, and are putative cellular reservoirs for pathogenic bacteria (Greub, La Scola, and Raoult 2004; Hilbi et al. 2007). In addition, FLA have been described as ‘trojan horses’ for many pathogens including *Legionella* and *mycobacteria* species (Molmeret et al. 2005). However, FLAs such as *A. castellanii* that represent powerful cellular models used for bacterial infection studies, lack the strong genetic tools necessary to decipher key host factors. The social amoeba *Dictyostelium discoideum* is a widespread and powerful eukaryotic laboratory model, as illustrated by the recent work by Bravo-Toncia and colleagues, which emphasized its advantages as pseudo-pluricellular model organism to perform an *in vivo* screen of anti-virulence compounds against *Pseudomonas aeruginosa* (Bravo-Toncio et al. 2016).

At the cellular level, the two intracellular bacteria used in the present study, *L. pneumophila* and *M. marinum* subvert cellular compartments and machineries to establish a permissive replication niche. After their uptake by the host cell, each pathogen develops specific strategies. *M. marinum*, like *M. tuberculosis* induces phagosome maturation arrest and restrict its acidification. Then, the phagosome becomes an active interface between the host cell and the mycobacteria, by interacting with host machineries such as autophagy, endosomal and other compartments, and finally loses its integrity, giving *M. marinum* access to the host cytosol (Cardenal-Munoz et al. 2017). On the other hand, *L. pneumophila* establishes a structurally unique spacious vacuole, the LCV, that shares characteristics with the host endoplamic-reticulum compartment (Escoll et al. 2013; Prashar and Terebiznik 2015). Indeed, to establish the LCV, *Legionella* deeply modifies the host vesicular traffic by delivering, via a type-IV secretion system, a panel of effectors interacting with host components (Simon and Hilbi 2015). In addition, the fates of each pathogen are also different. *L. pneumophila* resides and proliferates in the LCV and bacteria are possibly released by host cell lysis, while *M. marinum* escapes to the cytosol and finally egresses the host cell using various routes including a specific structure named ‘ejectosome’ (Hagedorn et al. 2009).

As a response, host cells developed strategies to fight against various pathogens. Protozoa use their intrinsic defences to recognize, contain or kill pathogens (Cosson and Soldati 2008; Dunn et al. 2017). In addition, modulation of host cell processes such as autophagy (Songane et al. 2012), metabolism (Gouzy, Poquet, and Neyrolles 2014), ion homeostasis (Bozzaro, Buracco, and Peracino 2013; Botella et al. 2011) and programmed cell death (Briken, Ahlbrand, and Shah 2013) have been shown to participate in the clearance of bacteria. In particular, *Dictyostelium discoideum* has become an important model organism to study the cell biology of professional phagocytes due to many shared molecular features with animal macrophages (Dunn et al. 2017). The broad range of existing genetic and biochemical tools, together with its suitability for cell culture and live microscopy, make *D. discoideum* ideal to investigate cellular mechanisms of cell motility, differentiation, and morphogenesis (Nichols, Veltman, and Kay 2015; Loomis 2015). Furthermore, *D. discoideum* possesses evolutionary conserved pathways involved in pathogen recognition and defences that are highly similar to macrophage ones (Bozzaro, Bucci, and Steinert 2008). In particular, thanks to the extreme conservation of phagosomal composition and function with human phagocytic cells (Boulais et al. 2010), *D. discoideum* is used as a recipient cell to investigate interactions with pathogenic bacteria such as *Salmonella, Mycobacteria, Legionella* or *Pseudomonas*, (Solomon, Leung, and Isberg 2003; Hagedorn and Soldati 2007; Steinert and Heuner 2005).

In recent year, several phenotypic screens performed in the context of infected animal host cells have identified compounds active against pathogenic mycobacteria. Interestingly, elucidation of the mode of action of hit compounds revealed emerging concepts in the field of host-pathogen interactions, demonstrating that intracellular bacteria proliferation can be affected by targeting bacterial metabolic pathways key to their intracellular life, but also by modulating host pathways (VanderVen et al. 2015; Sundaramurthy et al. 2014). Alternative *in cellulo* screening systems have also been established to take advantage of previously underexplored amoeba host-models (Kicka et al. 2014; Harrison et al. 2013). For example, *D. discoideum* and *A. castellanii* were used to characterize compounds of the GSK TB-set of anti-mycobacterial compounds (Trofimov et al. 2018). This study showed that most compounds previously selected for their antibiotic activity against *Mtb* and *M. smegmatis* (Ballell et al. 2013) were active against the closely related *M. marinum*, but showed little or no activity in the intracellular context of an infection. Most importantly, it demonstrated that compounds with anti-infective activities were similarly active in the *M*. marinum-Amoebae system and the more standard Mtb-macrophages models. This study also underlined the relevance of using evolutionary distant pathogen and host models to reveal conserved mechanisms of virulence and defence (Trofimov et al. 2018).

Here, we tested compounds from a proprietary library derived from a unique ligand-based virtual screening in a combination of three assays to determine their anti-infective properties in the two infection models *A castellanii* - *L. pneumophila* and *A. castellanii - M. marinum* for their abilities to inhibit intracellular growth. In parallel, the “anti-virulence” activity of the compounds was determined by monitoring their capacity at reverting the growth arrest of *D. discoideum* on lawns of food bacteria spiked with pathogenic *M. marinum*. Finally, the potency, specificity, and toxicity of the hit compounds was evaluated.

## RESULTS

### Design of a chemically highly diverse pathway-based library of compounds

*Legionella* and *Mycobacterium* interact with a host through different pathways in which we can search for potential drug targets. Instead of focusing on a single target for a drug design, we have explored the pathways involved in the host-pathogen interaction process, and not only those having a significant pathogen-host selectivity ratio. We selected in total 18 host and pathogen pathways as potential pharmacological targets. Ligands/metabolites known to interfere/interact with these host and pathogen pathways were identified from the literature, and used as queries or search templates to prepare pharmacophores for launching a campaign of ligand-based virtual screening (LVS) of the ZINC database (http://zinc.docking.org/) (Sterling and Irwin 2015) using ROCS, a tool from the OpenEye software package (http://eyesopen.com/rocs) (Swann et al. 2011).

Subsequently, we applied the following workflow (Figure 1): i) we screened the ZINC lead-like database composed of 2.5 million compounds saving the 25,000 best hits for each query; ii) we ranked the hits according to the ROCS TanimotoCombo score; iii) we selected the first hit and the following, if they passed a test of structural dissimilarity, using the Lingo method program (Vidal, Thormann, and Pons 2005) thereby increasing the chemical diversity and maximizing the coverage of the chemical space of the ZINC lead-like database; iv) we chose at most 2 analogues per series from each screened pathway, saving 100 selected compounds to the pool of potential hits composing the physical library for the experimental screen. The VS of the 18 different pathways yielded ~1,800 compounds of which 1,255 were purchased to compose the final highly diverse pathways-based library named Sinergia library (Slepikas et al. 2016).

**Figure 1.**
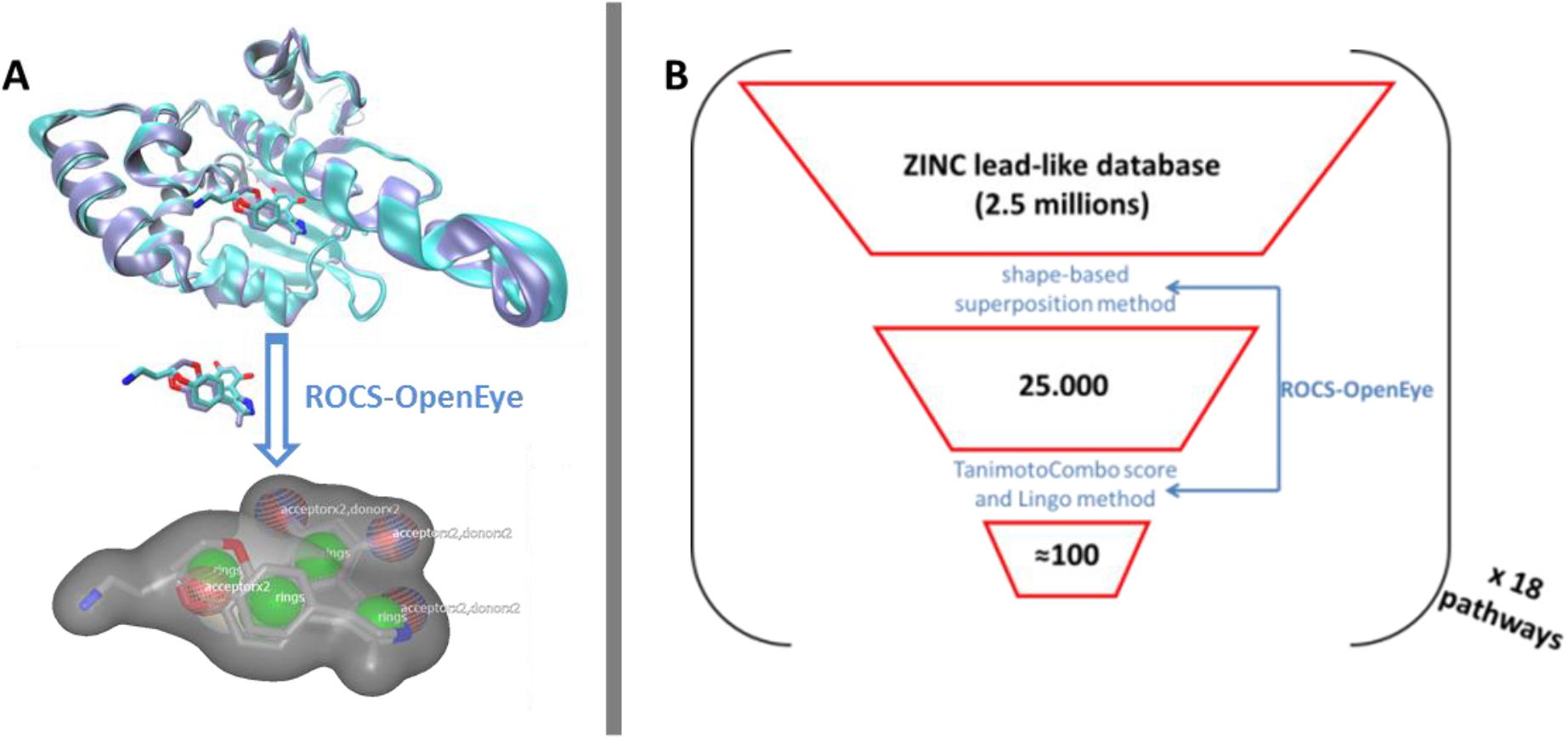
Schematic representation of the LVS workflow applied per each pathway. i) ligand-based virtual screen of the ZINC lead-like database; ii) ranking of the 25,000 best virtually screened hits according to the ROCS TanimotoCombo score; iii) selection of the first hit and then each next one if structurally dissimilar to already chosen ones by using the Lingo method; iv) we chose at most 2 analogues per series from each screened pathway saving 100 selected compounds to the pool of potential hit compounds composing the physical library for the experimental screen.

### Characterization of the pathways-based library properties: chemical diversity and drug likeness

The chemical diversity of the pathways-based library was investigated and analyzed on the basis of the Z-matrix calculated according to Tanimoto’s chemical similarity metrics (*T_sim_*; see Figure 2) using Canvas, a tool from the Schrödinger software package (Sindhikara and Borrelli 2018). The corresponding heat map drawn using Netwalker1.0 (Komurov et al. 2012) clearly shows that the library is highly diverse (Figure 2). We deliberately limited the number of analogues in the primary library to concentrate on the validated hits selected from the different screens. The analysis of the drug likeness of the compounds composing the library and the comparison with the known drug-likeness standards (Lipinski and Hopkins 2004; Lipinski 2016; Lipinski et al. 1997) reveals that the waste majority of the compounds have the properties to be drug-like (Figure 3).

**Figure 2.**
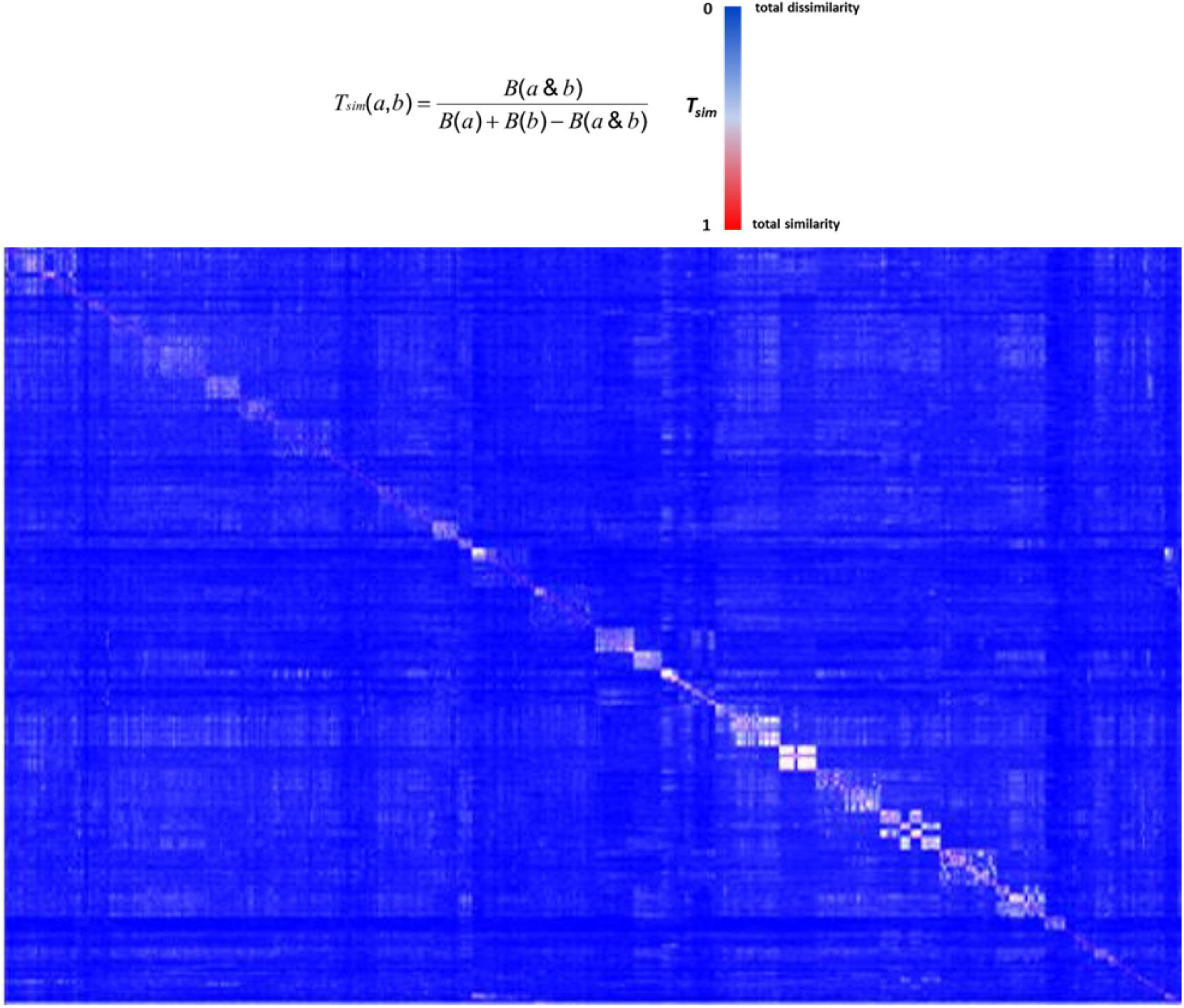
The heat map of pathways-based library. The Z-matrics was calculated according to Tsim values: T_sim_ = 0 for total dissimilarity (blue); T_sim_ = 1 for total similarity (red).

**Figure 3.**
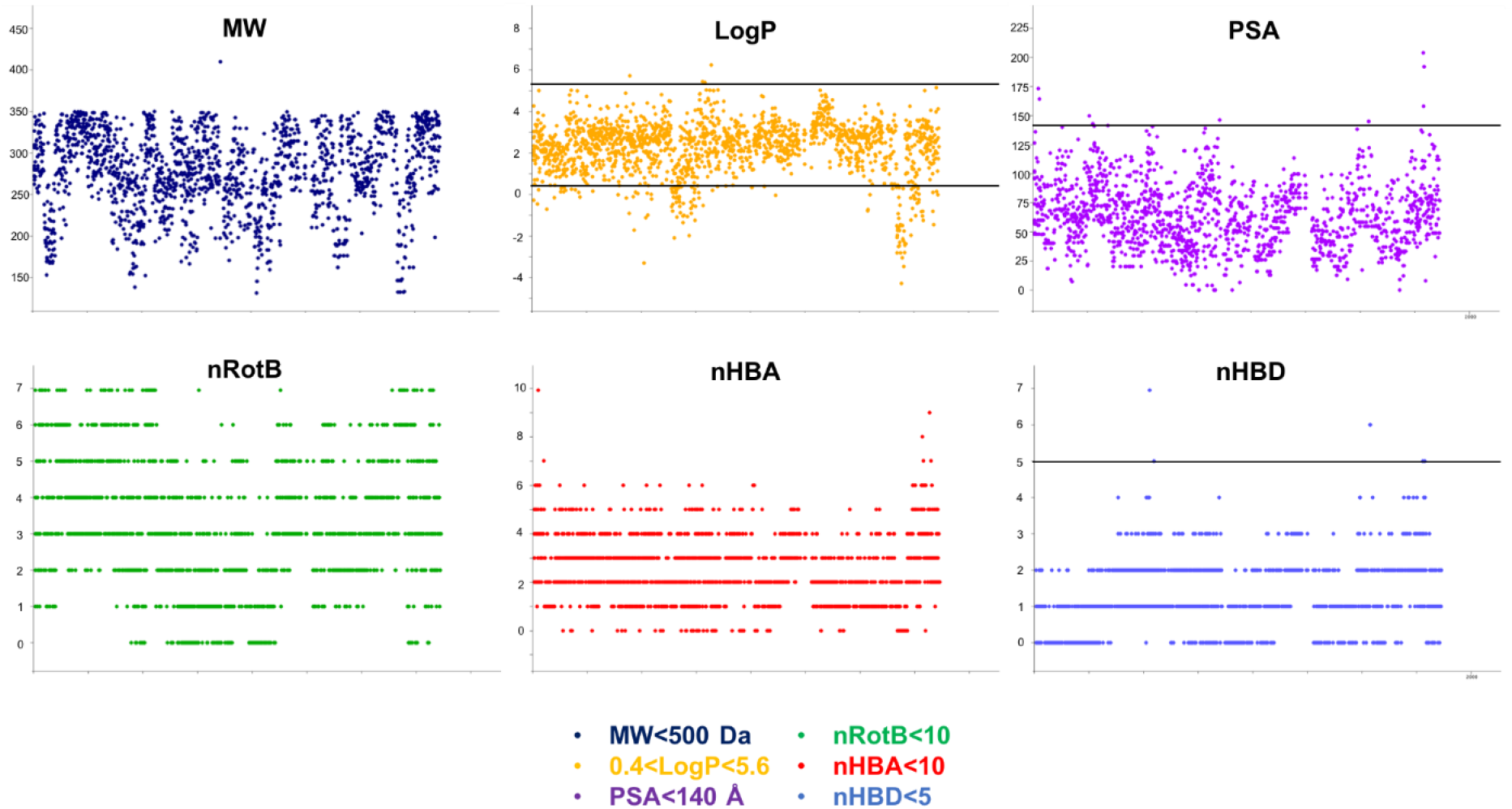
Physico-chemical prediction of the Sinergia Library. The prediction descriptors of the library was performed using the cheminformatics package Canvas to evaluate the drug-likeness properties of library’s compounds (Ghose, Viswanadhan, and Wendoloski 1999).

### Screens characteristics

1,255 compounds were assayed using our three biological phenotypic screens (see materials and methods). The categories of pharmacophore queries are indicated in Table 1, and the hallmarks of the various screens are summarized in Table 2.

**Table 1.** Host and pathogen metabolic pathways used for selecting the query molecules. The corresponding pharmacophores of the queries molecules have been used as reference molecules for the ligand-based virtual screen. Each pathway-based screen resulted in 100 new potential hits.

| Cellular pathways & targets | Query pharmacophores | References |
| --- | --- | --- |
| Calcineurin 1 | Pyrazolopyrimidines, CN585 | (Jayaraman et al. 2010) (Sieber and Baumgrass 2009) |
| E3-Ligase | HLI98 | (Yang et al. 2005) |
| AKT-PKB | Indoles, pyridopyrimidine, pyrazolopyrimidine, pyranonaphthoquinones, quinolines, azacarabazoles, oxazole-2- amines | (Tardy et al. 2014) |
| PI3K | Wortmanin, LY294002, staurosporin, ATP, myricetin, quercetin | (Walker et al. 2000) |
| Autophagy | Small molecule enhancers of rapamycin = SMERs: SMER18 and 28 from ref. Floto, 2007. Diphenylbutylpiperidines – cpds 15d, 15i, 15j from ref Chen, 2011 | (Sarkar et al. 2007) (Floto et al. 2007) (Chen et al. 2011) |
| ATP-ase | Salicylihalamide A, indole derivatives --- H<sup>+</sup>-ATPase; diaryl pyrazole resorcinol derivatives --- HSP90 ATPase; adociasulfate-2 --- kinesin ATPase | (Design 2009; Supino et al. 2008); (Sharp et al. 2007; Sakowicz et al. 1998) |
| CD36 | Cilostazol, Sulfo-N-succinimidyl derivatives | (Coort et al. 2002) |
| ABC-transporters | Verapamil, LY335979, OC144-093 | (Sharom 2008) |
| Trp | CdRP+ IGP; ATB107 --- IGPS target | (Hennig et al. 2002) (Shen et al. 2009) |
| Cys | WNI tripeptide; 3-hydroxybutanoic acid, 5-oxohexanoic acid, 4-methyl valeric acid --- O-acetyl serine synthetase target | (Salsi et al. 2010) |
| His | PRPP; PR-ATP; cpds 4 6 17 18; dicoumerol --- ATP-PRT target | (Cho, Ioerger, and Sacchettini 2008; Dall-Larsen, Kryvi, and Klungsoyr 1976) |
| Gln | Purine analog; --- glutamine synthetase target | (Nilsson et al. 2009) |
| Lipase | Orlistat, glimepiride | (Bertrand et al. 2010; Schalk-Hihi et al. 2011; Christelle et al. 2011; Pedicord et al. 2011; Bustanji et al. 2011) |
| FAS-II | Triclosan, Triclocarbon | (Lu, Huang, and You 2011) |
| Cations pumps | Acetazolamide, Zonisamide, Topiramate, Phenytoin, Lamatrigine, Sumatriptan, Verapamil | (Payandeh et al. 2011) (Long, Campbell, and Mackinnon 2005; Wu et al. 2010) |
| Proteasome | Lactacystin and omuralid, MG-132, Morphine | (Yang et al. 2008; Yoshida et al. 2009) |
| MeC | Spongotine A, Bromotopsentin, Bromodeoxytopsentin, cis-3,4-dihydrohamacanthin B | (Zoraghi et al. 2011) |
| Porins | Cadaverine, Putrescine, Spermine | (Iyer et al. 2000) |

**Table 2:** Screen assays characteristics.

| **A** | **Anti-infective screens** |  | **Anti-virulence screen** |
| --- | --- | --- | --- |
| Host | *A. castellanii* & *D. discoideum*. | *A. castellanii* | *D. discoideum* |
| Pathogen | *M. marinum* | *L. pneumophila* | *M. marinum* (mixed 1:50 with *K. pneumoniae* ) |
| Duration | 2-3 days | 2 days | At least 1 week |
| Read-out (type) | Intracellular bacterial growth (GFP reporter) (quantitative) |  | Host growth on mixed bacterial lawns (semi-quantitative) |
| Compound concentration | 30 μM (solution) | | 10 μM (agar) |
| **B** | **Hit categories** |  |  |
| Anti-infective | yes | yes | Possibly <sup>a</sup> |
| Anti-virulence | yes | yes | yes |
| Pro-infective | yes | yes | no |
| Antibiotic | yes<sup>b</sup> | yes<sup>b</sup> | yes (mycobacteria- specific) |
| Defence booster | yes | yes | yes |

<sup>a</sup>) Further investigations are needed to disentangle whether the activity of the compound is affecting the extracellular bacteria or the infecting ones
<sup>b</sup>) The anti-infective screen on *M. marinum* detected mainly hits with low antibiotic activity, whereas the screen on *L. pneumophila* detected mainly potent antibiotics.

Each of the three assays was established and optimized separately prior to testing the 1,255 compounds (Harrison et al. 2013; Kicka et al. 2014; Froquet et al. 2009). The three screens were run simultaneously using the same compounds batch. The compounds’ concentration was adjusted to 30 μM for the two cell infection screening assays, taking into account the shielding effect of *A. castellanii* cells on intracellular bacteria (Tosetti, Croxatto, and Greub 2014), whereas a final concentration of 10 μM was used in the anti-virulence assay monitoring *D. discoideum* growth on bacteria lawns (Table 2). Considering screen stringency variations, the threshold for hit detection was arbitrarily fixed at a minimum of 20% reduction of the intracellular mycobacteria growth, and 40% reduction for intracellular *Legionella* growth compared to the DMSO control. Fluorescence from GFP-expressing bacteria was used as a readout of intracellular growth for both anti-infective assays in *A. castellanii* host. Briefly, *A. castellanii* cells were infected with GFP-expressing *M. marinum* or *L. pneumophila*. Intracellular bacteria growth was monitored by measuring the fluorescence increase using a Synergy MX fluorescence plate reader with time-points taken every 3 hours for *M. marinum*, and at various time intervals for *L. pneumophila* as indicated in figure 4B. Extracellular *M. marinum* growth was inhibited by adding 10 μM amikacin in the medium, whereas no antibiotic was added to *L. pneumophila* since PYG media does not support the bacterial growth. Examples of the intracellular bacterial growth curves are presented in Figure 4A and B respectively for *M. marinum* and *L. pneumophila*. Normalization and analysis of the data generated in these two screens were performed as previously reported for the *A. castellanii – M. marinum* (Kicka et al. 2014; Ouertatani-Sakouhi et al. 2017) and the *A. castellanii - L. pneumophila* (Harrison et al. 2013) host-pathogen assays, respectively. In brief, normalization of bacterial growth in presence of each compound was calculated related to the DMSO carrier (=1). For the phagocytic plaque assay, a semi-quantitative visual inspection and scoring of the compounds was applied, as described previously (Froquet et al. 2009). As shown in Figure 4C, the potency of hit compounds to restore *D. discoideum* growth was evaluated to range from non-hit molecules (=0) to hits that fully restore host growth (=4). Following their identification in their respective initial screen, candidate hit compounds were independently validated at least three times in their respective assay.

**Figure 4:**
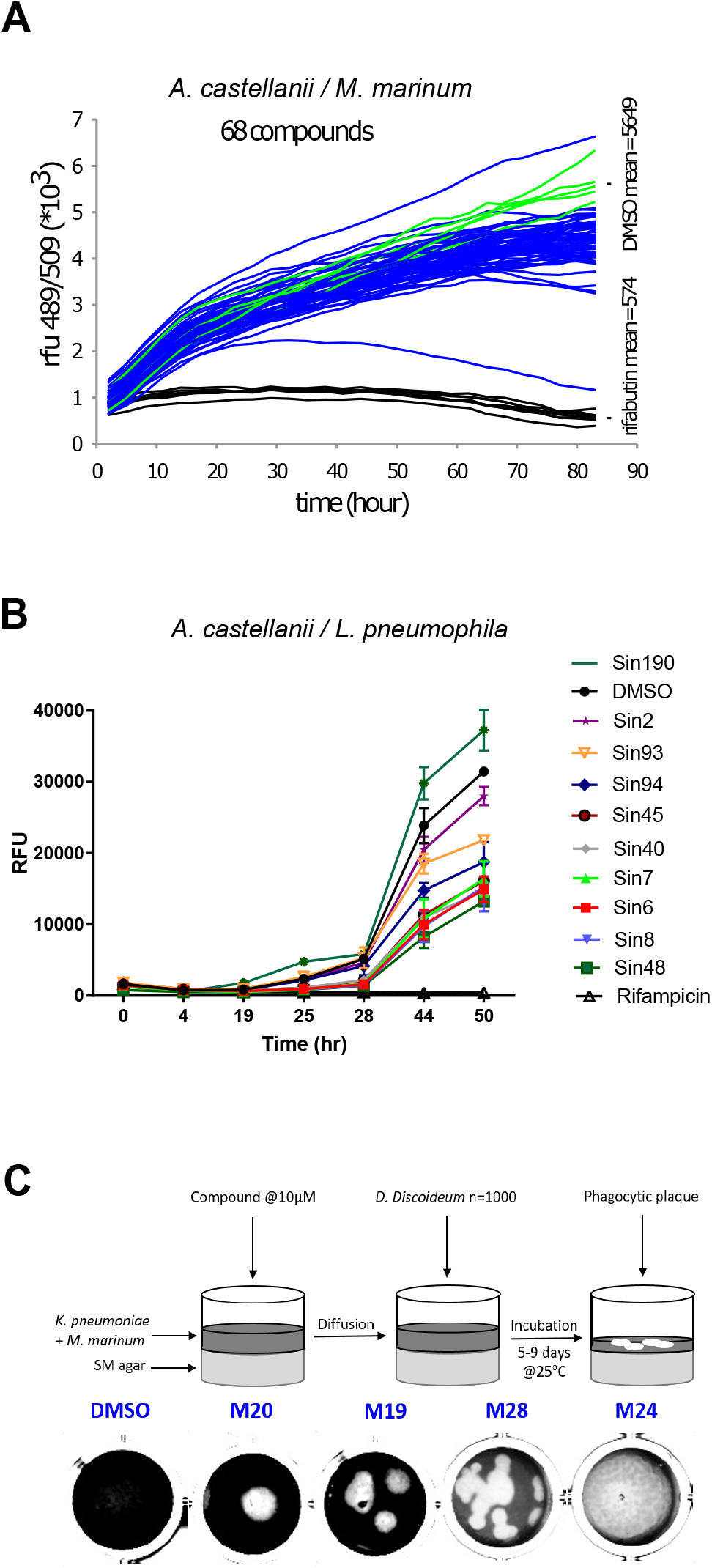
A representative selection of the Sinergia library compounds tested in the three different assays. Phagocytic host cell *A castellanii* was infected with (A) *M. marinum* msp12::GFP and (B) GFP-expressing *L. pneumophila*, 5 × 10^4^ infected cells were transferred to each well of a 96-well plate, The course of infection at 25°C was monitored by measuring fluorescence for 72 hours in the presence of 30 μM of corresponding compounds compared to DMSO control (green) and 10 μM rifabutin - only for *M. marinum* assay- (black). (C) Virulence assay. Each compound (10 μM) was added on SM-agar medium followed by the addition of *K. pneumoniae* and *M. marinum* mixture. 1,000 *D. discoideum* cells were deposited in the centre of the well and plates were incubated for 5–9 days at 25°C and the formation of phagocytic plaques was monitored visually.

### Hit frequency and overlap between screens

The two intracellular growth assays used for screening all 1,255 compounds resulted in a broad range of inhibitory activities. The hit rate was considerably higher for *L. pneumophila*, where 2.8% (n=35) of the compounds tested at 30 μM showed a growth inhibition of at least 40%, compared to 1.7% (n= 22) with at least a 20% inhibition for *M. marinum* (Fig. 5 A and B.). The *M. marinum* - *D. discoideum* phagocytic plaque assay showed that 1.2% (n=15) of the compounds at 10 μM are valuable hits and were able to restore host growth (fig. 5C). In comparison to infection assays, full restoration of amoeba growth on a bacteria lawn containing pathogenic mycobacteria appeared to be more restrictive and identified only 15 hits from the library. Taken together, sixty-four compounds showed anti-infective activity in one of the intracellular growth assay or did attenuate mycobacterial virulence. As depicted in the Venn diagram, the *A. castellanii – L. pneumophila* screen shared 8 hit compounds with the other two screens, among these 7 are common between the two infection assays and only 1 compound between the *L. pneumophila – A castellanii* model and the phagocytic plaque assay (Fig. 5D). Surprisingly, no hit was common to the two assays using *M. marinum* as pathogenic bacterium. Notably, we also identified a certain number of pro-infective compounds, namely chemicals that lead to a remarkable increase of the intracellular bacterial mass compared to the DMSO control. The *A. castellanii – L. pneumophila* assay identified only one compound that increased the intracellular growth more than 40% when compared to the DMSO control (Fig. 5B). In contrast, five pro-infective compounds were identified with at least 2-fold increase in the intracellular *M. marinum* bacterial load (Fig. 5A).

**Figure 5:**
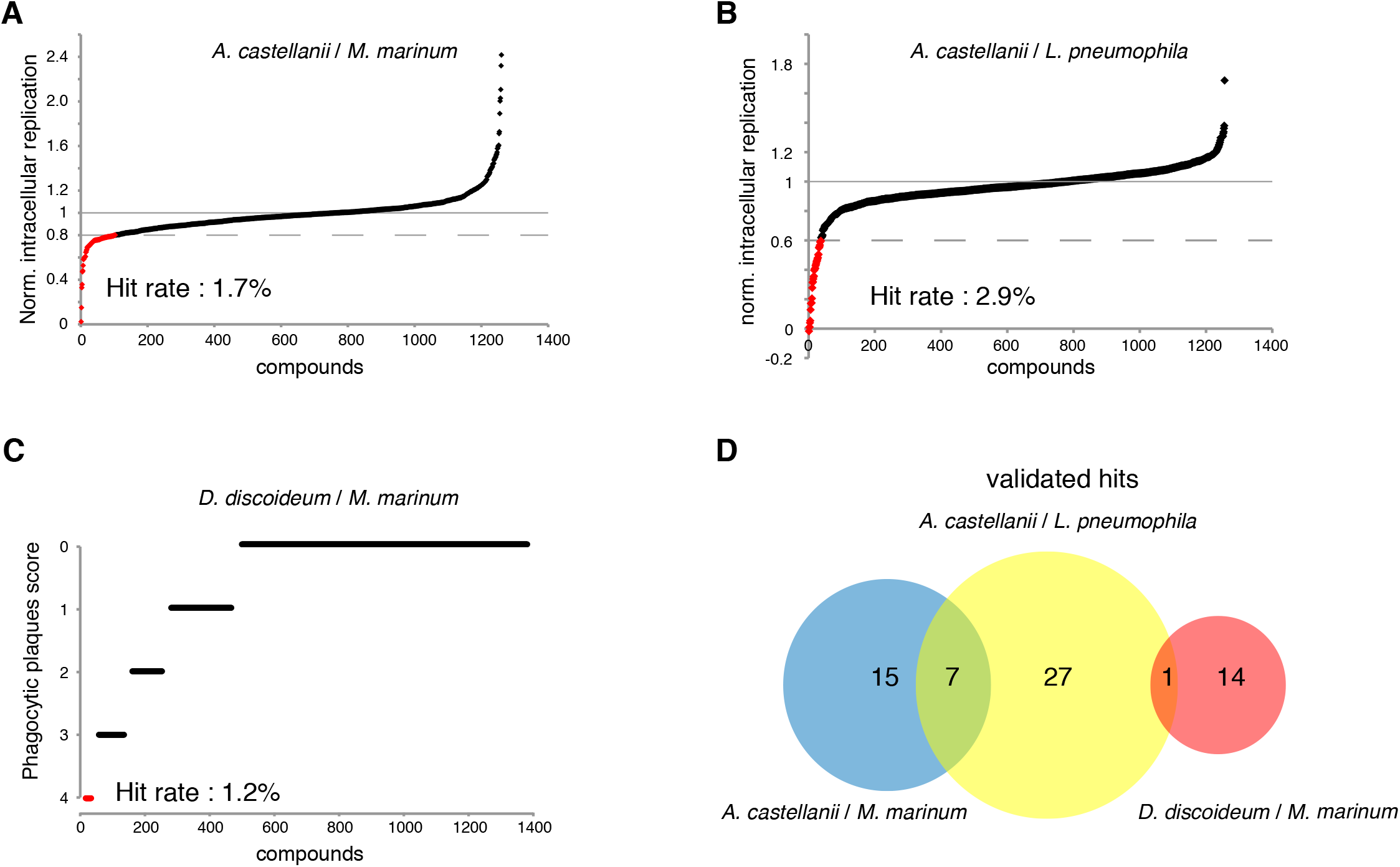
Percentage hit rate (panel A, B, C) and hits overlap of the 3 screens. All 1,255 compounds were tested for potential intracellular growth inhibition. Library compounds were plotted based on their anti-infective properties. Compounds resulting in decreased intracellular *M. marinum* and *L. pneumophila* replication by over 20% and 40% respectively at the screening concentration (30 μM) were defined as hit candidates. (C) Compounds score from the “phagocytic plaque assay”. The potency of each compound to restore *D. discoideum* growth was evaluated, hit compounds were determined as molecules that fully restore host growth (=4). (D) Venn diagram representing summarized results of the primary hits identified from the 3 assays. The analysed set included compounds that passed the aforementioned cut-offs.

### Effect of hit compounds on amoeba fitness

To evaluate the effect of hit compounds on host fitness, we used *D. discoideum* cells expressing GFP-ABD to measure toxic and growth inhibitory activities of compounds (Trofimov et al. 2018). In parallel, a cell viability test using Alamar blue was performed using *A. castellanii* 4 hours after hit compound addition at a 30μM concentration. For each assay, values were normalized to the DMSO carrier control (=1), compounds’ toxicity and growth inhibition data for each phenotypic screen are represented in Figure 6 A, B and C.

**Figure 6:**
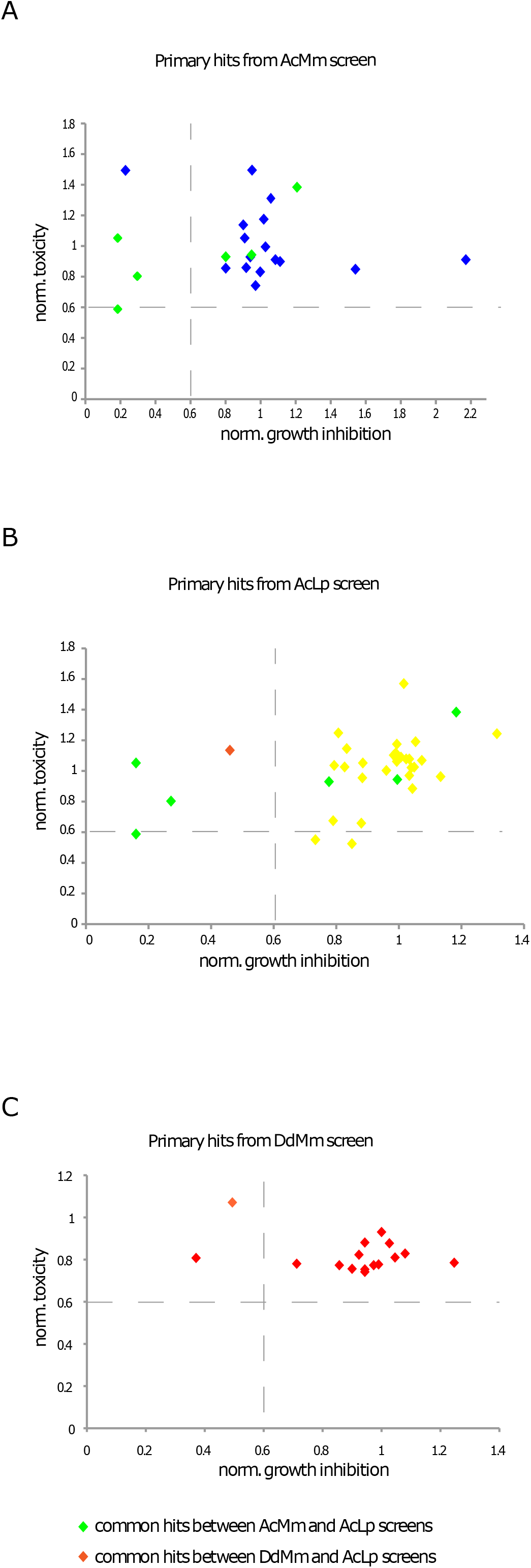
Characterization of cytotoxic and growth inhibitory activities of primary hit compounds. Growth kinetics of *A. castellani* (A, B) and *D. discoideum* (C) GFP-ABD in HL5c medium and in the presence of 30 μM of corresponding compounds values were normalized to the DMSO carrier control (=1). The analysed set included compounds that displayed inhibition of more than 60% normalized RFU. (B) Uninfected triplicate wells were treated with compounds in 100 μl LoFlo media. Plates were incubated at 30°C for 24 h, after which 10 μl Alamar Blue reagent was added, and plates incubated for a further 3 h. The fluorescence at 595 nm was measured, and data normalized between 1 (treatment with LoFlo alone) and 0 (SDS, total lysis of the cells).

Only one compound (ZINC01718072) was excluded because its strong fluorescence at the GFP emission wavelength confounded the results. At the end, five hit compounds did not pass the set threshold (more than 40% growth inhibition or 40% toxicity when compared to DMSO control) and were rejected for this deleterious effect on host fitness.

### Properties of the hit compounds

To determine whether hit compounds had antibiotic properties, we measured the activity of compounds against *M. marinum* and *L. pneumophila* growing in broth. Compounds at 30 μM in DMSO, were transferred to 96-well plates containing 10^5^ GFP-expressing *M. marinum* or *L. pneumophila* per well. Growth was monitored for 48 hours, and the total fluorescence intensity was used as a proxy to quantify the bacterial mass. In parallel, we tested the ability of hit compounds identified by the phagocytic plaque assay to directly inhibit *M. marinum* growth on agar at 10 μM. The normalized results (DMSO=1) of the hits detected during cell infection assays are shown in Figure 7 A and B for *M. marinum* and *L. pneumophila*, respectively. In summary, 19 of the 35 hit compounds against intracellular *L. pneumophila* showed an antibiotic activity, whereas only one out of twenty-one exhibited antibiotic properties against *M. marinum*. Lastly, eleven compounds out of fifteen from the anti-virulence hits against *M. marinum* were defined with mild to strong antibiotic activity when assayed for bacterial growth inhibition on Agar (Fig. 7C).

**Figure 7:**
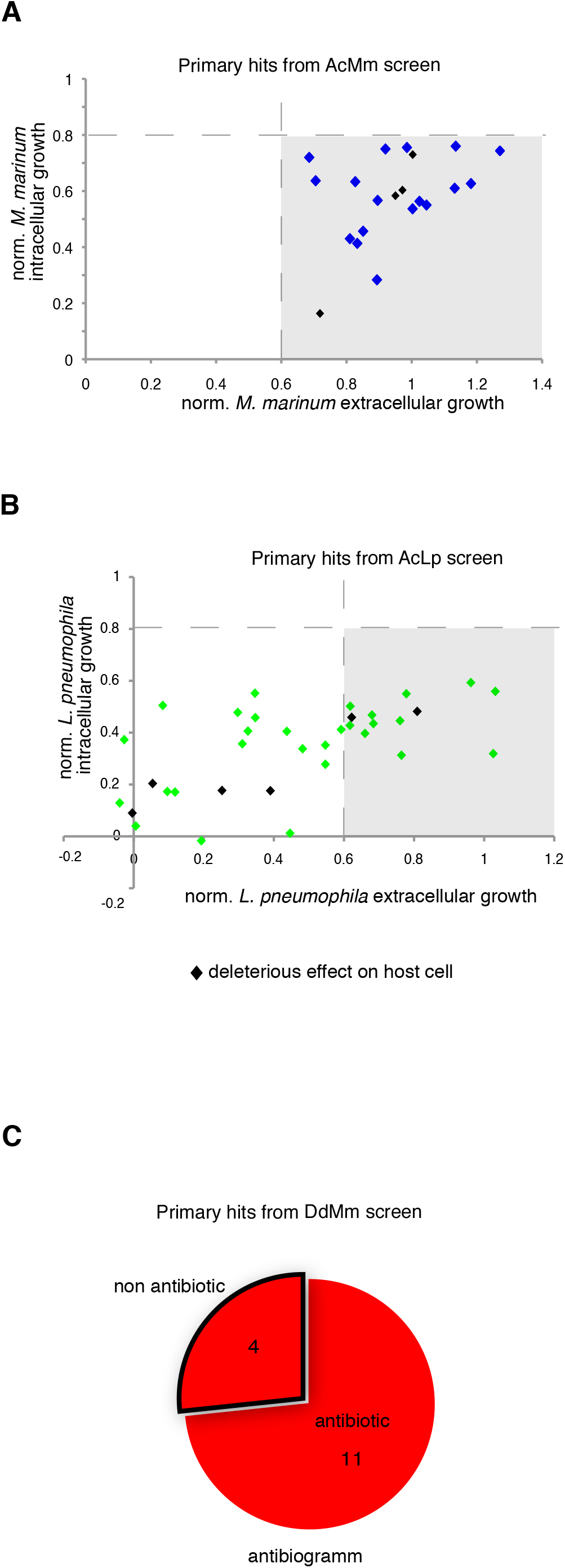
Intracellular and extracellular growth of the pathogenic bacteria in presence of the primary hit compounds. Growth of *M. marinum* (A) and *L. pneumophila* (B) in the presence of hit compounds (30 μM) was determined for both intracellular (*A. castellanii*) and extracellular replication (broth). The graphs indicate the fluorescence measurements normalized to 1 (vehicle control). (C), Antibiotic assay. Compounds (10 μM) identified in phagocytic plaques assay were added to 7H11 medium in each well, then 1,000 *M. marinum* bacteria were deposited in the well. Growth of mycobacteria was monitored after 6 days at 30°C.

### The *D. discoideum-M. marinum* host-pathogen model system

To validate the generality of the anti-infective screen performed using the *A. castellanii* - *M. marinum* system, we tested the previously identified 21 hit compounds using the *D. discoideum-M. marinum* host-pathogen model. Fourteen compounds out of twenty-one exhibited at least a 20% inhibitory activity on intracellular mycobacteria (Figure 8A). Interestingly, from the hit compounds identified in the “extracellular” anti-virulence plaque assay, almost 50% (7/15) of compounds presented a growth inhibitory effect on mycobacteria when tested in the intracellular *D. discoideum* infection model. Curiously, the antibiotic potency of these hit compounds, as detected using growth of bacteria on Agar, was barely detectable in the two-days bacteria growth assay in suspension and only 2 compound showed an interesting inhibition intra and extracellularly (Figure 8B – dots on the edge of quadrant 1).

**Figure 8:**
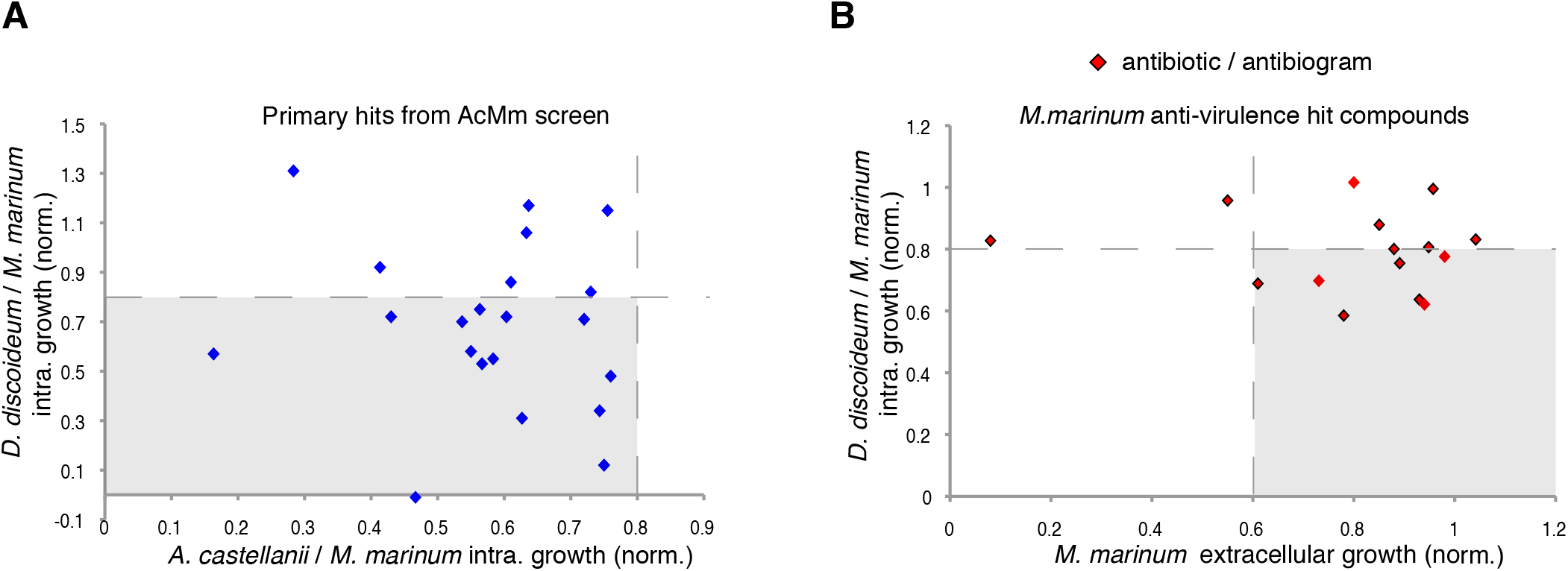
Comparison of two amoeba host models. A. Comparison of anti-infective properties of primary hits on intracellular *M. marinum* in *D. discoideum* and *A. castellanii* host cells. (B) Selected anti-virulence hits plotted based on their anti-infective (20% inhibition) and anti-bacterial (40%) activity in *D. discoideum – M. marinum* model. 10^5^ *D. discoideum* cells infected with *M. marinum* msp12::GFP were transferred to the wells, hit compounds were added at 30 μM. The fluorescence intensities were measured for 72 hours every 3 hours.

## Discussion

### Screen results reflect pathogen specificities and assays characteristics

Since mycobacterial infection is described as a complex and dynamic series of interactions between multiple host and bacterial components and pathways. We selected a total of 18 host and pathogen pathways as potential pharmacological targets, and an LVS of the ZINC database was launched to identify ligands/metabolites known to interfere/interact with these pathways. In our study, we tested 1,255 compounds from the ZINC lead-like database derived from a unique ligand-based virtual screening to determine their anti-infective properties. For this purpose, a combination of three phenotypic assays was used, as summarised in Table 2. Two assays to monitored the intracellular growth of 1) *M. marinum* and 2) *L pneumophila* in *A. castellanii* and identified antibiotics and potential anti-virulence and host defence boosters. In parallel, we used a phagocytic plaque assay 3) in which a compound restoring growth of *D. discoideum* on a lawn of *K. pneumophila* spiked with *M. marinum* has either selective antibiotic activity against M. *marinum*, or attenuates the virulence of infecting *M. marinum*.

We identified 64 compounds showing activity in at least one of the assays for intracellular bacteria growth inhibition, or attenuation of mycobacterial virulence (table S1). Although the same host was used for the two infection assays, only few hits were common against *M. marinum* and *L. pneumophila*. The number of identified hit compounds was considerably higher for *L. pneumophila* compared to *M. marinum* (Figure 5A and 5B). While no strong antibiotic hit was identified using the *A. castellanii - M. marinum* model, almost half of the hits identified in the *A. castellanii - L. pneumophila* screen (16/35) are potent antibiotics. A possible explanation might be the difference in growth rate between the two bacteria. *L. pneumophila* grows almost unrestrictedly inside the amoeba, with a doubling time close to 60 min, whereas *M. marinum* is a considerably slower grower, with doubling time of round 8 hours. Another related aspect is the temperature (T=25°C) used for the intracellular assays which is closer to the optimum for *L. pneumophila*. Another plausible reason might be that the two pathogens have a very distinct cell wall composition. In contrast to the Gram-negative *L. pneumophila* cell wall, *M. marinum* has an elaborate and highly hydrophobic structure with unique components such as arabinogalactan, a highly branched polysaccharide that connects the peptidoglycan with the outer mycolic acid layer, strongly limiting the permeability to compounds. In addition, cellular components like efflux pumps might also play an important role in the bioavailability of the compounds. Another aspect is the nature of the bacteria proliferation niche. While *L. pneumophila* resides and proliferates in an ER-like vacuole, *M. marinum* starts proliferating in a phagosome-derived vacuole and then continues after escape to the cytosol.

Surprisingly, the two assays used to identify anti-infective or anti-virulence compounds against *M. marinum* did not identify any common hit. As presented in table 2B (1^st^ and 2^nd^ column), hits hallmarks are linked to the intrinsic screen design. Indeed, in the two infection assays, the compounds are added post-infection, and therefore, compounds that hamper bacteria uptake cannot be detected. Thus, both assays detect anti-infective hits that inhibit virulence or boost host defences. On the contrary, in the anti-virulence assay (third column, Table 2), the uptake efficiency of *M. marinum* might be directly affected by compounds that either target the host phagocytosis machinery, or modify the mycobacterial cell wall. In addition, eleven of the fifteen anti-virulence hits identified in the phagocytic plaque assay had selective antibiotic activity against *M. marinum* (Figure 7C).

To better understand the contrasting results of the anti-infective and the anti-virulence screens on *M. marinum*, the hits from both primary assays were re-tested in the *D. discoideum-M. marinum* infection assay (1^st^ column, Table2 and Figure 8A). Satisfyingly, two-thirds of the anti-infective hits originally identified in the *A. castellanii* assay were confirmed using the *D. discoideum* infection assay, likely reflecting the overall conservation of basic metabolic pathways between these two evolutionarily close organisms. In contrast, only half of the anti-virulence hits showed mild but significant activity in the *D. discoideum* infection assay (figure 8B). One possibility is the presence of an MCV membrane around *M. marinum* that restricts the bioavailability of the compounds. In addition, it is known that *M. marinum* undergoes drastic metabolic adaptations to different carbon sources when transitioning between its extracellular life to the intracellular environment or, possibly explaining the differential sensitivity to the anti-virulence hits.

### Pro-infective, another facet to better understand the *Mycobacterium-host* interactions

On the other hand, identification of infection enhancers in the set of compounds was quite surprising, although the same observation was already reported in the phenotypic screen of the GlaxoSmithKlyne TB-set of anti-mycobacterial antibiotics in the A. *castellanii-M. marinum* model of infection (Trofimov et al. 2018). Notably here, five compounds were identified that increased at least 2-fold the intracellular *M. marinum* bacterial load (Fig. 5A). As discussed in the Trofimov *et al*. paper, such compounds may be targeting and disarming crucial anti-bacterial host defence pathways and therefore, might lead to a better understanding of these pathways and might ultimately lead to the design of host-directed anti-mycobacterial therapies.

### Confirmation of the low cytotoxicity of compounds on *D. discoideum*

Measurement of the hit compounds’ cytotoxicity/growth inhibition activity was performed by monitoring the growth of GFP-ABD *D. discoideum*. Four out of six compounds that are either toxic for *D. discoideum* or affect its growth are common between two screens (figure 6A, B and C). This low number of toxic compounds might be explained by the fact that the pathways selected for the LVS are non-essential for host metabolism and survival. In addition, the LVS likely enriched the library for compounds with drug-like properties that are anticipated to have low toxicity. The anti-infective hits with mild cell growth inhibition activity should be further investigated to optimize their therapeutic window.

### Combination of 3R model assays increases the chances of identifying potential anti-Tb compounds

In conclusion, our data show that the ligand-based virtual screening efficiently selected anti-bacterials from the 2.5 Mio lead-like compounds of the ZINC library, giving rise to hit rates 2 to 3 times superior to the 1% usually observed by random screening. The data also demonstrate that the validated virtual hits are chemically diverse, suggesting that they most likely target different pathways within the host pathogen system. We suggest that our combination of cost-effective, 3R compliant amoebae-based phenotypic assays to screen structurally diverse chemical libraries efficiently identifies a variety of promising non-toxic anti-infective compounds that then will be validated in more complex infection systems such as the zebrafish or mouse models.

## Materials and methods

### Design and characterization of the chemically highly diverse pathway-based library for phenotypic screening

In recent years, the design of diverse libraries based on the principle of functional diversity has become a major trend in library design (Shelat and Guy 2007). This includes designing libraries containing privileged structures (REF) as well as diverse scaffolds to best cover the chemical space. In this work, we selected in total 18 host and pathogen pathways (Table 1) as potential pharmacological targets to develop pharmacophores queries based on ligands/metabolites known to interfere/intervene within these host and pathogen pathways (Table 1). These pharmacophore queries have been used in ligand-based virtual screening of the ZINC lead-like database (http://zinc.docking.org/) (Sterling and Irwin 2015) using ROCS, a tool from the OpenEye software package (http://eyesopen.com/rocs) (Swann et al. 2011). The ligand-based VS was performed with ROCS using previously published default settings (Kirchmair et al. 2009) and the TanimotoCombo score. To ensure chemical diversity and maximize the coverage of the chemical space of the ZINC lead-like database we applied the Lingo method program (Vidal, Thormann, and Pons 2005). We screened the ZINC lead-like database composed of 2.5 million compounds and finally selected 100 compounds per host and pathogen pathways for a total of 1’800 compounds using the workflow described in Figure 1. Thus, the selected virtual hits correspond to 0.07% of the whole ZINC lead-like database. The number of virtual hits was tractable by the medium throughput assays described in this work.

The pathways-based library properties were characterized in term of chemical diversity and drug likeness. To assess chemical diversity, we used the Z-matrix calculated according to Tanimoto’s chemical similarity metrics (*T_sim_*) using Canvas, a tool from the Schrödinger software package (Sindhikara and Borrelli 2018). The results have been displayed as heat map drawn using Netwalker1.0 (Komurov et al. 2012). The drug-likeliness of compounds of the library was assessed by predicting physico-chemical descriptors using Canvas [Ghosh et al., J Comb Chem 1999, 1(1): 55–68] and comparing them with the different known rules in drug discovery (Lipinski et al. 1997; Lipinski and Hopkins 2004; Lipinski 2016).

### Bacterial and cell cultures

*A. castellanii* (ATCC 30234) was grown in PYG medium at 25°C as described (Moffat and Tompkins, 1992; Segal and Shuman, 1999) using proteose peptone (Becton Dickinson Biosciences) and yeast extract (Difco). The *D. discoideum* strain was grown in HL5c medium at 22°C.

*L. pneumophila* were re-suspended from plates in appropriate growth medium, ACES Yeast Extract (AYE) or Luria Broth (LB), and diluted to a starting OD600 of 0.01. Compounds were added to these cultures such that the maximal DMSO concentration was 0.1%. Cultures were grown overnight and the OD600 measured.

*M. marinum* were cultured in Middlebrook 7H9 (Difco) supplemented with 10% OADC (Becton Dickinson), 5% glycerol and 0.2% Tween80 (Sigma Aldrich) at 32°C in shaking culture. The *M. marinum* strain constitutively expressing GFP was obtained by transformation with msp12::GFP, and cultivated in the presence of 20 μg/ml kanamycin.

### Intracellular replication of *M. marinum* in *A. castellanii*

*A. castellanii* were cultured in PYG medium in 10 cm Petri dishes at 25°C, and passaged the day prior to infection to reach 90% confluence. *M. marinum* were cultivated in a shaking culture at 32°C to an OD600 of 0.8-1 in 7H9 medium. Mycobacteria were centrifuged at RT at 500 g for two periods of 10 min onto a monolayer of *Acanthamoeba* cells at an MOI of 10 to promote efficient and synchronous uptake, followed by an additional 20-30 min incubation. Un-ingested bacteria were washed off with PYG and infected cells re-suspended in PYG containing 10 μM amikacin. 5 × 10^4^ infected cells were transferred to each well of a 96-well plate (Cell Carrier, black, transparent bottom from Perkin-Elmer) with pre-plated compounds and controls The course of infection at 25°C was monitored by measuring fluorescence in a plate reader (Synergy H1, BioTek) for 72 hours with time points taken every 3 hours. Only experiments with a Z-factor > 0.6 (calculated from DMSO and rifabutin controls) were taken into account for analysis. Time courses were plotted and data from all time points (using cumulative curves) were used to determine the effect of compounds versus vehicle controls. The primary hit rate cut off was set at 20% inhibition for *M. marinum*.

### Intracellular replication of *M. marinum* in *D. discoideum*

*D. discoideum* were cultured in HL5c medium in 10 cm Petri dishes at 25°C, and passaged the day prior to infection to reach 90% confluency Mycobacteria were grown in 7H9 medium to a density of OD600=0.8-1.0 (5×10^8^ bacteria ml^−1^), centrifuged and re-suspended in HL5c medium and clumps disrupted by passaging through a 26-gauge needle. GFP-expressing *M. marinum* were added at an MOI of 10 and centrifuged onto the *Dictyostelium* cells at 500 *g* twice for 10 min. The cells were left at 25°C for an additional 10–20 min before uningested bacteria were washed off by 3 washes with HL5c and Attached cells were then re-suspended in HL5c containing 10 μM amikacin. The course of infection was monitored as described above.

### Intracellular replication of *L. pneumophila* in *A. castellanii*

*A. castellanii* were cultured in PYG medium at 25°C, and passaged the day prior to infection such that 2 × 10^4^ cells were present in each well of a 96-well plate (Cell Carrier, black, transparent bottom from PerkinElmer). Cultures of *L. pneumophila* were re-suspended from plate to a starting OD600 of 0.1 in AYE medium, and grown overnight in shaking conditions at 37°C to an OD_600_ of 3. Re-suspended bacteria in LoFlo medium (ForMedium) were centrifuged onto a monolayer of *A. castellanii* cells at an MOI of 20 to promote efficient and synchronous uptake. Compounds were added to at least triplicate wells after infection, and infected cells were incubated at 30°C. GFP fluorescence was measured by a plate spectrophotometer at appropriate intervals (Optima FluoStar, BMG Labtech). Because the culture media used for *A. castellanii* do not support the growth of *L. pneumophila*, GFP fluorescence accurately reflects intracellular replication. The hit rate cut off was set at 40% inhibition for *L. pneumophila*. Time courses were constructed and data was used to determine the effect of compounds versus vehicle control.

### Anti-virulence assay against *M. marinum*

To test the effect of the compounds on M. *marinum* virulence, 10 ml of mid-log phase mycobacterial cultures (OD600 around 0.8–1.2) were pelleted by centrifugation and re-suspended in 5 ml of an overnight culture of K. *pneumoniae* diluted to 10^−5^ in LB medium (Alibaud et al. 2011). The mixture was de-clumped by passaging through a 25-gauge blunt needle. In each well of a 24-well plate, 10 μM of each compound was added and allowed to diffuse on 2 ml of solid standard medium (SM) agar supplemented with glucose followed by the addition of 50 μL of the bacterial suspension Once dried, 1,000 *D. discoideum* cells were added in the centre of the well. Plates were incubated for 5–9 days at 25°C and the formation of phagocytic plaques was monitored visually, a negative control (Bacteria + *D. discoideum* + DMSO) was included in every plate.

### Antibiotic activity assays

Antibiograms used to monitor the inhibitory effect of compounds on mycobacterial growth were performed as described previously (Ouertatani-Sakouhi et al. 2017), each molecule was added in a 24-well plate well containing 2 ml of 7H11 agar medium at 10 μM. Once dried, 1,000 bacteria were deposited in each well and plates were incubated at 32°C for 7 days to allow bacterial growth.

To monitor *M. marinum* growth, GFP-expressing bacteria were cultivated in shaking at 32°C in 7H9 medium supplemented with OADC up to an OD600 of 0.8–1. 10^5^ GFP-expressing *M. marinum* were transferred into each well of 96-well white plates. To monitor *L. pneumophila* growth, a pre-culture of GFP-expressing bacteria were diluted to a starting OD600 of 0.01 and grown overnight. Compounds at 30 μM in DMSO were added in each well, seeded with 100 μl of pre-culture, and bacterial growth was monitored for at least 48 hours by measuring the fluorescence in a plate reader (Synergy H1) every 3 hours.

### *D. discoideum* growth inhibition assay

10^4^ GFP-ABD-expressing *Dictyostelium* cells were transferred to each well of 96-well plates allowed to attach for 20-30 min. Cell growth at 25°C was monitored by measuring the GFP fluorescence in a fluorescent plate reader (Synergy H1, company) for 72 hours with the time point every 3 hours.

### Cytotoxicity Assay

Cytotoxicity of compounds against *A. castellanii* was determined using the Alamar Blue reagent (Life Technologies). To mimic the conditions found in the intracellular replication assay, 96-well plates were set up as previously described and uninfected triplicate wells were treated with compounds in 100 μl LoFlo media (Harrison et al. 2013). Plates were incubated at 30°C for 24 h, after which 10 μl Alamar Blue reagent was added, and plates incubated for a further 3 h. The fluorescence at 595 nm was measured, and data normalized between 1 (treatment with LoFlo alone) and 0 (SDS, total lysis of the cells). Means from each individual experiment were then combined for analysis.

### Data analysis

Data analysis was performed using Microsoft Excel and GraphPad Prism 5. To compare the effect of compound treatment on intracellular replication, fluorescence values were taken from the first time point following entry to stationary phase. The results were then normalised such that media-only wells (no bacteria) were ‘0’, while vehicle-treated wells were ‘1’ (normal replication). The average of the replicate wells (minimum 3 per plate) was then plotted as dose-response curves, such that each individual point represented the average of a single experiment. Compound treatments were repeated a minimum of 3 times to control for the increased variability of bacteria-host cell interactions.

## Acknowledgements

Acknowledgements and funding statement

We gratefully acknowledge D. Moreau from the Bioimaging Center for Microscopy and Image Analysis at the Faculty of Sciences in Geneva for his precious help. This work was supported by Swiss National Science Foundation (SNF) ‘Sinergia’ grant CRSI33_130016 (awarded to L.S., T.S., P.C. and H.H.), an RTD grant from SystemsX.ch (awarded to TS, PC, HH). TS is a member of iGE3 (http://www.ige3.unige.ch). The funders had no role in study design, data collection and analysis, decision to publish, or preparation of the manuscript.

---

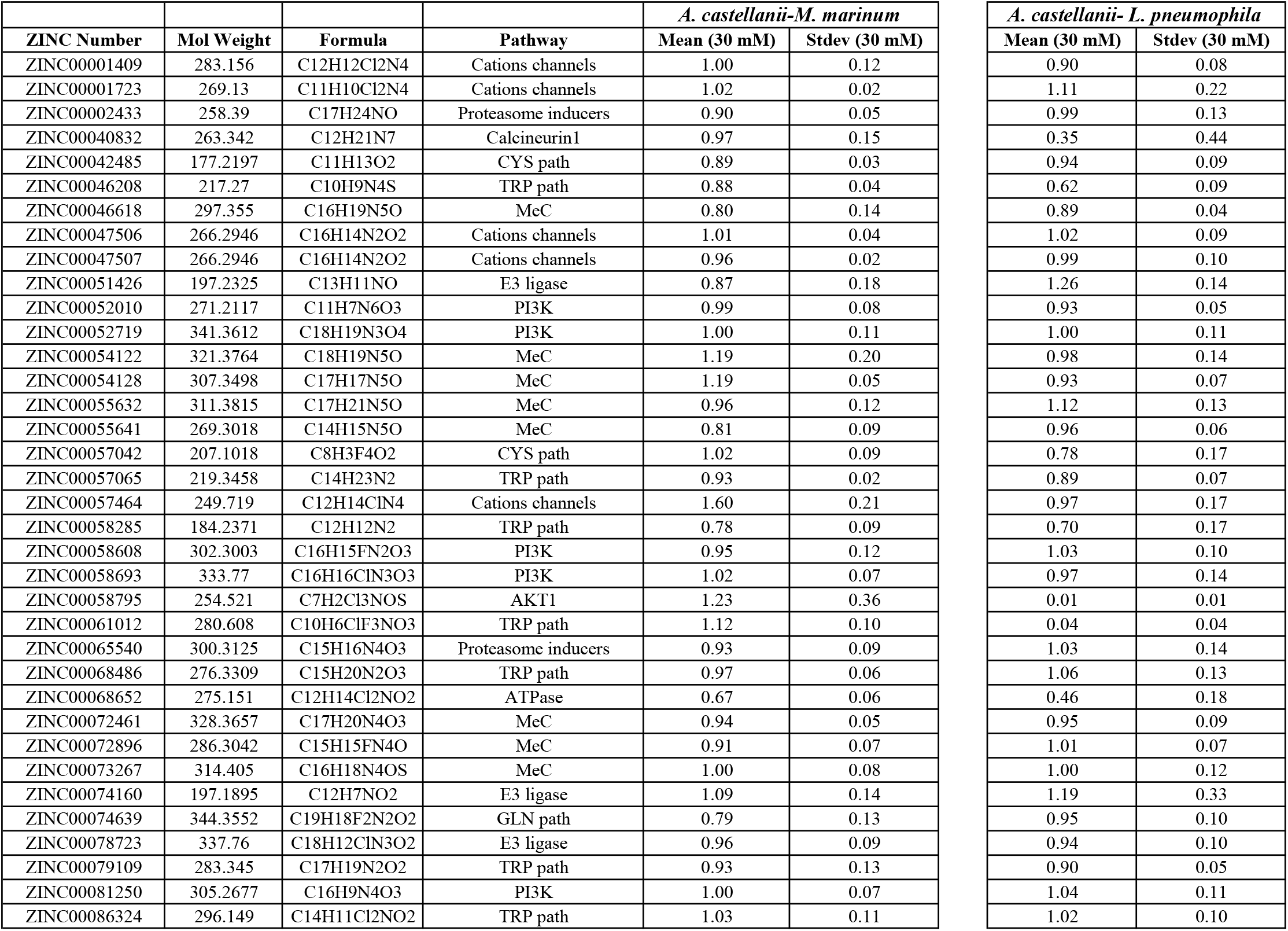
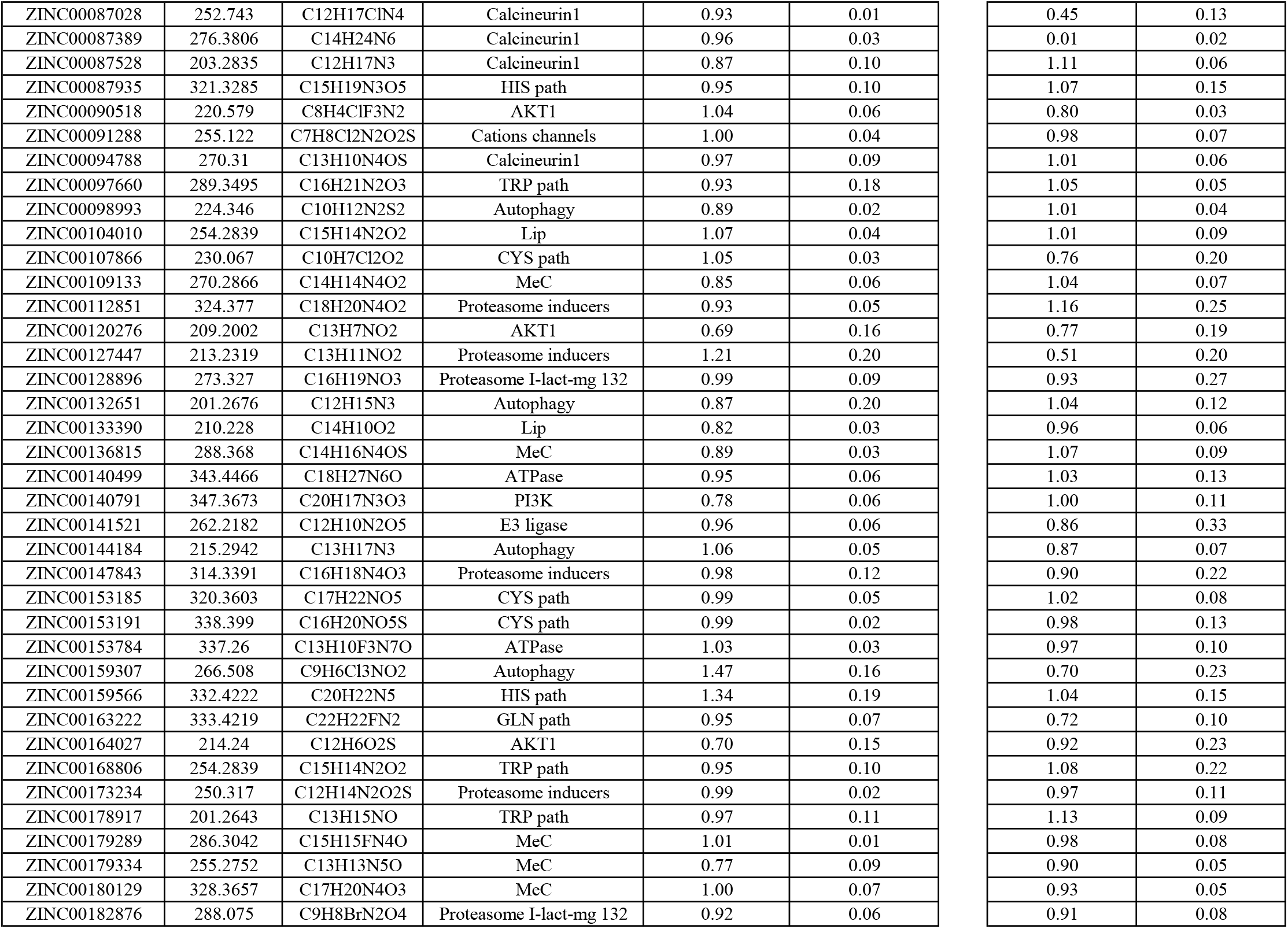
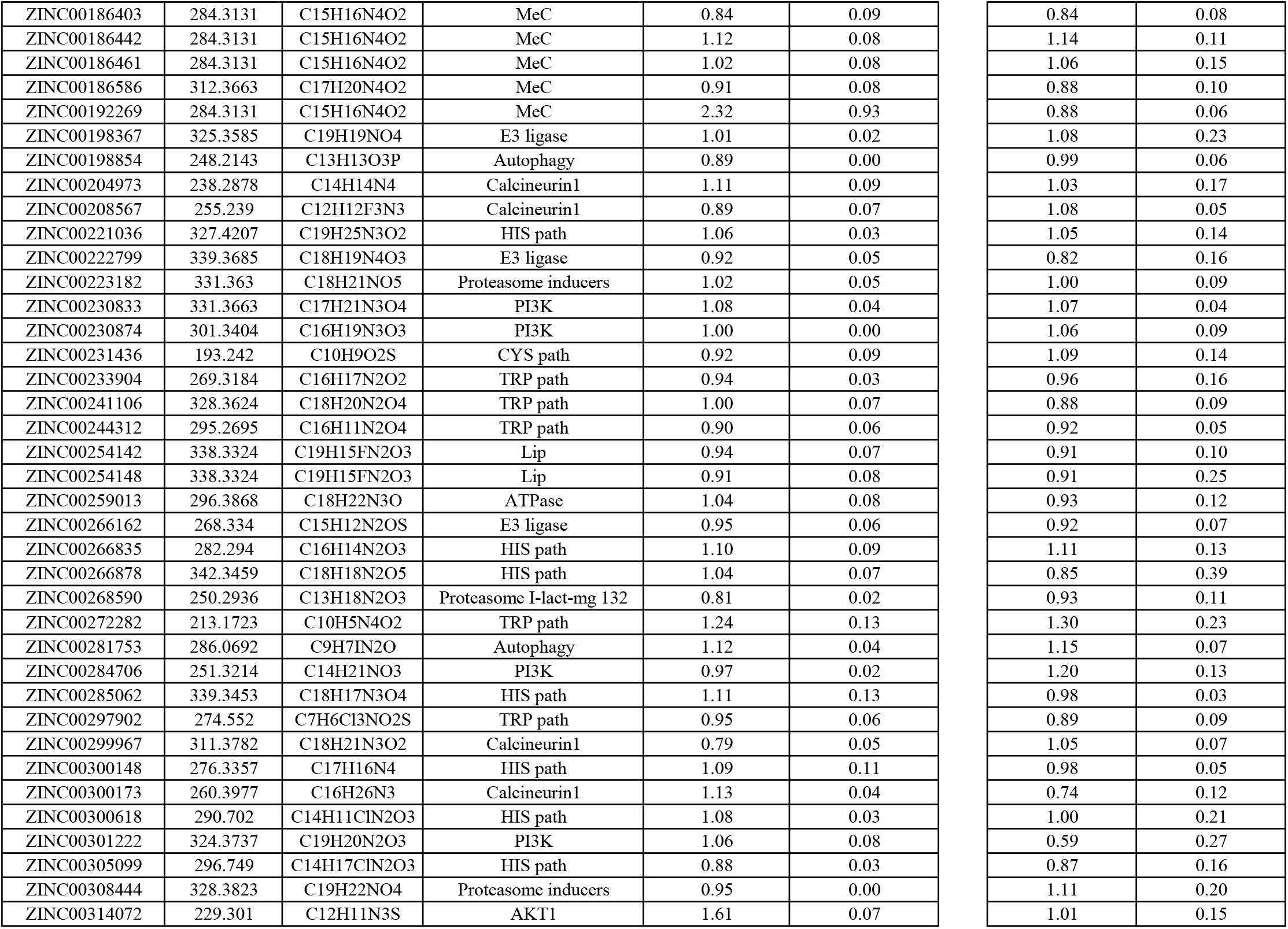
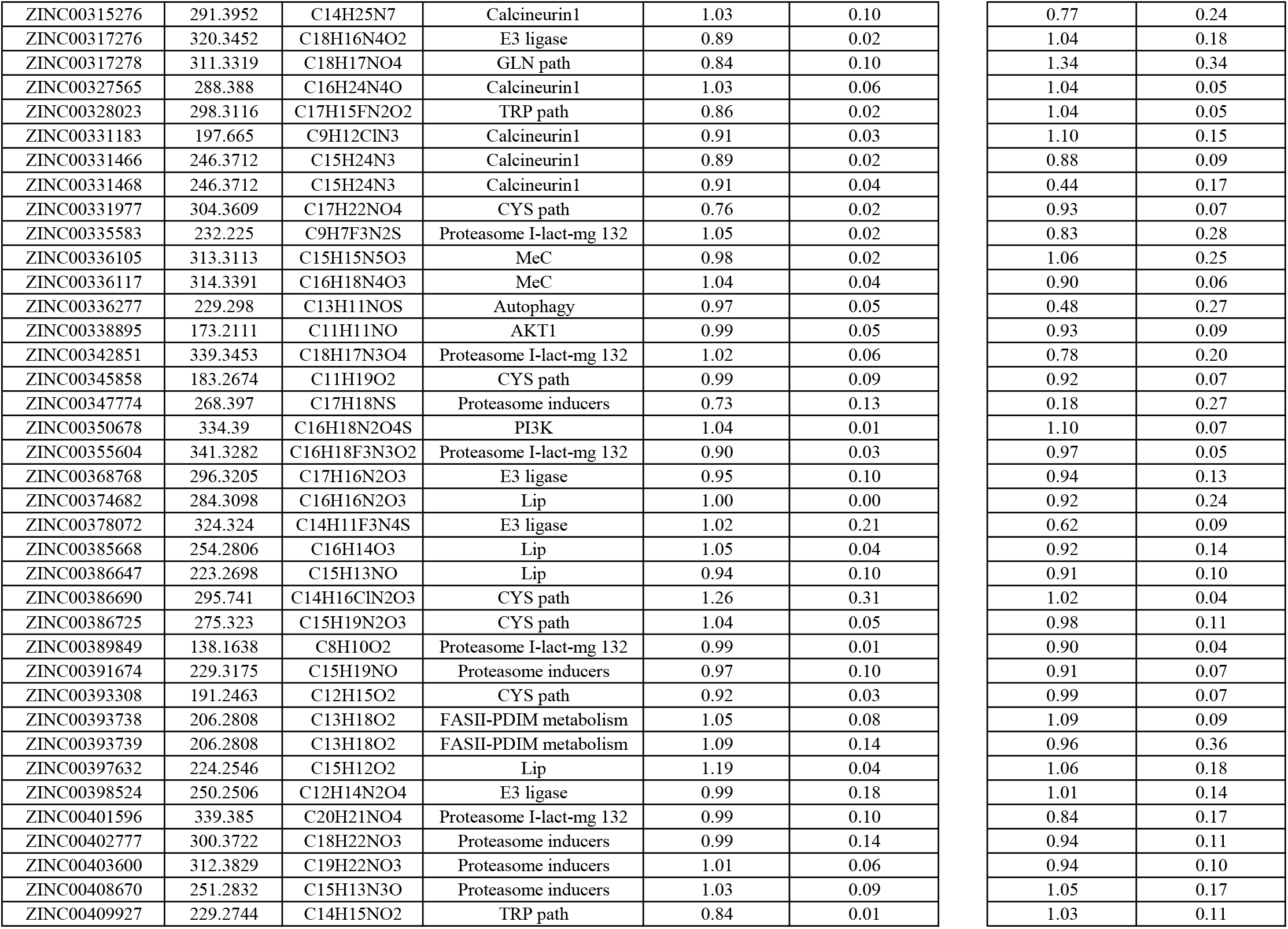
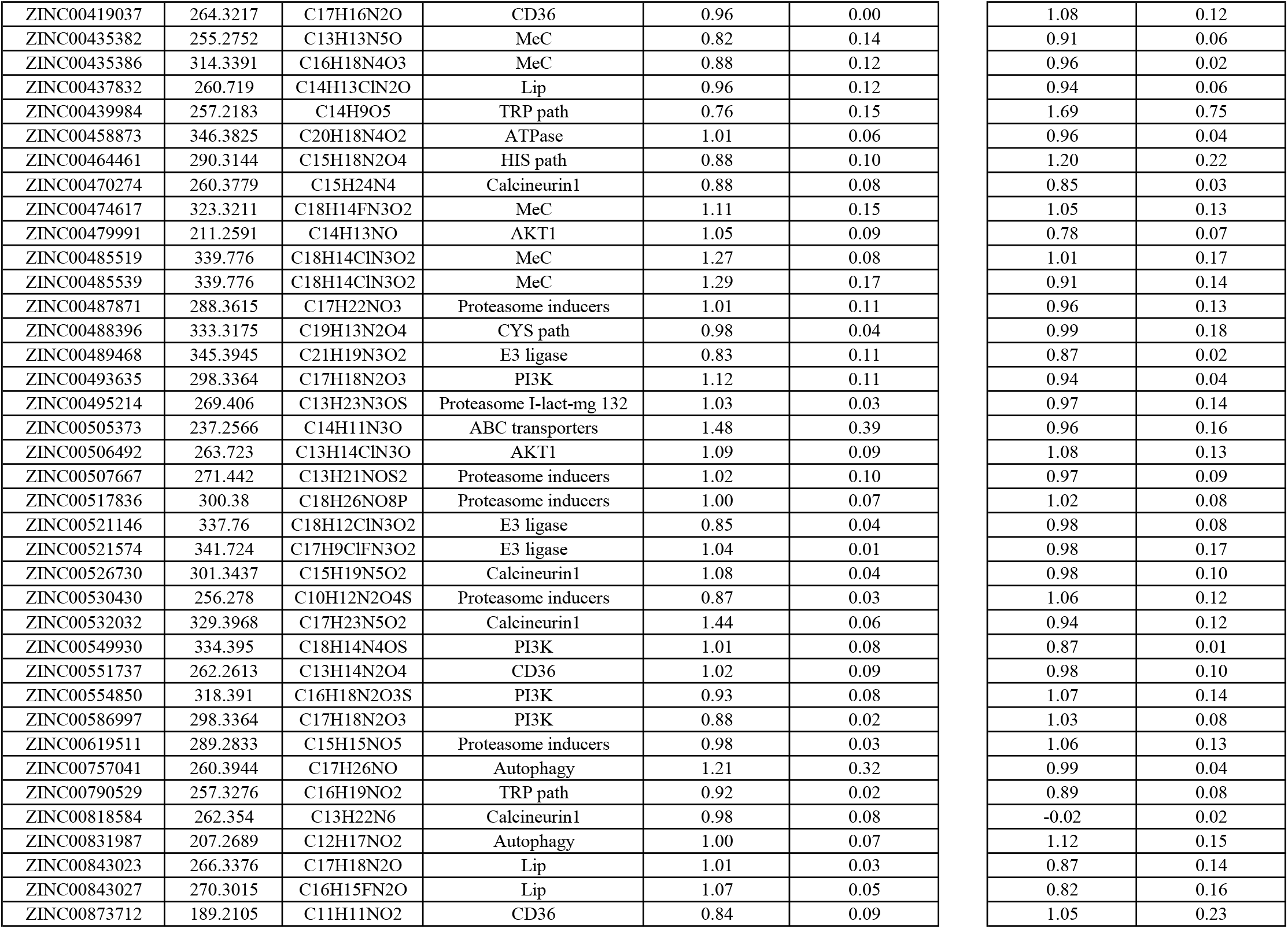
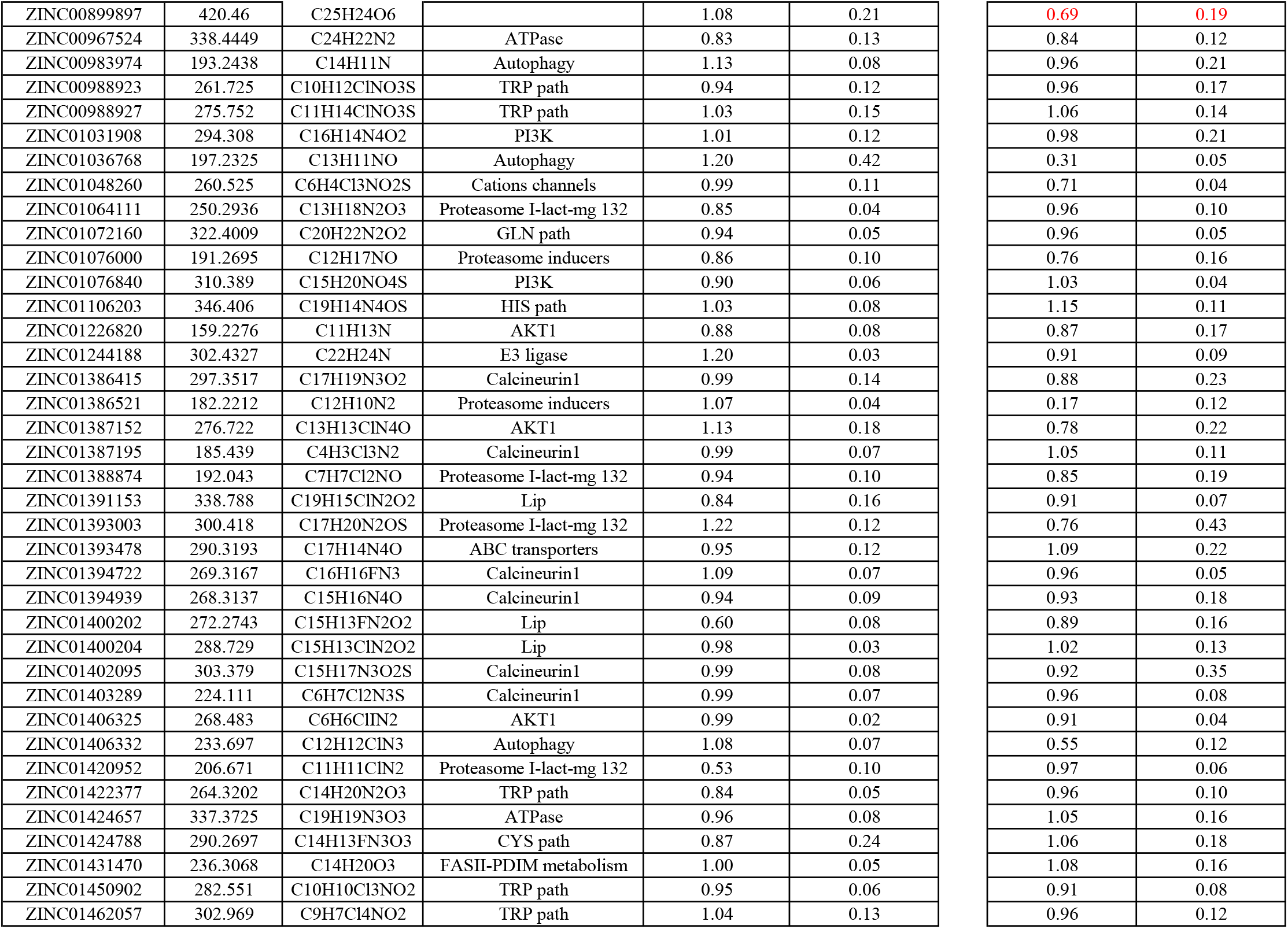
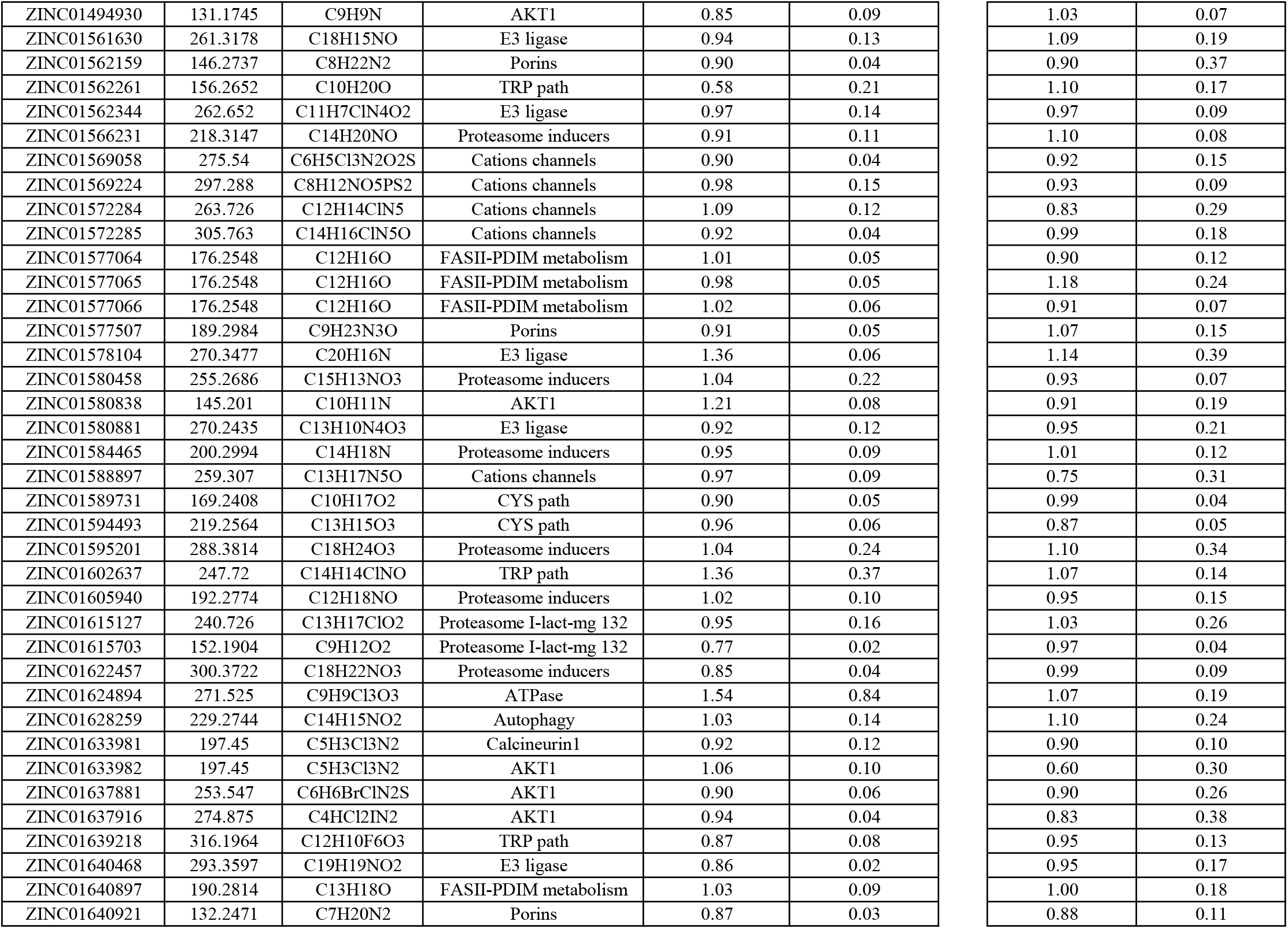
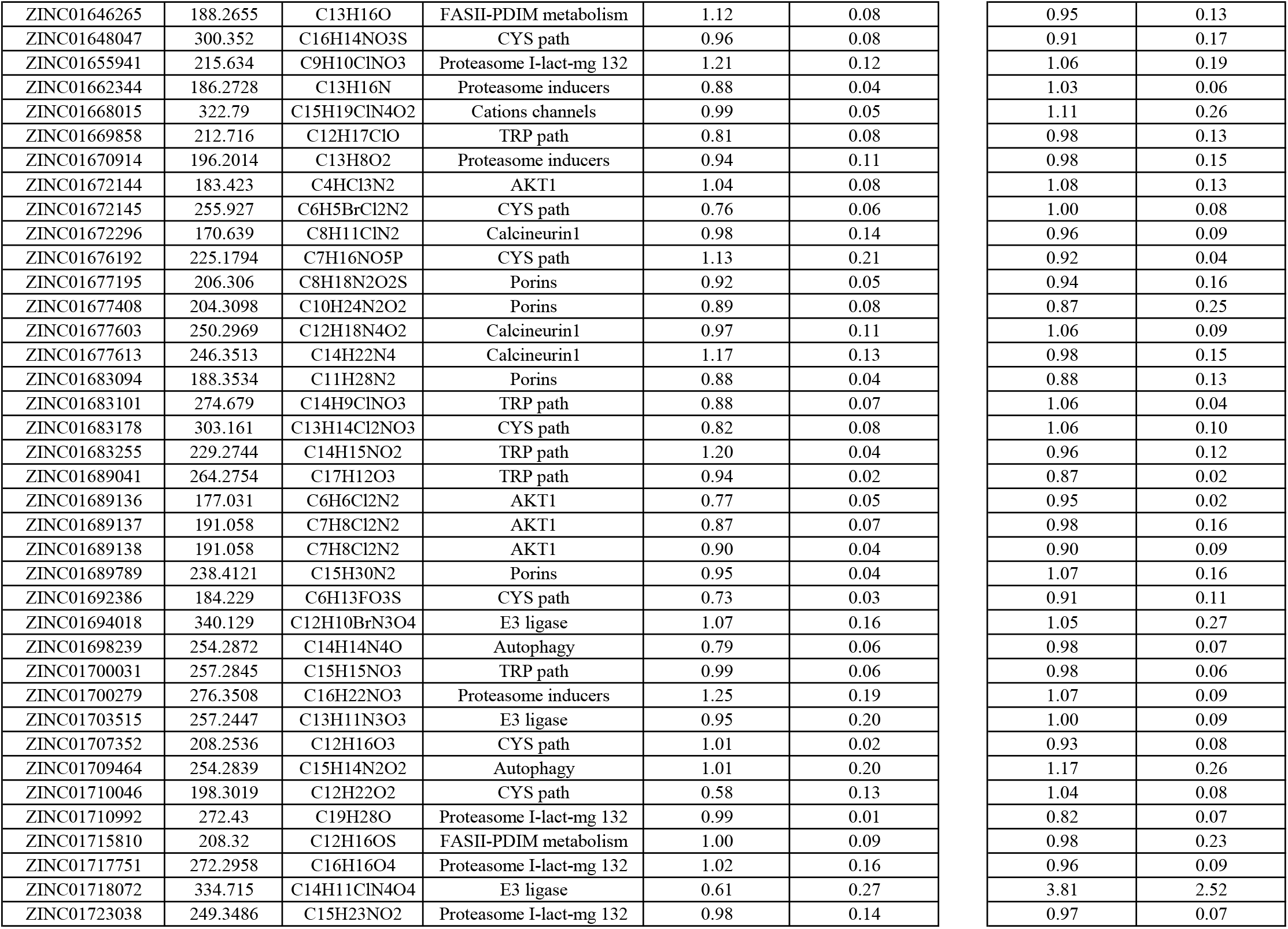
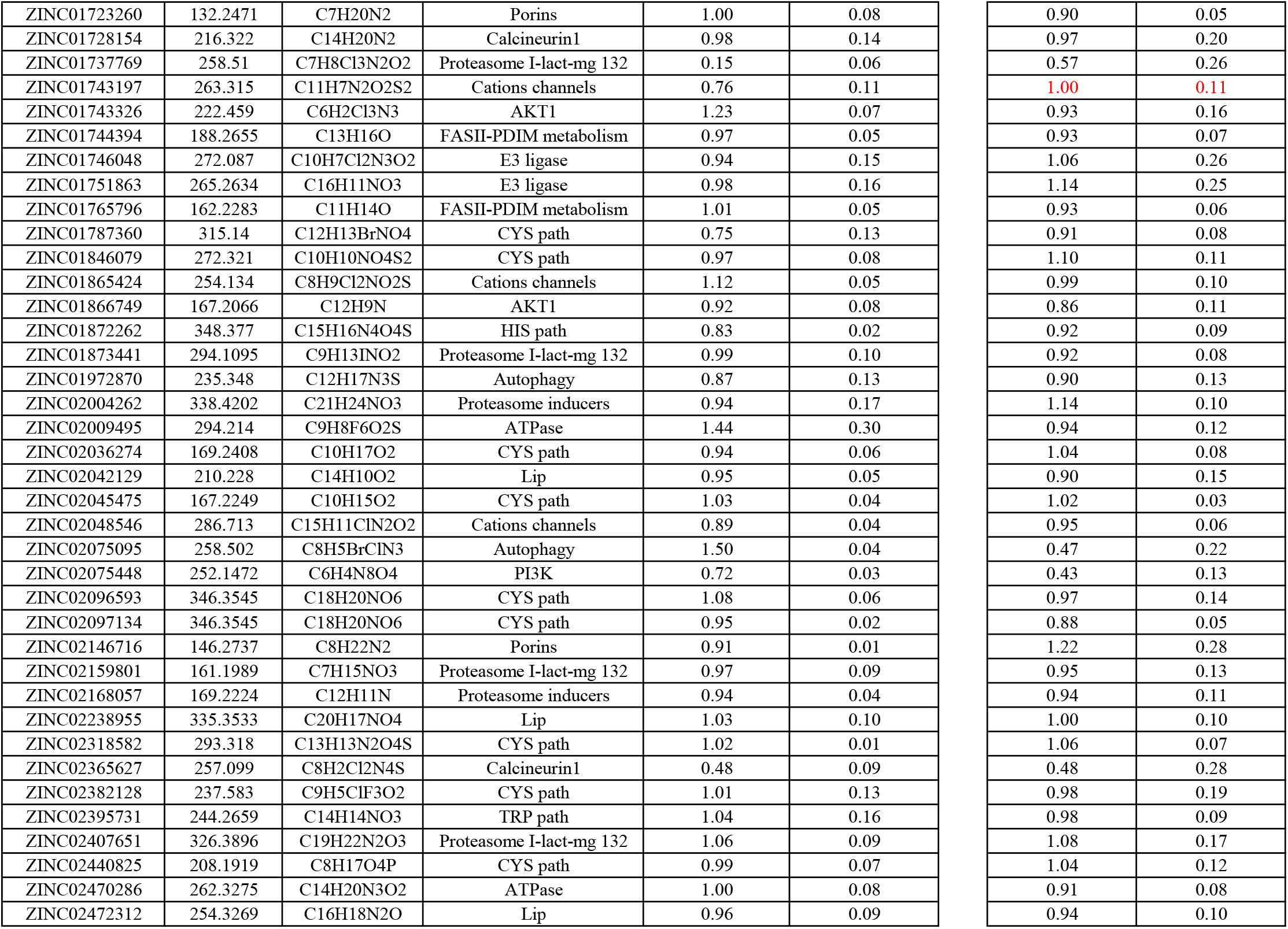
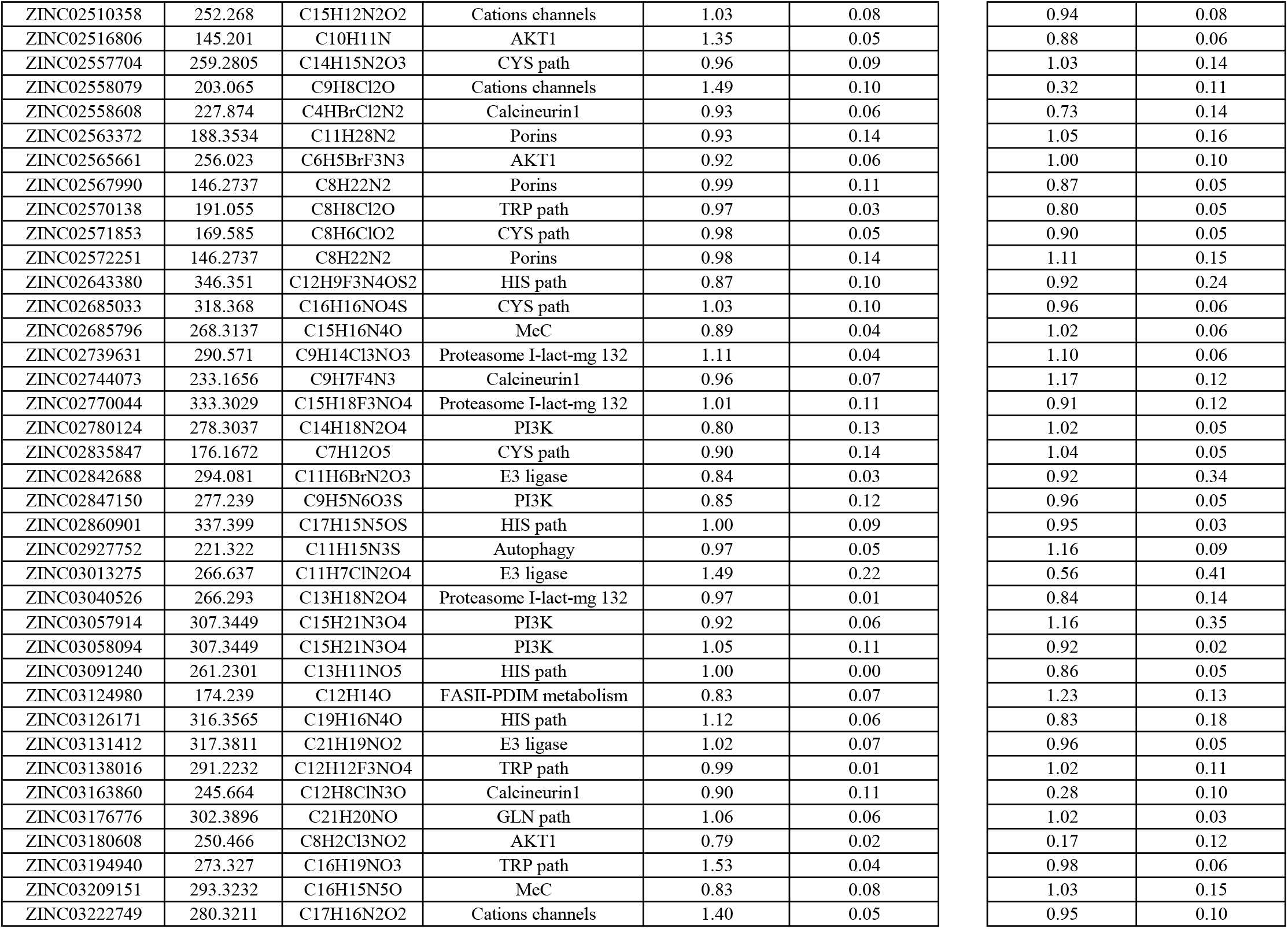
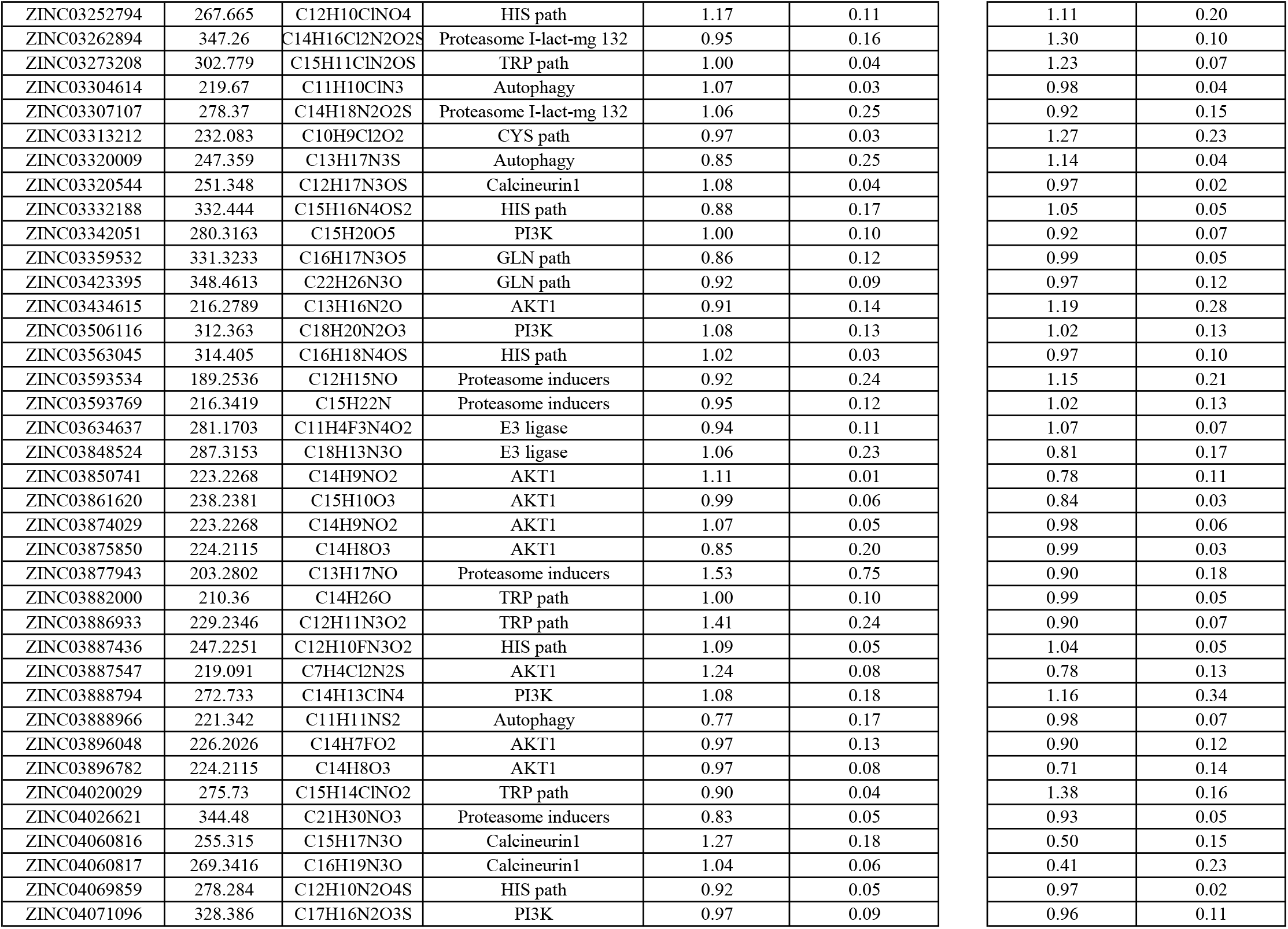
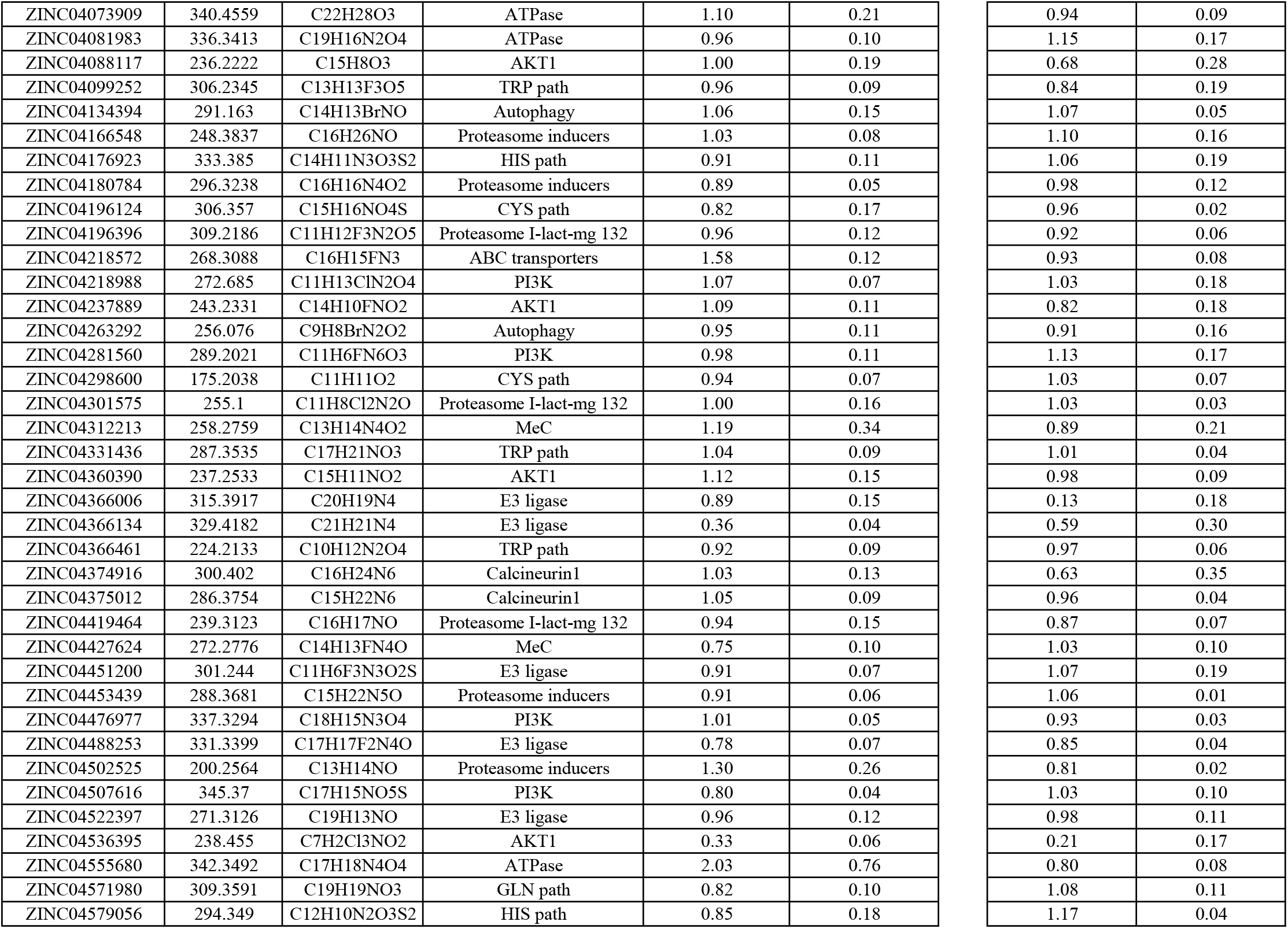
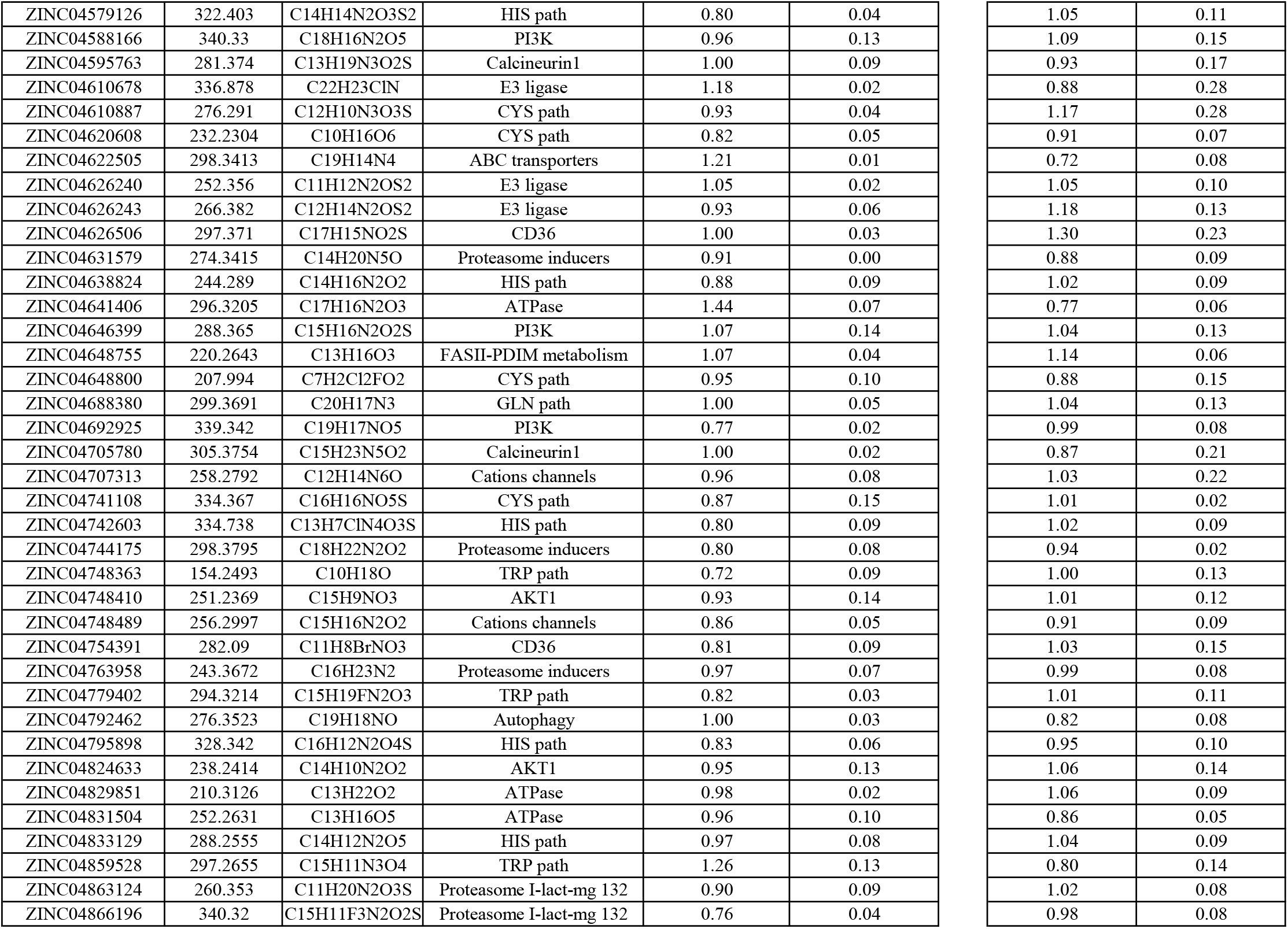
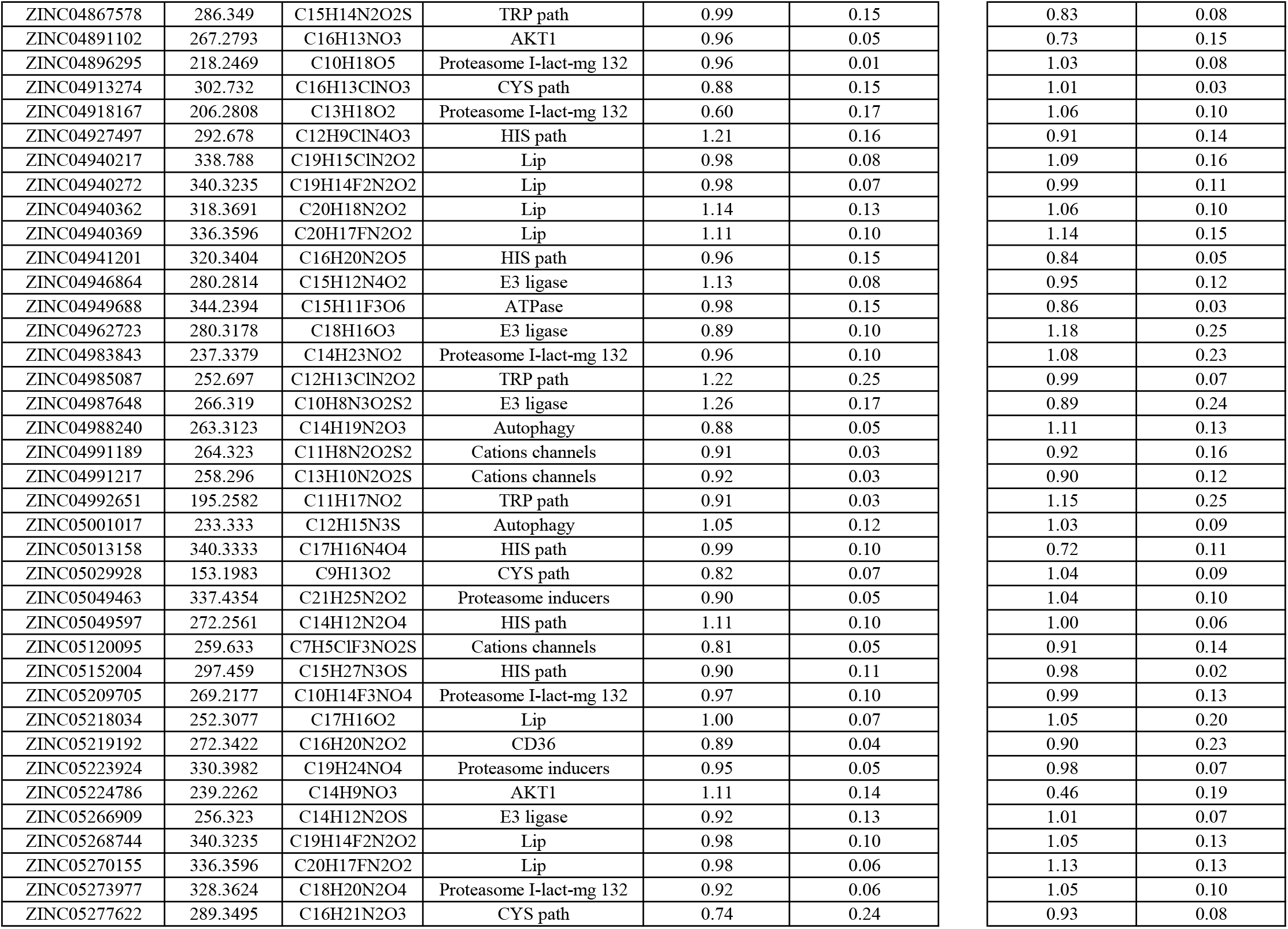
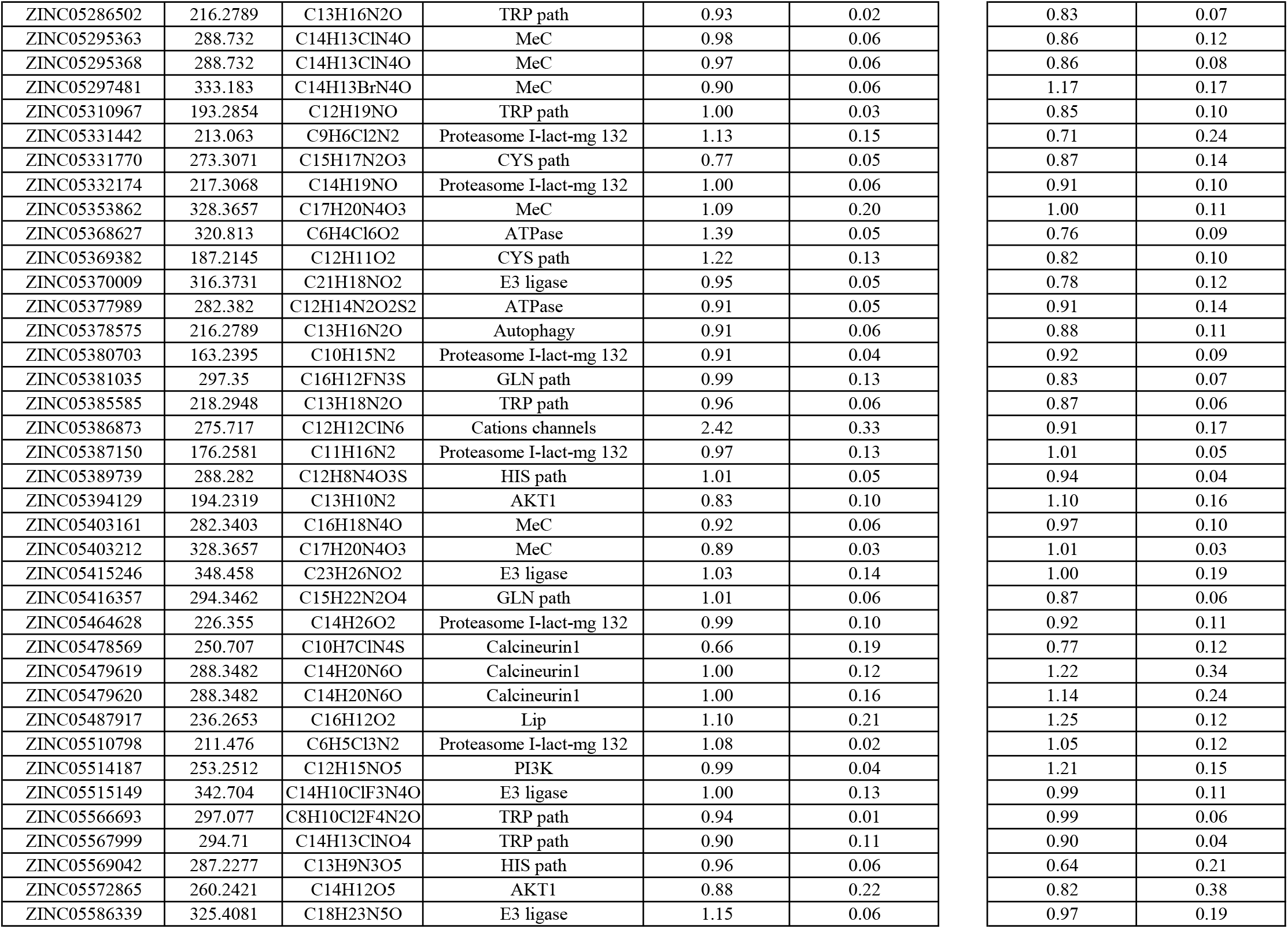
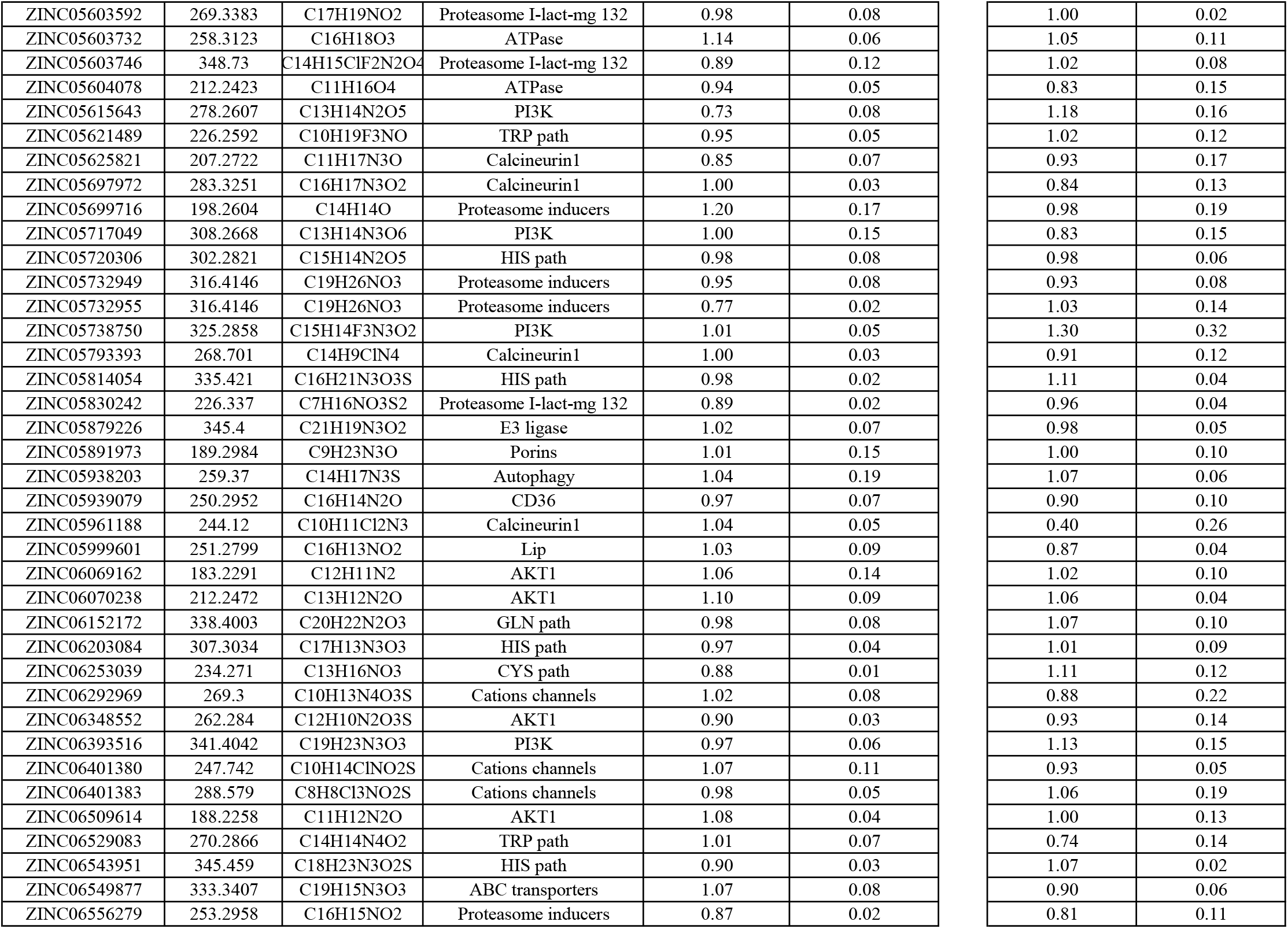
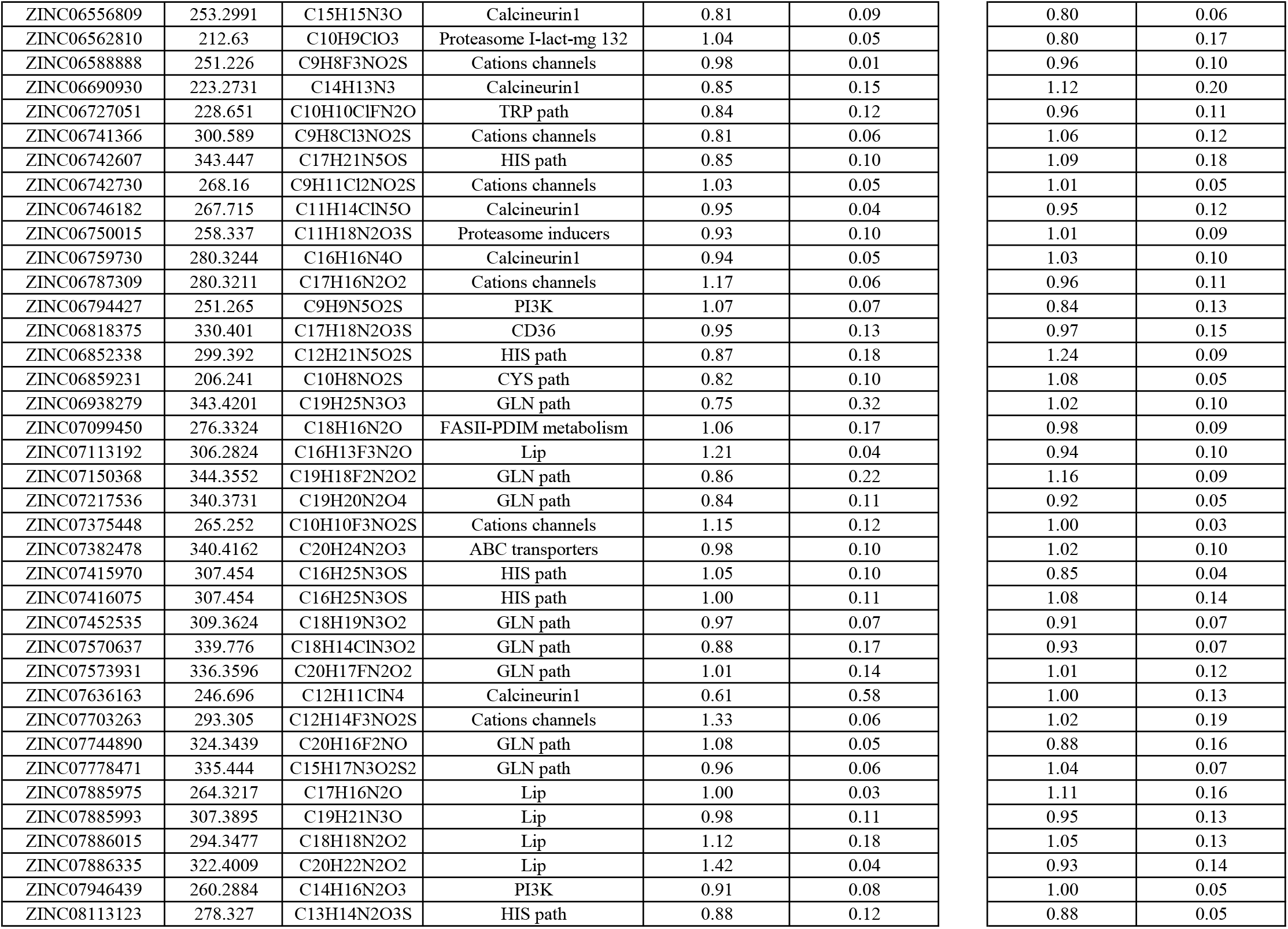
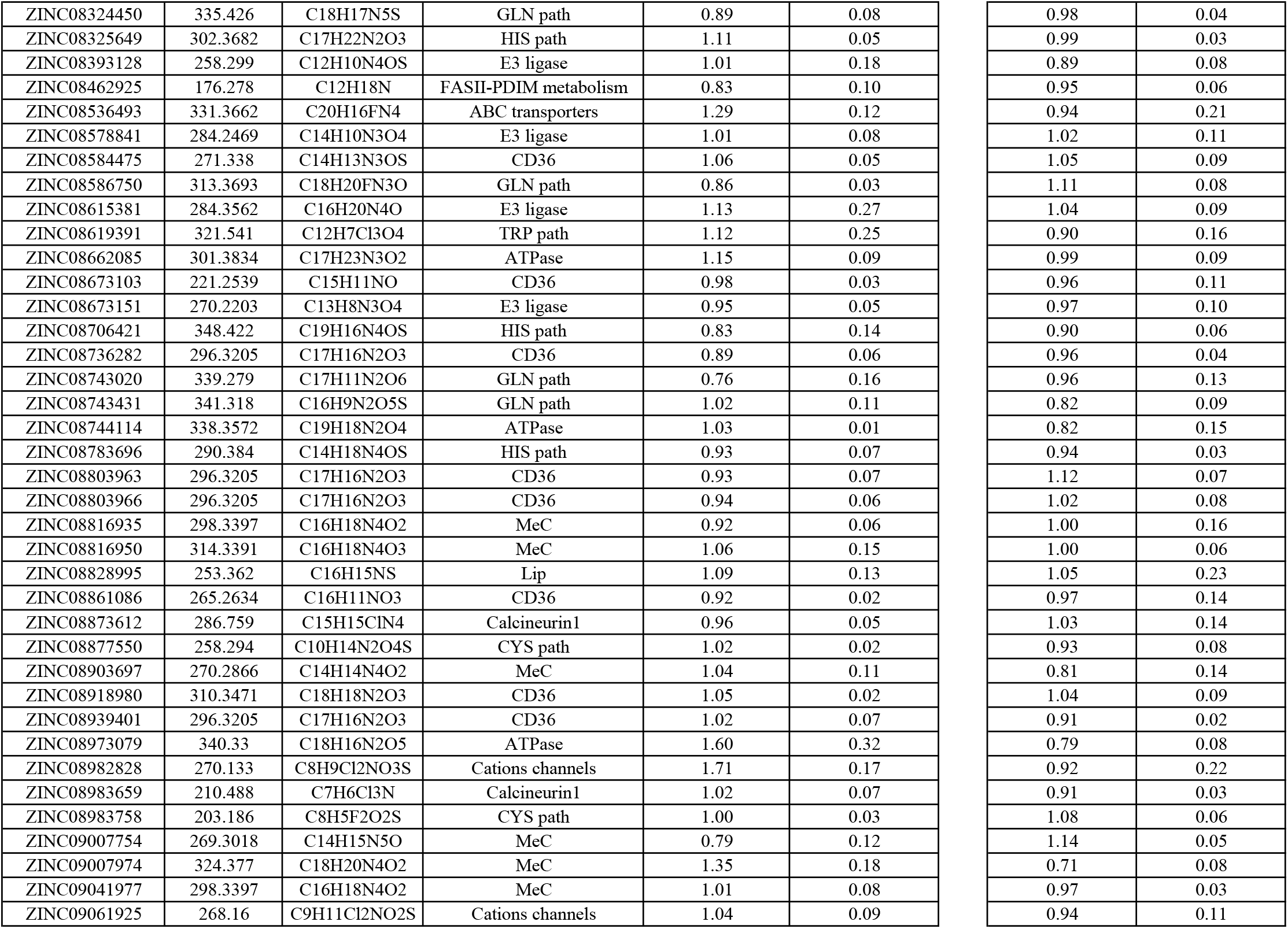
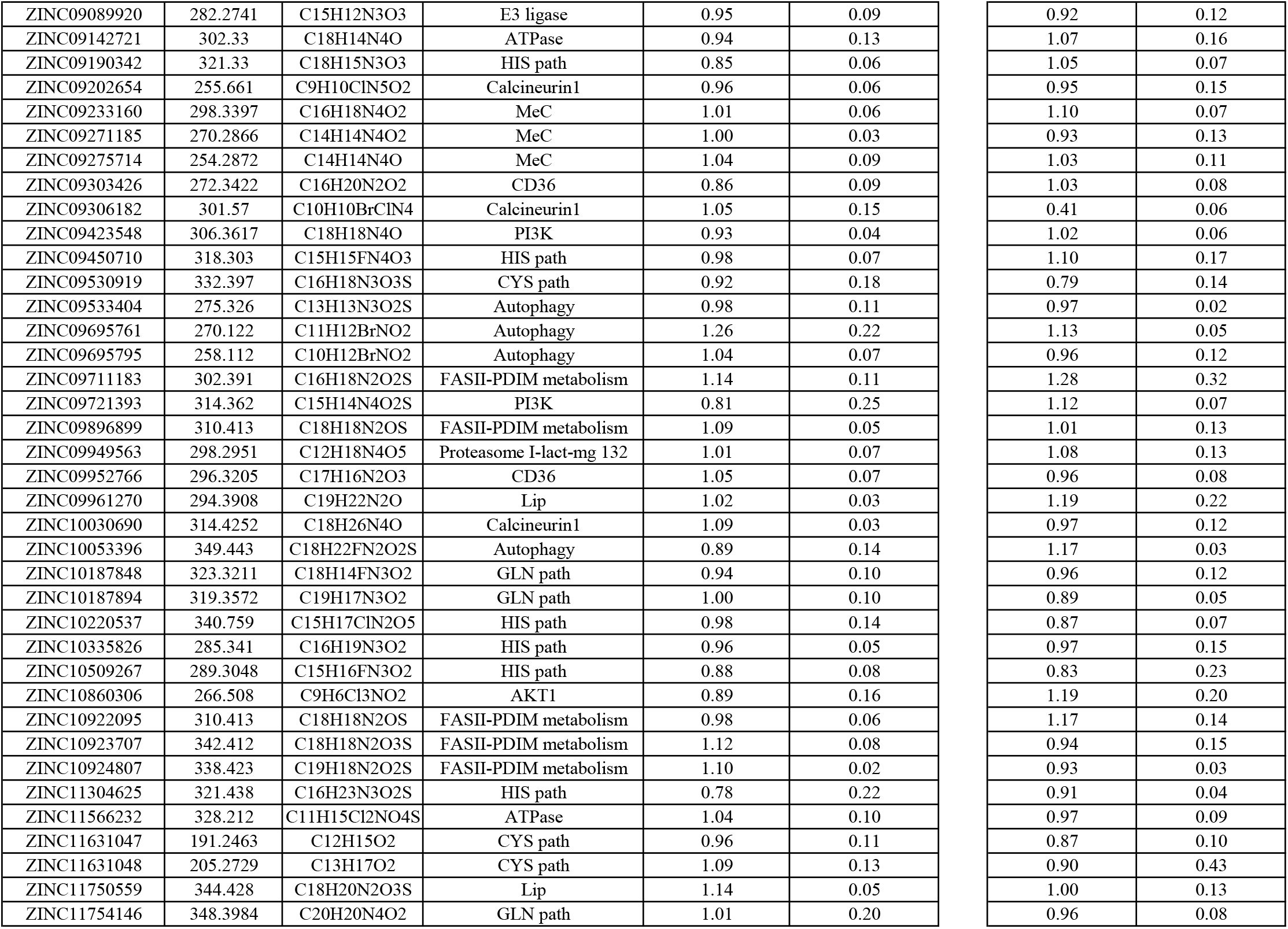
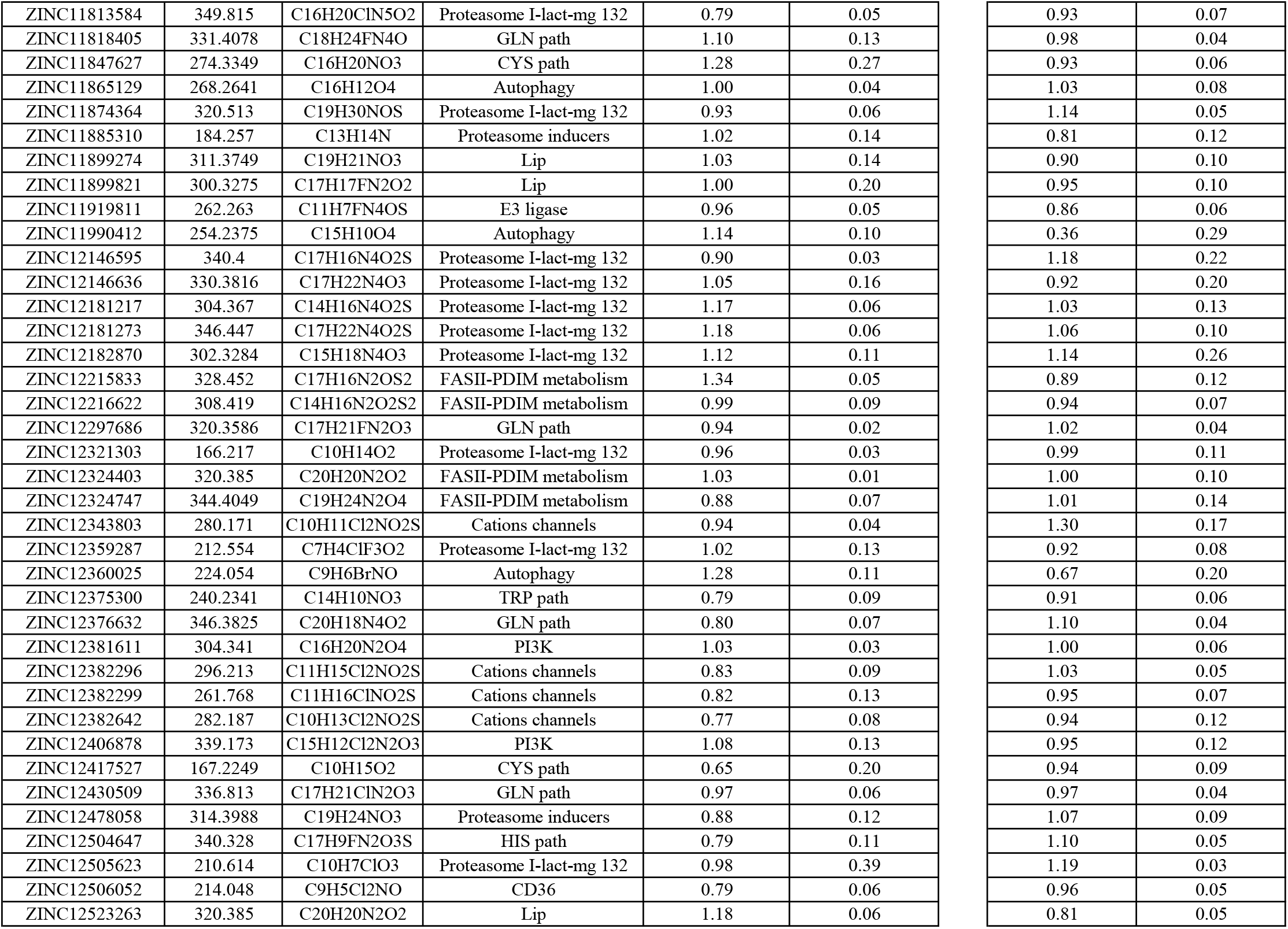
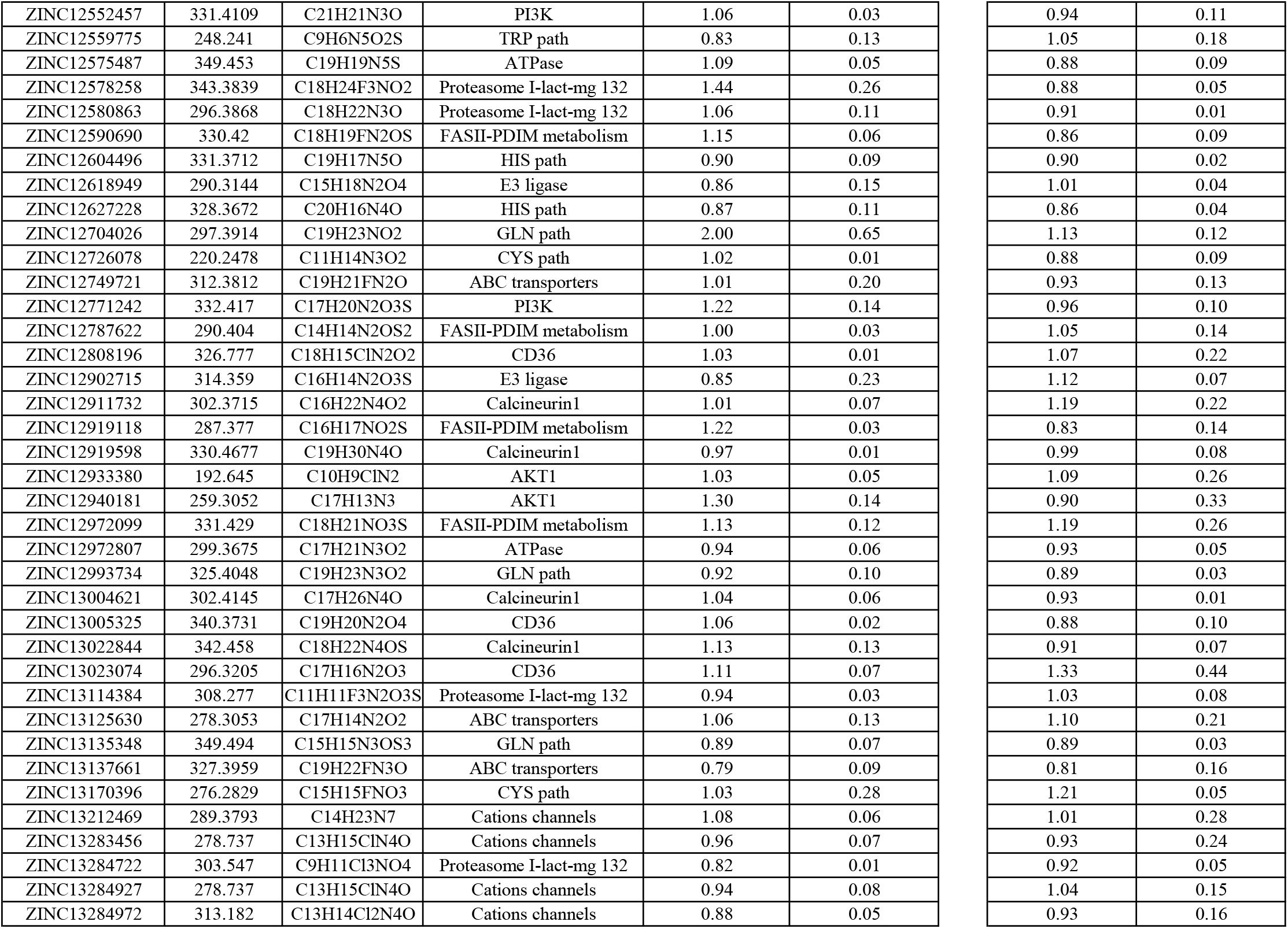
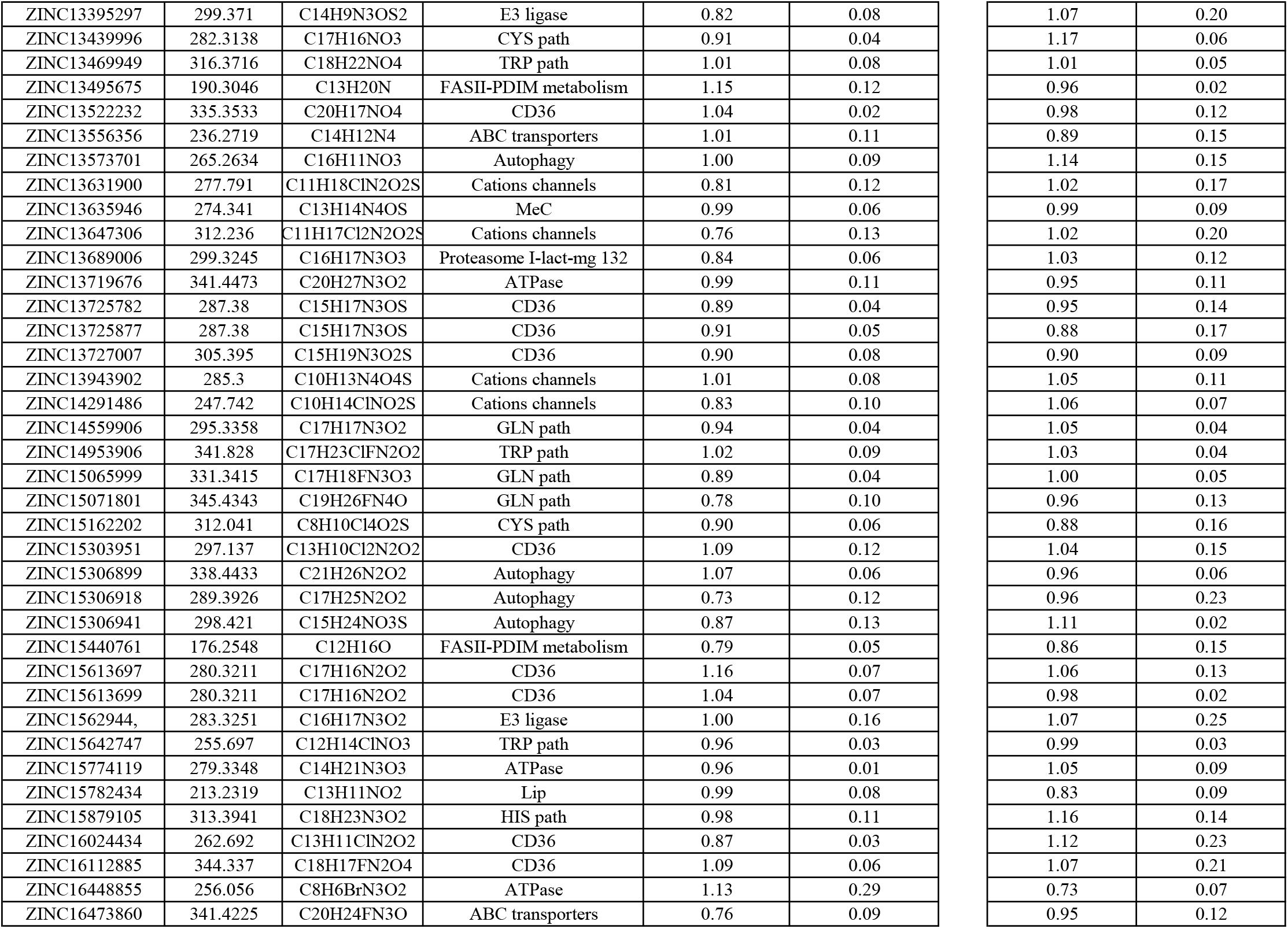
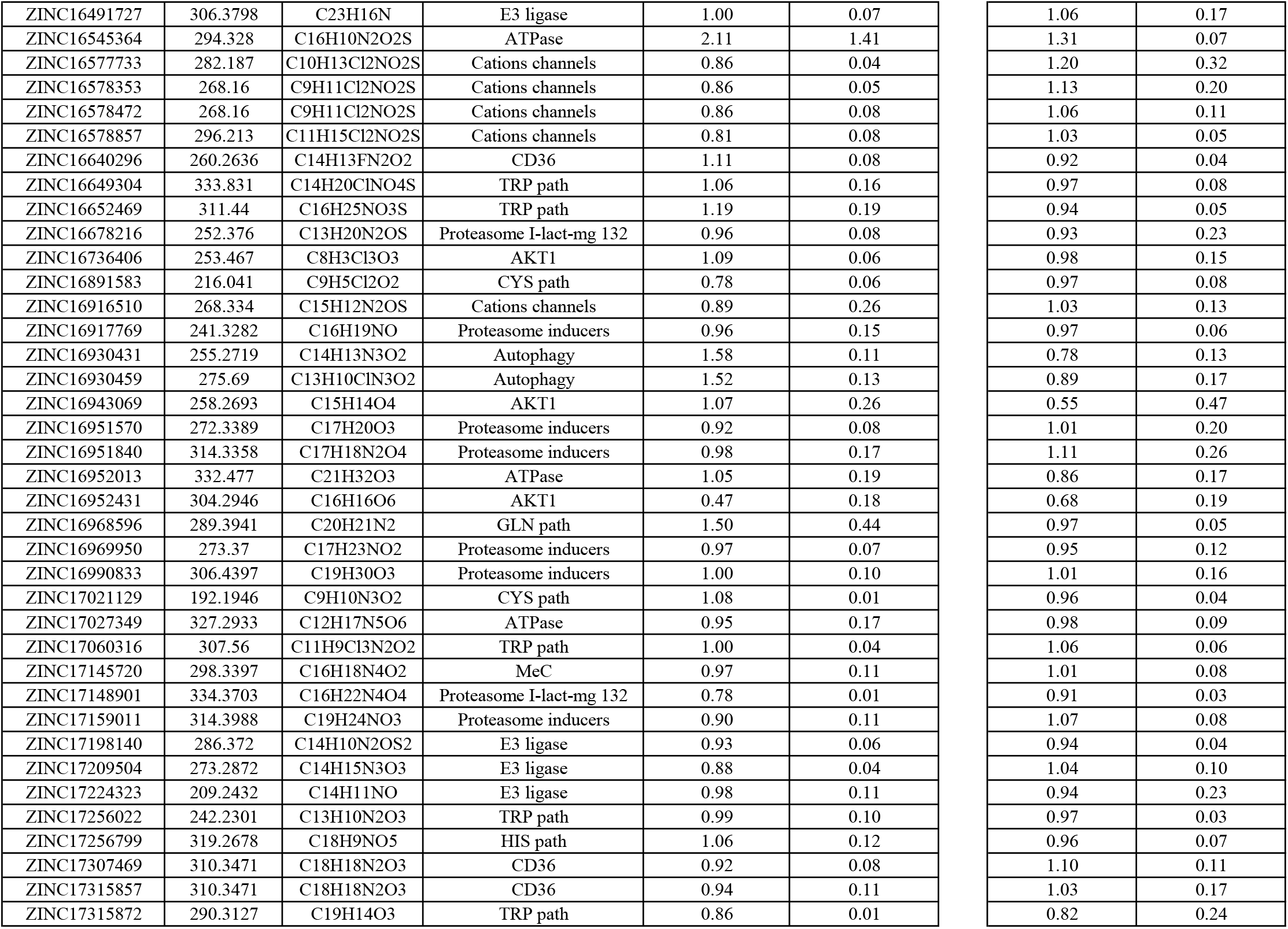
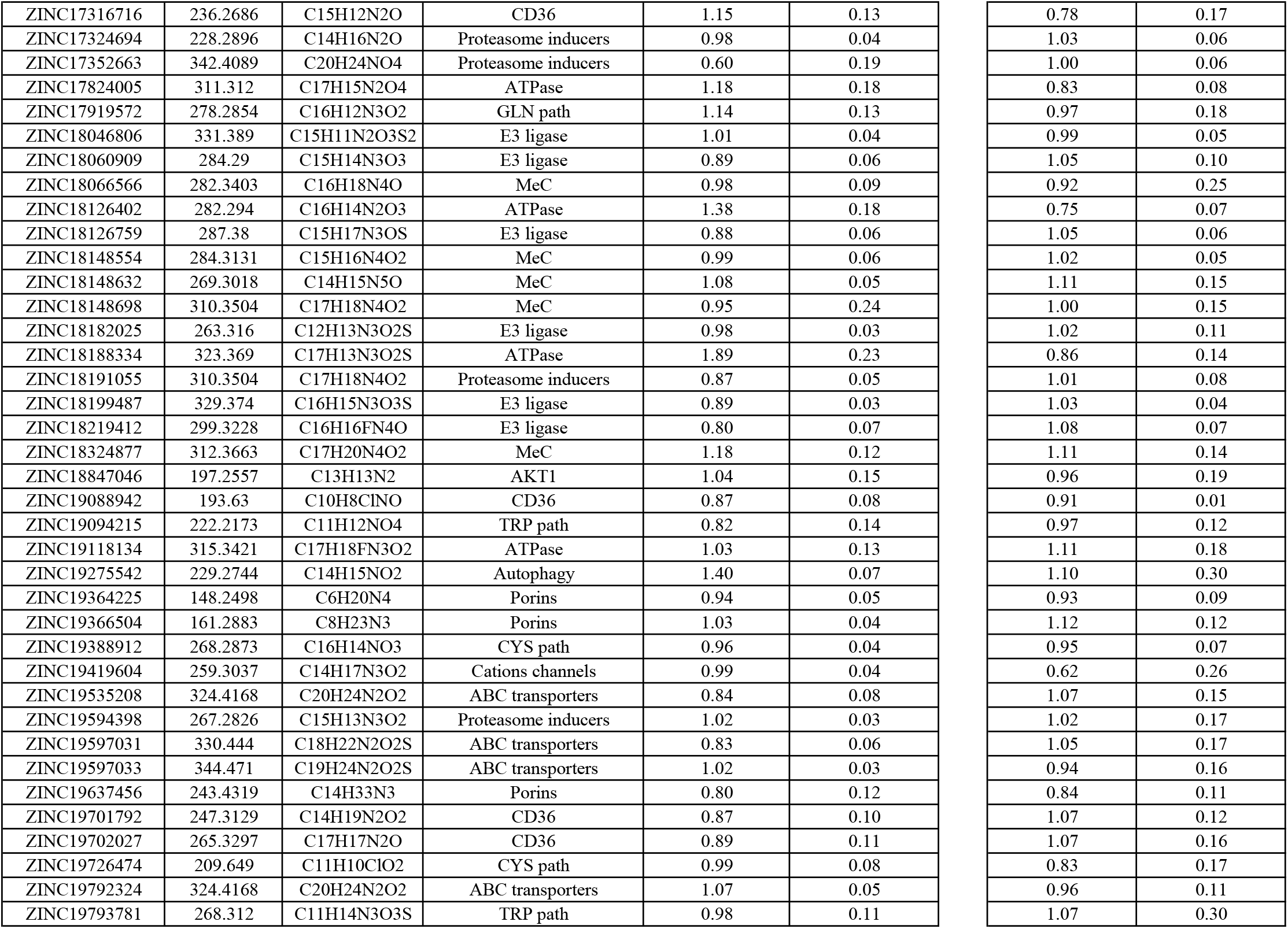
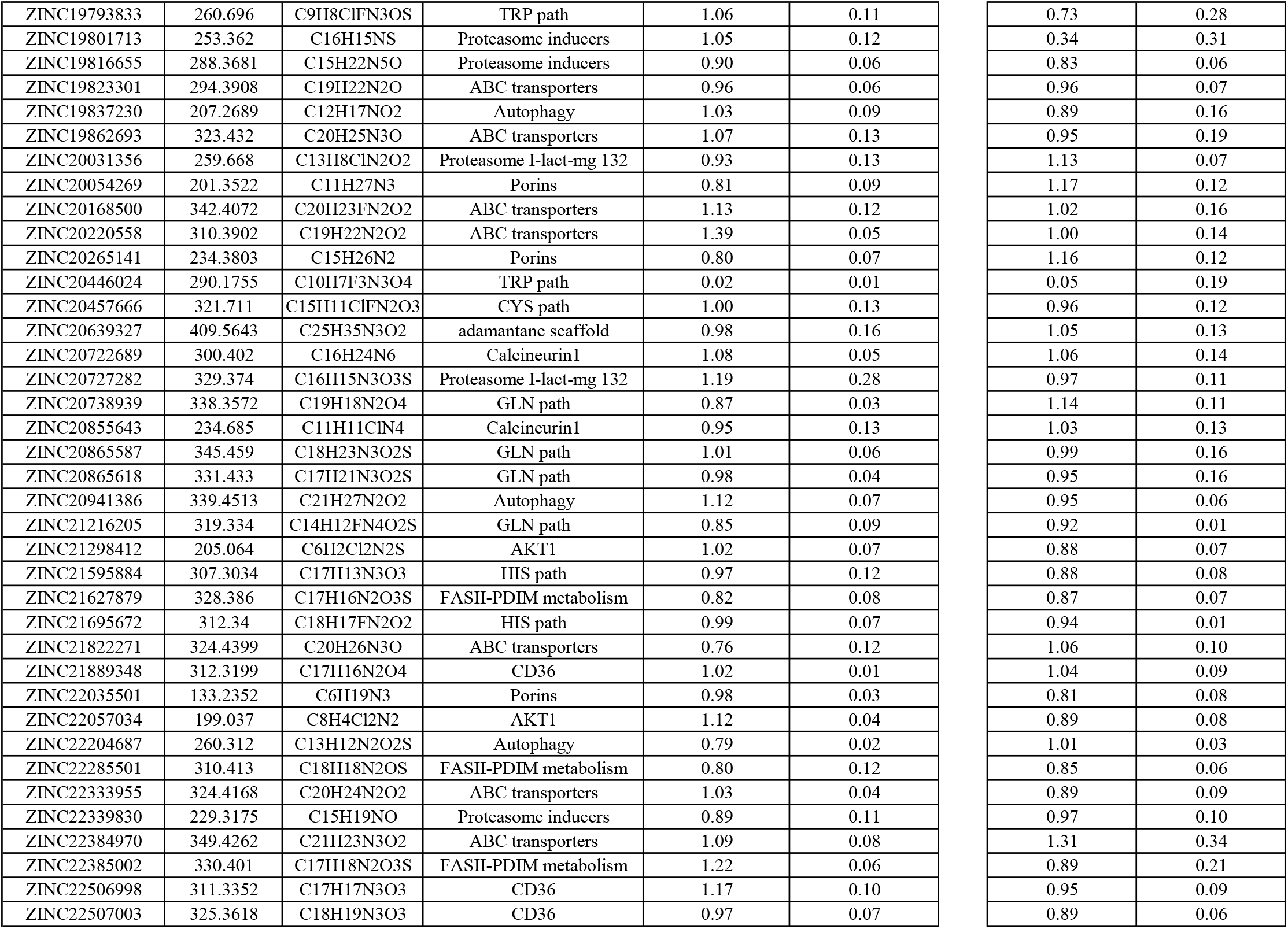
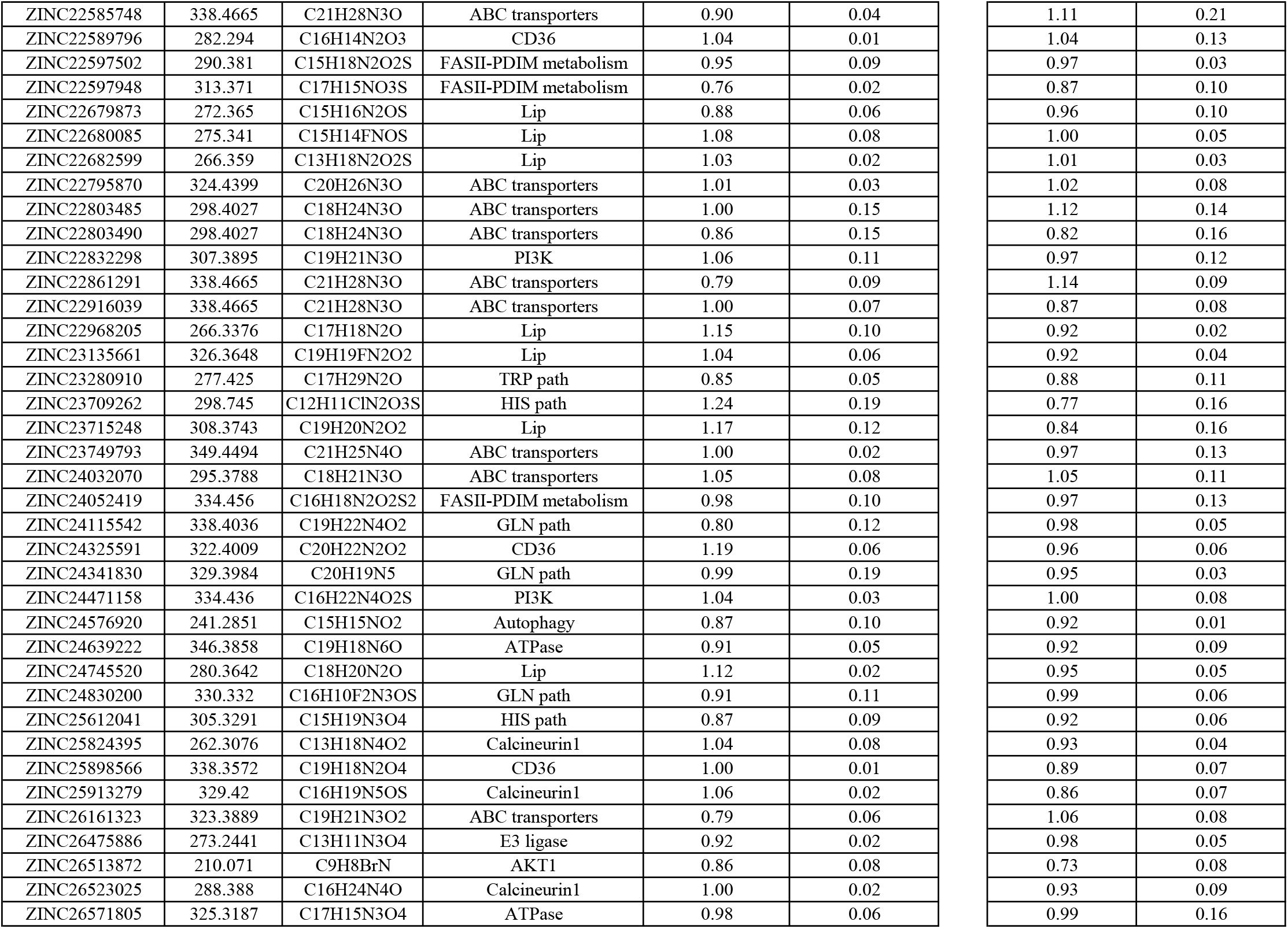
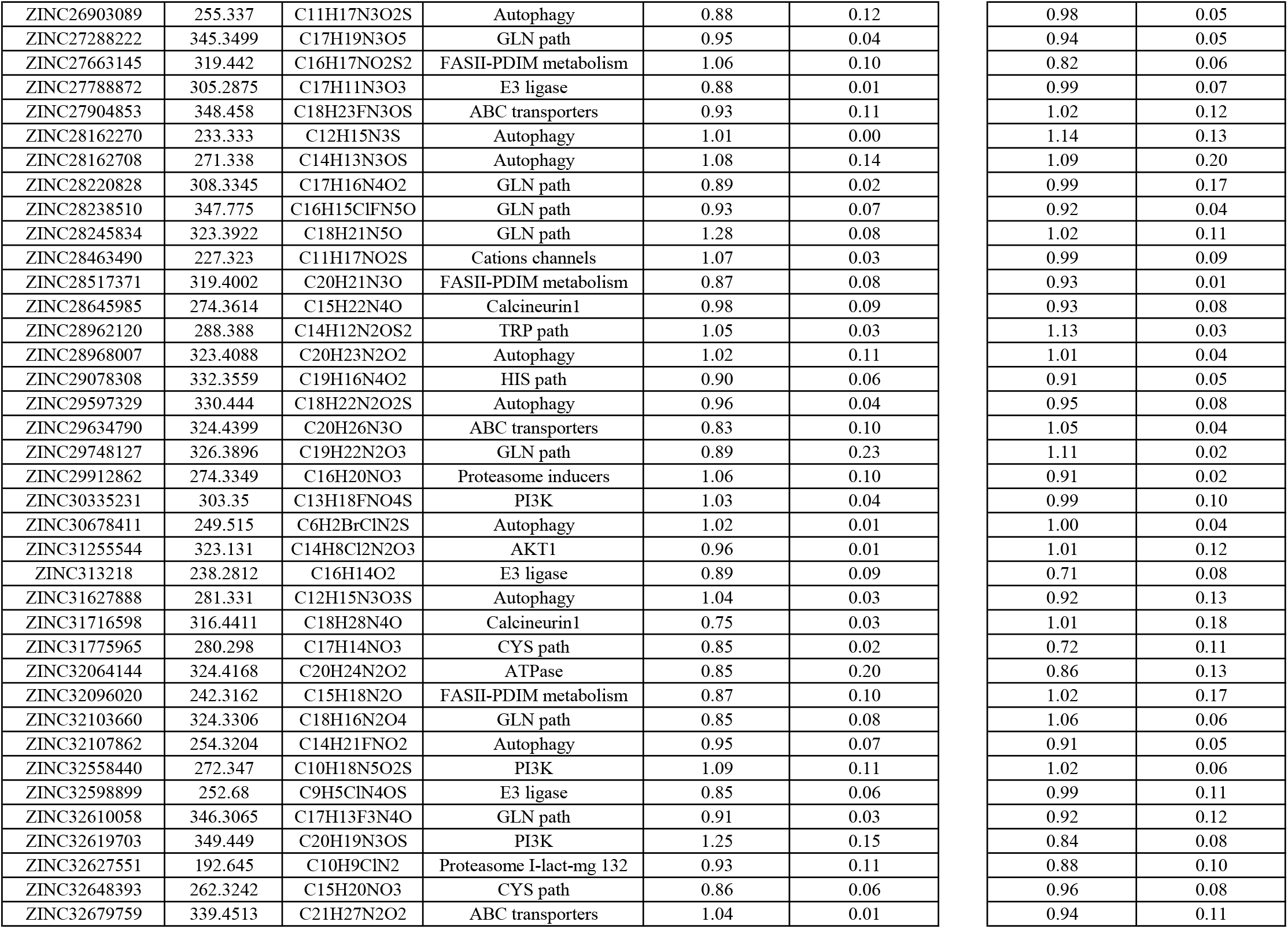
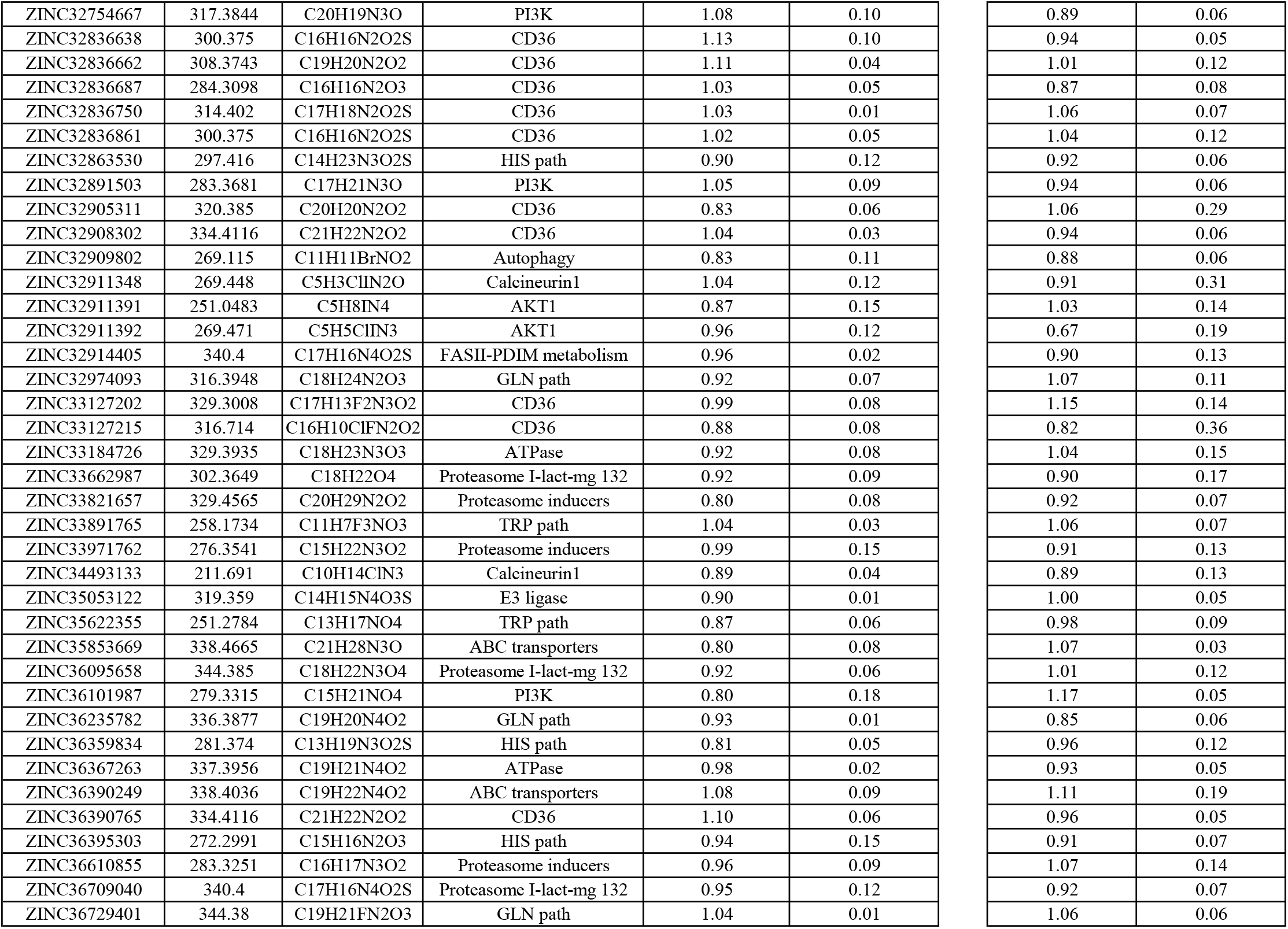
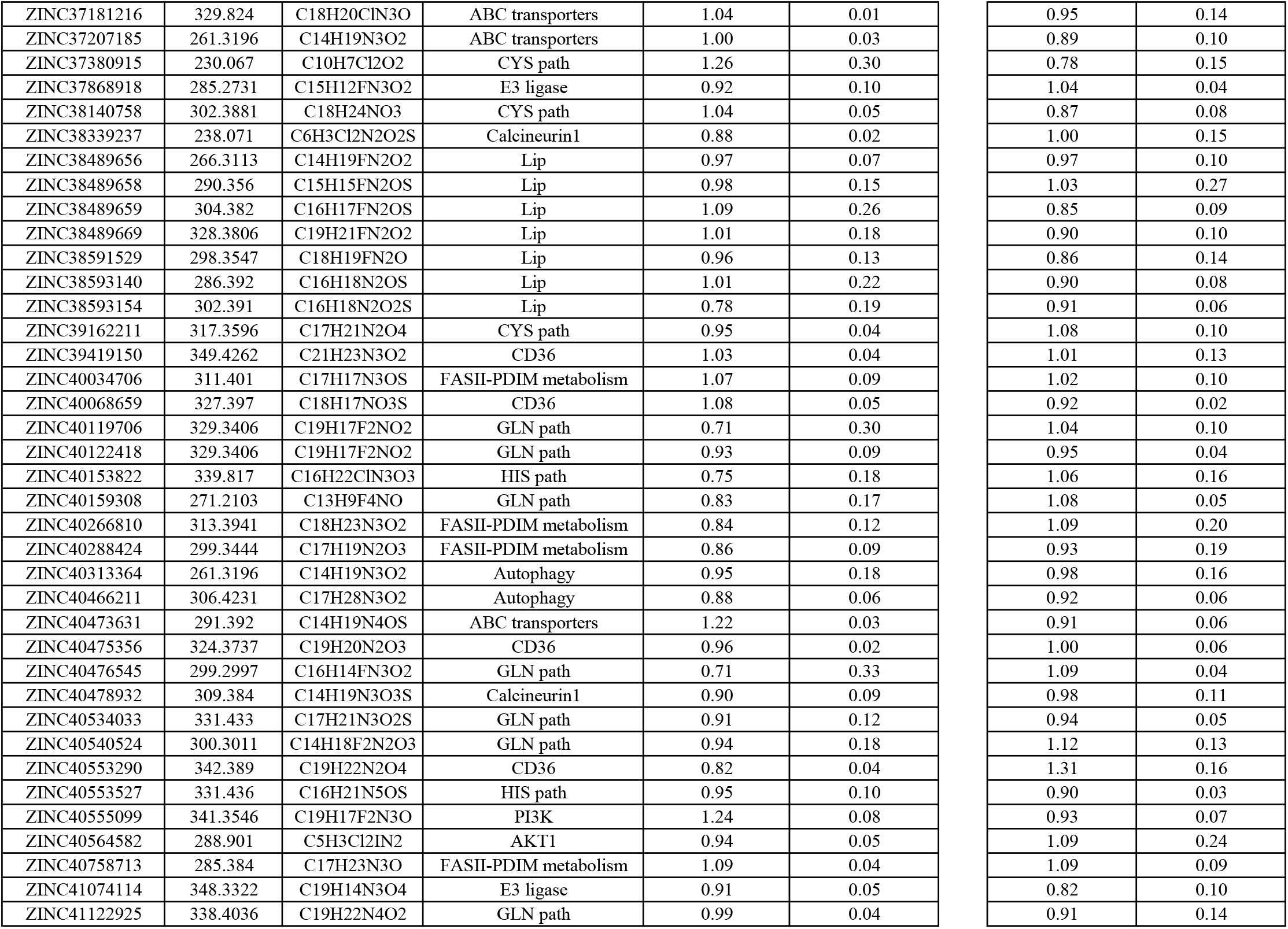
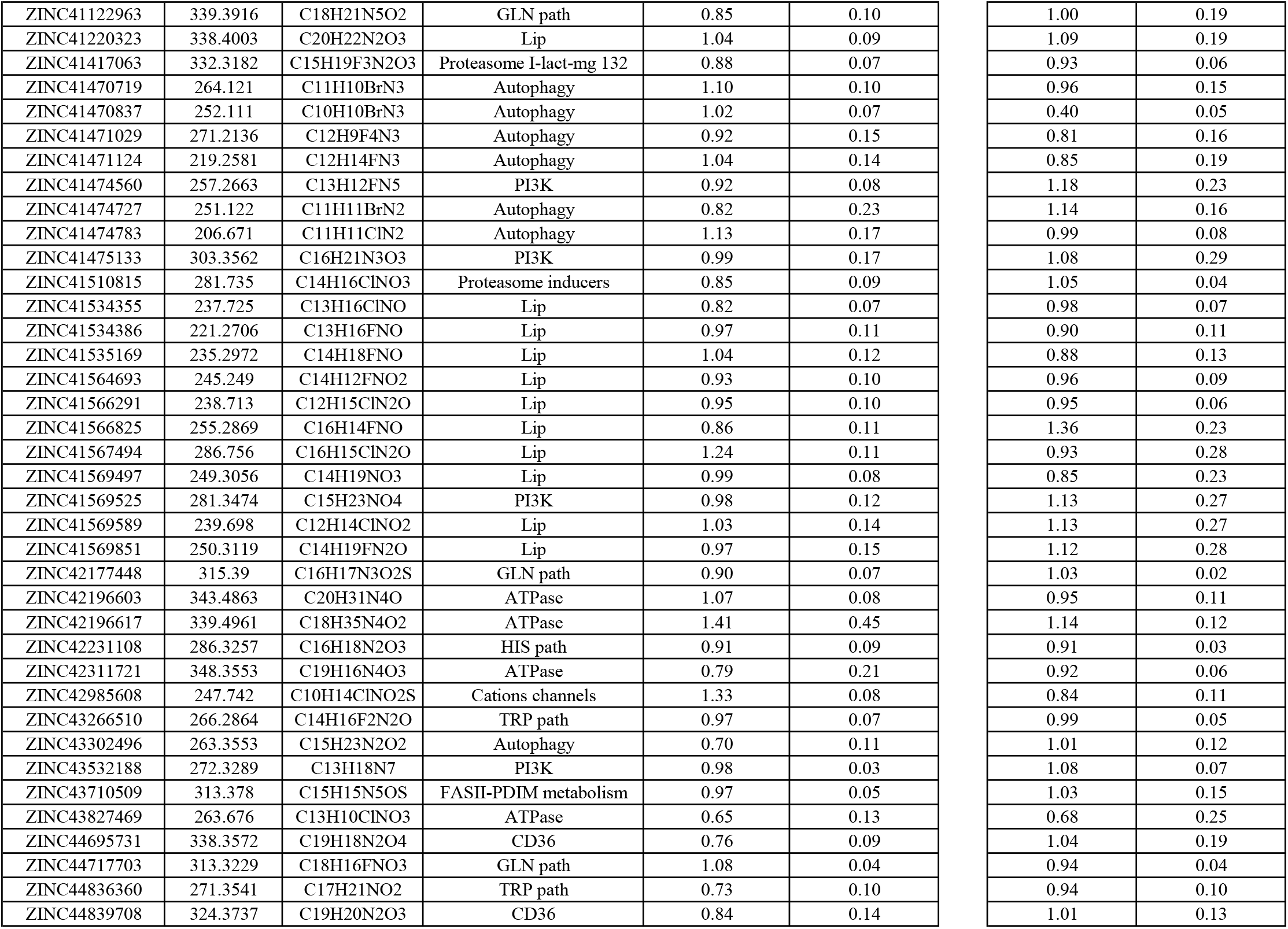
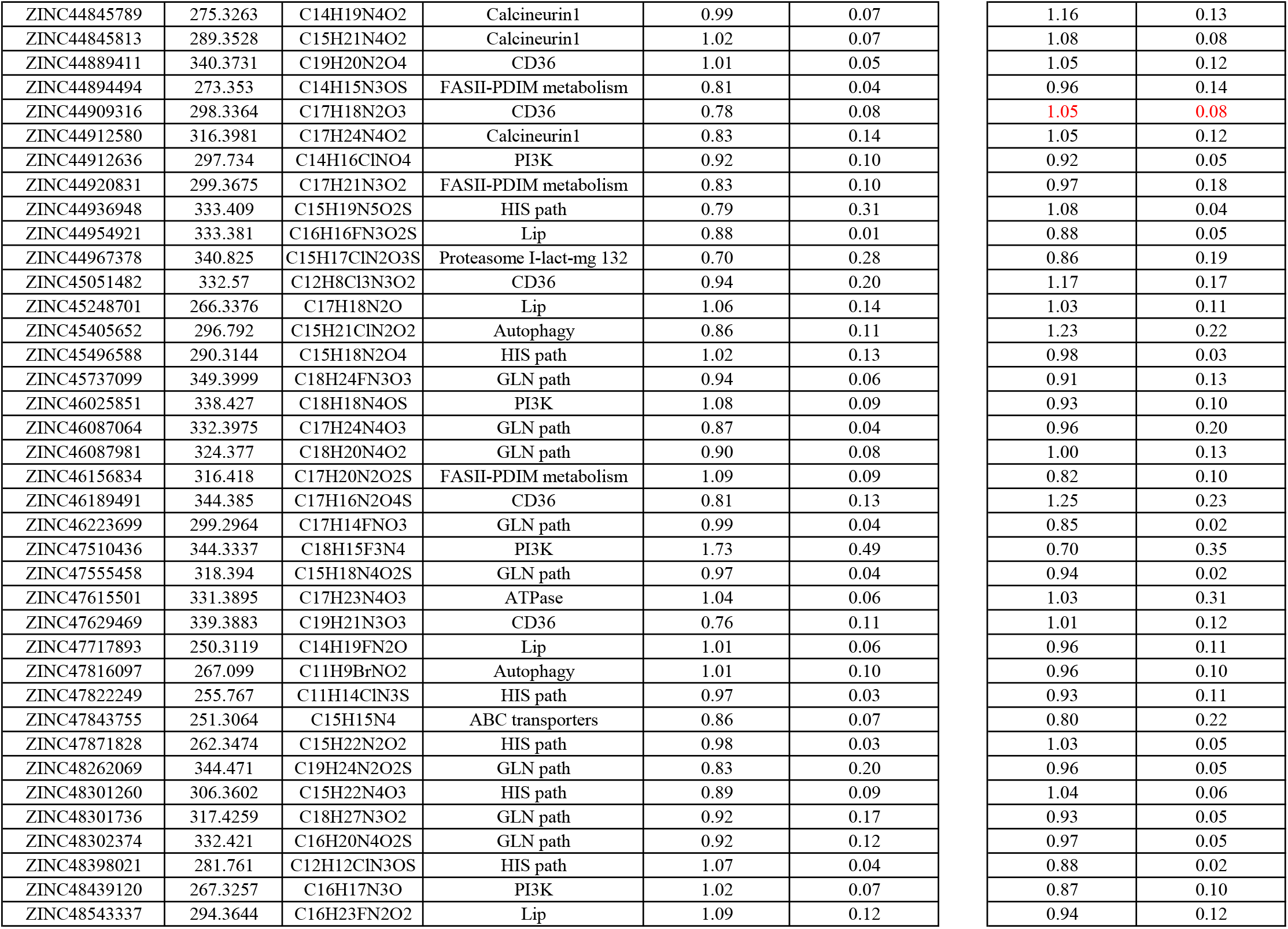
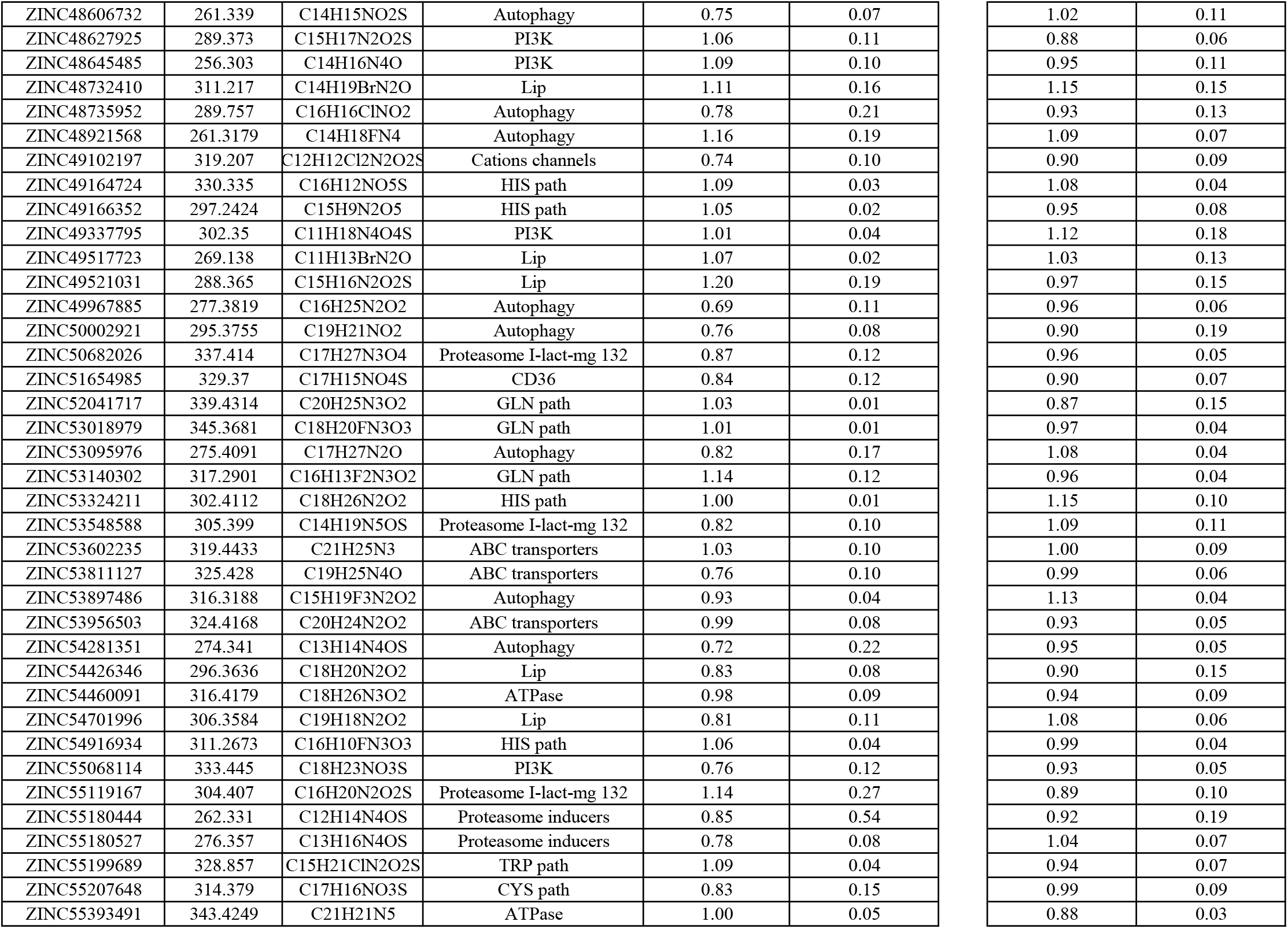
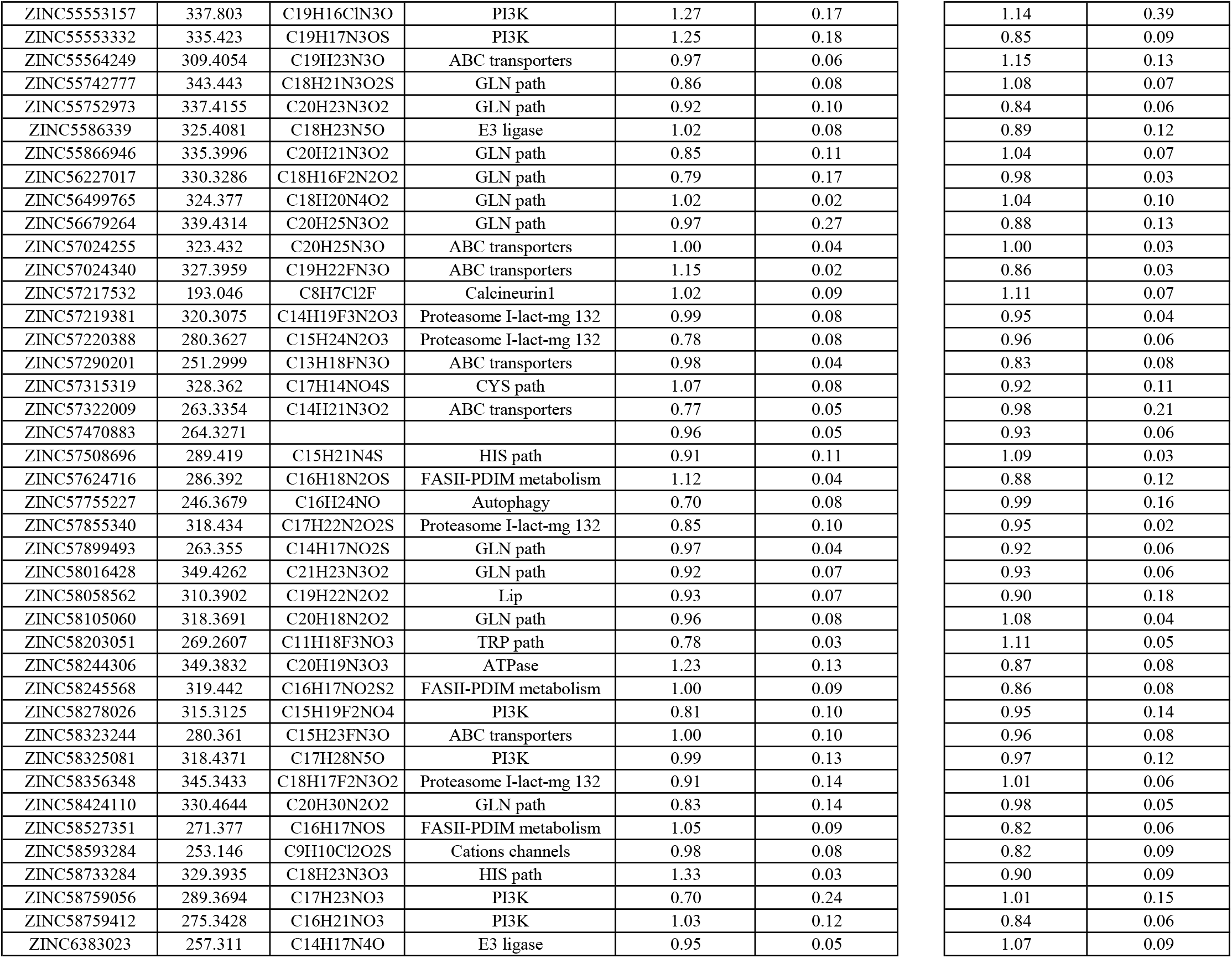

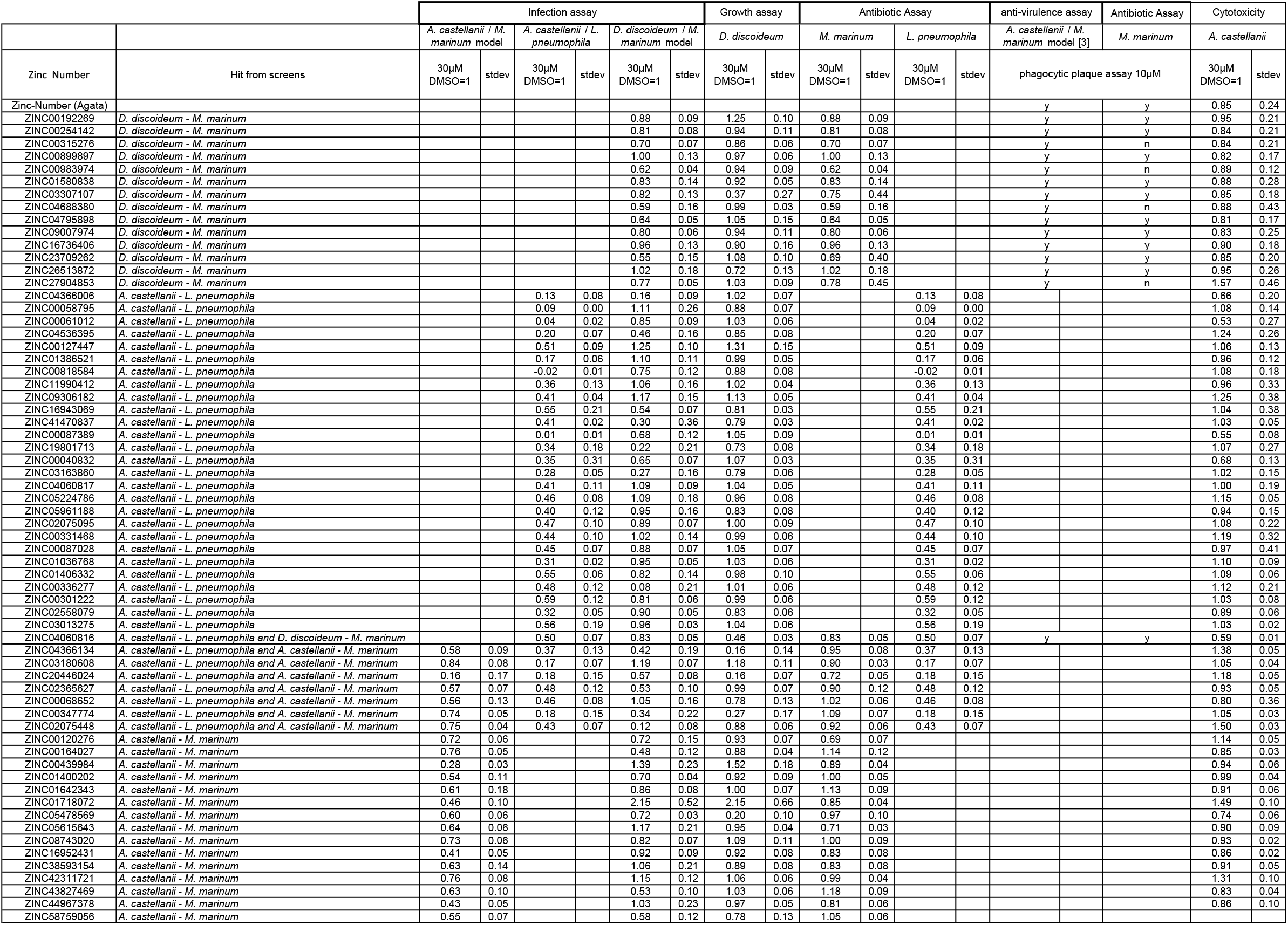

|  |  | Infection assay |  |  |  |  |  | Growth assay |  | Antibiotic Assay |  |  |  | anti-virulence assay | Antibiotic Assay | Cytotoxicity |  |
| --- | --- | --- | --- | --- | --- | --- | --- | --- | --- | --- | --- | --- | --- | --- | --- | --- | --- |
|  |  | A. castellanii / M. marinum model |  | A. castellanii / L. pneumophila |  | D. discoideum / M. marinum model |  | D. discoideum |  | M. marinum |  | L. pneumophila |  | A. castellanii / M. marinum model [3] | M. marinum | A. castellanii |  |
| Zinc Number | Hit from screens | 30µM DMSO=1 | stdev | 30µM DMSO=1 | stdev | 30µM DMSO=1 | stdev | 30µM DMSO=1 | stdev | 30µM DMSO=1 | stdev | 30µM DMSO=1 | stdev | phagocytic plaque assay 10µM |  | 30µM DMSO=1 | stdev |
| Zinc-Number (Agata) |  |  |  |  |  |  |  |  |  |  |  |  |  | y | y | 0.85 | 0.24 |
| ZINC00192269 | D. discoideum - M. marinum |  |  |  |  | 0.88 | 0.09 | 1.25 | 0.10 | 0.88 | 0.09 |  |  | y | y | 0.95 | 0.21 |
| ZINC00254142 | D. discoideum - M. marinum |  |  |  |  | 0.81 | 0.08 | 0.94 | 0.11 | 0.81 | 0.08 |  |  | y | y | 0.84 | 0.21 |
| ZINC00315276 | D. discoideum - M. marinum |  |  |  |  | 0.70 | 0.07 | 0.86 | 0.06 | 0.70 | 0.07 |  |  | y | n | 0.84 | 0.21 |
| ZINC00899897 | D. discoideum - M. marinum |  |  |  |  | 1.00 | 0.13 | 0.97 | 0.06 | 1.00 | 0.13 |  |  | y | y | 0.82 | 0.17 |
| ZINC00983974 | D. discoideum - M. marinum |  |  |  |  | 0.62 | 0.04 | 0.94 | 0.09 | 0.62 | 0.04 |  |  | y | n | 0.89 | 0.12 |
| ZINC01580838 | D. discoideum - M. marinum |  |  |  |  | 0.83 | 0.14 | 0.92 | 0.05 | 0.83 | 0.14 |  |  | y | y | 0.88 | 0.28 |
| ZINC03307107 | D. discoideum - M. marinum |  |  |  |  | 0.82 | 0.13 | 0.37 | 0.27 | 0.75 | 0.44 |  |  | y | y | 0.85 | 0.18 |
| ZINC04688380 | D. discoideum - M. marinum |  |  |  |  | 0.59 | 0.16 | 0.99 | 0.03 | 0.59 | 0.16 |  |  | y | n | 0.88 | 0.43 |
| ZINC04795898 | D. discoideum - M. marinum |  |  |  |  | 0.64 | 0.05 | 1.05 | 0.15 | 0.64 | 0.05 |  |  | y | y | 0.81 | 0.17 |
| ZINC09007974 | D. discoideum - M. marinum |  |  |  |  | 0.80 | 0.06 | 0.94 | 0.11 | 0.80 | 0.06 |  |  | y | y | 0.83 | 0.25 |
| ZINC16736406 | D. discoideum - M. marinum |  |  |  |  | 0.96 | 0.13 | 0.90 | 0.16 | 0.96 | 0.13 |  |  | y | y | 0.90 | 0.18 |
| ZINC23709262 | D. discoideum - M. marinum |  |  |  |  | 0.55 | 0.15 | 1.08 | 0.10 | 0.69 | 0.40 |  |  | y | y | 0.85 | 0.20 |
| ZINC26513872 | D. discoideum - M. marinum |  |  |  |  | 1.02 | 0.18 | 0.72 | 0.13 | 1.02 | 0.18 |  |  | y | y | 0.95 | 0.26 |
| ZINC27904853 | D. discoideum - M. marinum |  |  |  |  | 0.77 | 0.05 | 1.03 | 0.09 | 0.78 | 0.45 |  |  | y | n | 1.57 | 0.46 |
| ZINC04366006 | A. castellanii - L. pneumophila |  |  | 0.13 | 0.08 | 0.16 | 0.09 | 1.02 | 0.07 |  |  | 0.13 | 0.08 |  |  | 0.66 | 0.20 |
| ZINC00058795 | A. castellanii - L. pneumophila |  |  | 0.09 | 0.00 | 1.11 | 0.26 | 0.88 | 0.07 |  |  | 0.09 | 0.00 |  |  | 1.08 | 0.14 |
| ZINC00061012 | A. castellanii - L. pneumophila |  |  | 0.04 | 0.02 | 0.85 | 0.09 | 1.03 | 0.06 |  |  | 0.04 | 0.02 |  |  | 0.53 | 0.27 |
| ZINC04536395 | A. castellanii - L. pneumophila |  |  | 0.20 | 0.07 | 0.46 | 0.16 | 0.85 | 0.08 |  |  | 0.20 | 0.07 |  |  | 1.24 | 0.26 |
| ZINC00127447 | A. castellanii - L. pneumophila |  |  | 0.51 | 0.09 | 1.25 | 0.10 | 1.31 | 0.15 |  |  | 0.51 | 0.09 |  |  | 1.06 | 0.13 |
| ZINC01386521 | A. castellanii - L. pneumophila |  |  | 0.17 | 0.06 | 1.10 | 0.11 | 0.99 | 0.05 |  |  | 0.17 | 0.06 |  |  | 0.96 | 0.12 |
| ZINC00818584 | A. castellanii - L. pneumophila |  |  | -0.02 | 0.01 | 0.75 | 0.12 | 0.88 | 0.08 |  |  | -0.02 | 0.01 |  |  | 1.08 | 0.18 |
| ZINC11990412 | A. castellanii - L. pneumophila |  |  | 0.36 | 0.13 | 1.06 | 0.16 | 1.02 | 0.04 |  |  | 0.36 | 0.13 |  |  | 0.96 | 0.33 |
| ZINC09306182 | A. castellanii - L. pneumophila |  |  | 0.41 | 0.04 | 1.17 | 0.15 | 1.13 | 0.05 |  |  | 0.41 | 0.04 |  |  | 1.25 | 0.38 |
| ZINC16943069 | A. castellanii - L. pneumophila |  |  | 0.55 | 0.21 | 0.54 | 0.07 | 0.81 | 0.03 |  |  | 0.55 | 0.21 |  |  | 1.04 | 0.38 |
| ZINC41470837 | A. castellanii - L. pneumophila |  |  | 0.41 | 0.02 | 0.30 | 0.36 | 0.79 | 0.03 |  |  | 0.41 | 0.02 |  |  | 1.03 | 0.05 |
| ZINC00087389 | A. castellanii - L. pneumophila |  |  | 0.01 | 0.01 | 0.68 | 0.12 | 1.05 | 0.09 |  |  | 0.01 | 0.01 |  |  | 0.55 | 0.08 |
| ZINC19801713 | A. castellanii - L. pneumophila |  |  | 0.34 | 0.18 | 0.22 | 0.21 | 0.73 | 0.08 |  |  | 0.34 | 0.18 |  |  | 1.07 | 0.27 |
| ZINC00040832 | A. castellanii - L. pneumophila |  |  | 0.35 | 0.31 | 0.65 | 0.07 | 1.07 | 0.03 |  |  | 0.35 | 0.31 |  |  | 0.68 | 0.13 |
| ZINC03163860 | A. castellanii - L. pneumophila |  |  | 0.28 | 0.05 | 0.27 | 0.16 | 0.79 | 0.06 |  |  | 0.28 | 0.05 |  |  | 1.02 | 0.15 |
| ZINC04060817 | A. castellanii - L. pneumophila |  |  | 0.41 | 0.11 | 1.09 | 0.09 | 1.04 | 0.05 |  |  | 0.41 | 0.11 |  |  | 1.00 | 0.19 |
| ZINC05224786 | A. castellanii - L. pneumophila |  |  | 0.46 | 0.08 | 1.09 | 0.18 | 0.96 | 0.08 |  |  | 0.46 | 0.08 |  |  | 1.15 | 0.05 |
| ZINC05961188 | A. castellanii - L. pneumophila |  |  | 0.40 | 0.12 | 0.95 | 0.16 | 0.83 | 0.08 |  |  | 0.40 | 0.12 |  |  | 0.94 | 0.15 |
| ZINC02075095 | A. castellanii - L. pneumophila |  |  | 0.47 | 0.10 | 0.89 | 0.07 | 1.00 | 0.09 |  |  | 0.47 | 0.10 |  |  | 1.08 | 0.22 |
| ZINC00331468 | A. castellanii - L. pneumophila |  |  | 0.44 | 0.10 | 1.02 | 0.14 | 0.99 | 0.06 |  |  | 0.44 | 0.10 |  |  | 1.19 | 0.32 |
| ZINC00087028 | A. castellanii - L. pneumophila |  |  | 0.45 | 0.07 | 0.88 | 0.07 | 1.05 | 0.07 |  |  | 0.45 | 0.07 |  |  | 0.97 | 0.41 |
| ZINC01036768 | A. castellanii - L. pneumophila |  |  | 0.31 | 0.02 | 0.95 | 0.05 | 1.03 | 0.06 |  |  | 0.31 | 0.02 |  |  | 1.10 | 0.09 |
| ZINC01406332 | A. castellanii - L. pneumophila |  |  | 0.55 | 0.06 | 0.82 | 0.14 | 0.98 | 0.10 |  |  | 0.55 | 0.06 |  |  | 1.09 | 0.06 |
| ZINC00336277 | A. castellanii - L. pneumophila |  |  | 0.48 | 0.12 | 0.08 | 0.21 | 1.01 | 0.06 |  |  | 0.48 | 0.12 |  |  | 1.12 | 0.21 |
| ZINC00301222 | A. castellanii - L. pneumophila |  |  | 0.59 | 0.12 | 0.81 | 0.06 | 0.99 | 0.06 |  |  | 0.59 | 0.12 |  |  | 1.03 | 0.08 |
| ZINC02558079 | A. castellanii - L. pneumophila |  |  | 0.32 | 0.05 | 0.90 | 0.05 | 0.83 | 0.06 |  |  | 0.32 | 0.05 |  |  | 0.89 | 0.06 |
| ZINC03013275 | A. castellanii - L. pneumophila |  |  | 0.56 | 0.19 | 0.96 | 0.03 | 1.04 | 0.06 |  |  | 0.56 | 0.19 |  |  | 1.03 | 0.02 |
| ZINC04060816 | A. castellanii - L. pneumophila and D. discoideum - M. marinum |  |  | 0.50 | 0.07 | 0.83 | 0.05 | 0.46 | 0.03 | 0.83 | 0.05 | 0.50 | 0.07 | y | y | 0.59 | 0.01 |
| ZINC04366134 | A. castellanii - L. pneumophila and A. castellanii - M. marinum | 0.58 | 0.09 | 0.37 | 0.13 | 0.42 | 0.19 | 0.16 | 0.14 | 0.95 | 0.08 | 0.37 | 0.13 |  |  | 1.38 | 0.05 |
| ZINC03180608 | A. castellanii - L. pneumophila and A. castellanii - M. marinum | 0.84 | 0.08 | 0.17 | 0.07 | 1.19 | 0.07 | 1.18 | 0.11 | 0.90 | 0.03 | 0.17 | 0.07 |  |  | 1.05 | 0.04 |
| ZINC20446024 | A. castellanii - L. pneumophila and A. castellanii - M. marinum | 0.16 | 0.17 | 0.18 | 0.15 | 0.57 | 0.08 | 0.16 | 0.07 | 0.72 | 0.05 | 0.18 | 0.15 |  |  | 1.18 | 0.05 |
| ZINC02365627 | A. castellanii - L. pneumophila and A. castellanii - M. marinum | 0.57 | 0.07 | 0.48 | 0.12 | 0.53 | 0.10 | 0.99 | 0.07 | 0.90 | 0.12 | 0.48 | 0.12 |  |  | 0.93 | 0.05 |
| ZINC00068652 | A. castellanii - L. pneumophila and A. castellanii - M. marinum | 0.56 | 0.13 | 0.46 | 0.08 | 1.05 | 0.16 | 0.78 | 0.13 | 1.02 | 0.06 | 0.46 | 0.08 |  |  | 0.80 | 0.36 |
| ZINC00347774 | A. castellanii - L. pneumophila and A. castellanii - M. marinum | 0.74 | 0.05 | 0.18 | 0.15 | 0.34 | 0.22 | 0.27 | 0.17 | 1.09 | 0.07 | 0.18 | 0.15 |  |  | 1.05 | 0.03 |
| ZINC02075448 | A. castellanii - L. pneumophila and A. castellanii - M. marinum | 0.75 | 0.04 | 0.43 | 0.07 | 0.12 | 0.08 | 0.88 | 0.06 | 0.92 | 0.06 | 0.43 | 0.07 |  |  | 1.50 | 0.03 |
| ZINC00120276 | A. castellanii - M. marinum | 0.72 | 0.06 |  |  | 0.72 | 0.15 | 0.93 | 0.07 | 0.69 | 0.07 |  |  |  |  | 1.14 | 0.05 |
| ZINC00164027 | A. castellanii - M. marinum | 0.76 | 0.05 |  |  | 0.48 | 0.12 | 0.88 | 0.04 | 1.14 | 0.12 |  |  |  |  | 0.85 | 0.03 |
| ZINC00439984 | A. castellanii - M. marinum | 0.28 | 0.03 |  |  | 1.39 | 0.23 | 1.52 | 0.18 | 0.89 | 0.04 |  |  |  |  | 0.94 | 0.06 |
| ZINC01400202 | A. castellanii - M. marinum | 0.54 | 0.11 |  |  | 0.70 | 0.04 | 0.92 | 0.09 | 1.00 | 0.05 |  |  |  |  | 0.99 | 0.04 |
| ZINC01642343 | A. castellanii - M. marinum | 0.61 | 0.18 |  |  | 0.86 | 0.08 | 1.00 | 0.07 | 1.13 | 0.09 |  |  |  |  | 0.91 | 0.06 |
| ZINC01718072 | A. castellanii - M. marinum | 0.46 | 0.10 |  |  | 2.15 | 0.52 | 2.15 | 0.66 | 0.85 | 0.04 |  |  |  |  | 1.49 | 0.10 |
| ZINC05478569 | A. castellanii - M. marinum | 0.60 | 0.06 |  |  | 0.72 | 0.03 | 0.20 | 0.10 | 0.97 | 0.10 |  |  |  |  | 0.74 | 0.06 |
| ZINC05615643 | A. castellanii - M. marinum | 0.64 | 0.06 |  |  | 1.17 | 0.21 | 0.95 | 0.04 | 0.71 | 0.03 |  |  |  |  | 0.90 | 0.09 |
| ZINC08743020 | A. castellanii - M. marinum | 0.73 | 0.06 |  |  | 0.82 | 0.07 | 1.09 | 0.11 | 1.00 | 0.09 |  |  |  |  | 0.93 | 0.02 |
| ZINC16952431 | A. castellanii - M. marinum | 0.41 | 0.05 |  |  | 0.92 | 0.09 | 0.92 | 0.08 | 0.83 | 0.08 |  |  |  |  | 0.86 | 0.02 |
| ZINC38593154 | A. castellanii - M. marinum | 0.63 | 0.14 |  |  | 1.06 | 0.21 | 0.89 | 0.08 | 0.83 | 0.08 |  |  |  |  | 0.91 | 0.05 |
| ZINC42311721 | A. castellanii - M. marinum | 0.76 | 0.08 |  |  | 1.15 | 0.12 | 1.06 | 0.06 | 0.99 | 0.04 |  |  |  |  | 1.31 | 0.10 |
| ZINC43827469 | A. castellanii - M. marinum | 0.63 | 0.10 |  |  | 0.53 | 0.10 | 1.03 | 0.06 | 1.18 | 0.09 |  |  |  |  | 0.83 | 0.04 |
| ZINC44967378 | A. castellanii - M. marinum | 0.43 | 0.05 |  |  | 1.03 | 0.23 | 0.97 | 0.05 | 0.81 | 0.06 |  |  |  |  | 0.86 | 0.10 |
| ZINC58759056 | A. castellanii - M. marinum | 0.55 | 0.07 |  |  | 0.58 | 0.12 | 0.78 | 0.13 | 1.05 | 0.06 |  |  |  |  |  |  |

## References

Alibaud, L., Y. Rombouts, X. Trivelli, A. Burguiere, S. L. Cirillo, J. D. Cirillo, J. F. Dubremetz, Y. Guerardel, G. Lutfalla, and L. Kremer. 2011. ‘A Mycobacterium marinum TesA mutant defective for major cell wall-associated lipids is highly attenuated in Dictyostelium discoideum and zebrafish embryos’, Mol Microbiol, 80: 919–34.

Ballell, L., R. H. Bates, R. J. Young, D. Alvarez-Gomez, E. Alvarez-Ruiz, V. Barroso, D. Blanco, B. Crespo, J. Escribano, R. Gonzalez, S. Lozano, S. Huss, A. Santos-Villarejo, J. J. Martin-Plaza, A. Mendoza, M. J. Rebollo-Lopez, M. Remuinan-Blanco, J. L. Lavandera, E. Perez-Herran, F. J. Gamo-Benito, J. F. Garcia-Bustos, D. Barros, J. P. Castro, and N. Cammack. 2013. ‘Fueling open-source drug discovery: 177 small-molecule leads against tuberculosis’, ChemMedChem, 8: 313–21.

Barisch, C., and T. Soldati. 2017. ‘Mycobacterium marinum Degrades Both Triacylglycerols and Phospholipids from Its Dictyostelium Host to Synthesise Its Own Triacylglycerols and Generate Lipid Inclusions’, PLoS Pathog, 13: e1006095.

Bertrand, T., F. Auge, J. Houtmann, A. Rak, F. Vallee, V. Mikol, P. F. Berne, N. Michot, D. Cheuret, C. Hoornaert, and M. Mathieu. 2010. ‘Structural basis for human monoglyceride lipase inhibition’, J Mol Biol, 396: 663–73.

Botella, H., P. Peyron, F. Levillain, R. Poincloux, Y. Poquet, I. Brandli, C. Wang, L. Tailleux, S. Tilleul, G. M. Charriere, S. J. Waddell, M. Foti, G. Lugo-Villarino, Q. Gao, I. Maridonneau-Parini, P. D. Butcher, P. R. Castagnoli, B. Gicquel, C. de Chastellier, and O. Neyrolles. 2011. ‘Mycobacterial p(1)-type ATPases mediate resistance to zinc poisoning in human macrophages’, Cell Host Microbe, 10: 248–59.

Boulais, J., M. Trost, C. R. Landry, R. Dieckmann, E. D. Levy, T. Soldati, S. W. Michnick, P. Thibault, and M. Desjardins. 2010. ‘Molecular characterization of the evolution of phagosomes’, Mol Syst Biol, 6: 423.

Bozzaro, S., C. Bucci, and M. Steinert. 2008. ‘Phagocytosis and host-pathogen interactions in Dictyostelium with a look at macrophages’, Int Rev Cell Mol Biol, 271: 253–300.

Bozzaro, S., S. Buracco, and B. Peracino. 2013. ‘Iron metabolism and resistance to infection by invasive bacteria in the social amoeba Dictyostelium discoideum’, Front Cell Infect Microbiol, 3: 50.

Bravo-Toncio, C., J. A. Alvarez, F. Campos, J. Ortiz-Severin, M. Varas, R. Cabrera, C. F. Lagos, and F. P. Chavez. 2016. ‘Dictyostelium discoideum as a surrogate host-microbe model for antivirulence screening in Pseudomonas aeruginosa PAO1’, Int J Antimicrob Agents, 47: 403–9.

Briken, V., S. E. Ahlbrand, and S. Shah. 2013. ‘Mycobacterium tuberculosis and the host cell inflammasome: a complex relationship’, Front Cell Infect Microbiol, 3: 62.

Bustanji, Y., I. M. Al-Masri, M. Mohammad, M. Hudaib, K. Tawaha, H. Tarazi, and H. S. Alkhatib. 2011. ‘Pancreatic lipase inhibition activity of trilactone terpenes of Ginkgo biloba’, J Enzyme Inhib Med Chem, 26: 453–9.

Cardenal-Munoz, E., C. Barisch, L. H. Lefrancois, A. T. Lopez-Jimenez, and T. Soldati. 2017. ‘When Dicty Met Myco, a (Not So) Romantic Story about One Amoeba and Its Intracellular Pathogen’, Front Cell Infect Microbiol, 7: 529.

Carlet, Jean, Claude Rambaud, and Céline Pulcini. 2014. ‘Save antibiotics: a call for action of the world alliance against antibiotic resistance (WAAAR)’, BMC infectious diseases, 14: 436.

Chen, Qianyi, Vashti C. Bryant, Hernando Lopez, David L. Kelly, Xu Luo, and Amarnath Natarajan. 2011. ‘2,3-Substituted quinoxalin-6-amine analogs as antiproliferatives: A structure activity relationship study’, Bioorganic & medicinal chemistry letters, 21: 1929–32.

Cho, Y., T. R. Ioerger, and J. C. Sacchettini. 2008. ‘Discovery of novel nitrobenzothiazole inhibitors for Mycobacterium tuberculosis ATP phosphoribosyl transferase (HisG) through virtual screening’, J Med Chem, 51: 5984–92.

Christelle, B., O. Eduardo Bde, C. Latifa, M. Elaine-Rose, M. Bernard, R. H. Evelyne, G. Mohamed, E. Jean-Marc, and H. Catherine. 2011. ‘Combined docking and molecular dynamics simulations to enlighten the capacity of Pseudomonas cepacia and Candida antarctica lipases to catalyze quercetin acetylation’, J Biotechnol, 156: 203–10.

Coort, Susan LM, Jodil Willems, Will A Coumans, Ger J Van Der Vusse, Arend Bonen, Jan FC Glatz, and Joost JFP Luiken. 2002. ‘Sulfo-N-succinimidyl esters of long chain fatty acids specifically inhibit fatty acid translocase (FAT/CD36)-mediated cellular fatty acid uptake’, Molecular and cellular biochemistry, 239: 213–19.

Cosson, P., and T. Soldati. 2008. ‘Eat, kill or die: when amoeba meets bacteria’, Curr Opin Microbiol, 11: 271–6.

Cunha, Burke A, Gina Wu, and Muhammed Raza. 2015. ‘Clinical Diagnosis of Legionnaire’s Disease: Six Characteristic Clinical Predictors’, The American journal of medicine, 128: e21–e22.

Dall-Larsen, T., H. Kryvi, and L. Klungsoyr. 1976. ‘Dinitrophenol, dicoumarol and pentachlorophenol as inhibitors and parasite substrates in the ATP phosphoribosyltransferase reaction’, Eur J Biochem, 66: 443–6.

Design, Synthesis. 2009. ‘Biological Evaluation of Fluorinated Analogues of Salicylihalamide Sugimoto, Yoshinori; Konoki, Keiichi; Murata, Michio; Matsushita, Masafumi; Kanazawa, Hiroshi; Oishi’, Journal of Medicinal Chemistry, 52: 798–806.

Dheda, K., C. E. Barry, 3rd, and G. Maartens. 2016. ‘Tuberculosis’, Lancet, 387: 1211–26.

Diop, E. A., E. F. Queiroz, S. Kicka, S. Rudaz, T. Diop, T. Soldati, and J. L. Wolfender. 2018. ‘Survey on medicinal plants traditionally used in Senegal for the treatment of tuberculosis (TB) and assessment of their antimycobacterial activity’, J Ethnopharmacol, 216: 71–78.

Dunn, J. D., C. Bosmani, C. Barisch, L. Raykov, L. H. Lefrancois, E. Cardenal-Munoz, A. T. Lopez-Jimenez, and T. Soldati. 2017. ‘Eat Prey, Live: Dictyostelium discoideum As a Model for Cell-Autonomous Defenses’, Front Immunol, 8: 1906.

Escoll, P., M. Rolando, L. Gomez-Valero, and C. Buchrieser. 2013. ‘From amoeba to macrophages: exploring the molecular mechanisms of Legionella pneumophila infection in both hosts’, Curr Top Microbiol Immunol, 376: 1–34.

Farnsworth, N. R., O. Akerele, A. S. Bingel, D. D. Soejarto, and Z. Guo. 1985. ‘Medicinal plants in therapy’, Bull World Health Organ, 63: 965–81.

Floto, R Andres, Sovan Sarkar, Ethan O Perlstein, Beate Kampmann, Stuart L Schreiber, and David C Rubinsztein. 2007. ‘Small molecule enhancers of rapamycin-induced TOR inhibition promote autophagy, reduce toxicity in Huntington’s disease models and enhance killing of mycobacteria by macrophages’, Autophagy, 3: 620–22.

Froquet, R., E. Lelong, A. Marchetti, and P. Cosson. 2009. ‘Dictyostelium discoideum: a model host to measure bacterial virulence’, Nat Protoc, 4: 25–30.

Ghose, A. K., V. N. Viswanadhan, and J. J. Wendoloski. 1999. ‘A knowledge-based approach in designing combinatorial or medicinal chemistry libraries for drug discovery. 1. A qualitative and quantitative characterization of known drug databases’, J Comb Chem, 1: 55–68.

Gopaldass, N., D. Patel, R. Kratzke, R. Dieckmann, S. Hausherr, M. Hagedorn, R. Monroy, J. Kruger, E. M. Neuhaus, E. Hoffmann, K. Hille, S. A. Kuznetsov, and T. Soldati. 2012. ‘Dynamin A, Myosin IB and Abp1 couple phagosome maturation to F-actin binding’, Traffic, 13: 120–30.

Gouzy, A., Y. Poquet, and O. Neyrolles. 2014. ‘Nitrogen metabolism in Mycobacterium tuberculosis physiology and virulence’, Nat Rev Microbiol, 12: 729–37.

Greub, G., B. La Scola, and D. Raoult. 2004. ‘Amoebae-resisting bacteria isolated from human nasal swabs by amoebal coculture’, Emerg Infect Dis, 10: 470–7.

Hagedorn, M., K. H. Rohde, D. G. Russell, and T. Soldati. 2009. ‘Infection by tubercular mycobacteria is spread by nonlytic ejection from their amoeba hosts’, Science, 323: 1729–33.

Hagedorn, M., and T. Soldati. 2007. ‘Flotillin and RacH modulate the intracellular immunity of Dictyostelium to Mycobacterium marinum infection’, Cell Microbiol, 9: 2716–33.

Harrison, C. F., S. Kicka, V. Trofimov, K. Berschl, H. Ouertatani-Sakouhi, N. Ackermann, C. Hedberg, P. Cosson, T. Soldati, and H. Hilbi. 2013. ‘Exploring anti-bacterial compounds against intracellular Legionella’, PLoS One, 8: e74813.

Hennig, M., B. D. Darimont, J. N. Jansonius, and K. Kirschner. 2002. ‘The catalytic mechanism of indole-3-glycerol phosphate synthase: crystal structures of complexes of the enzyme from Sulfolobus solfataricus with substrate analogue, substrate, and product’, J Mol Biol, 319: 757–66.

Hilbi, H., S. S. Weber, C. Ragaz, Y. Nyfeler, and S. Urwyler. 2007. ‘Environmental predators as models for bacterial pathogenesis’, Environ Microbiol, 9: 563–75.

Iyer, R., Z. Wu, P. M. Woster, and A. H. Delcour. 2000. ‘Molecular basis for the polyamine-ompF porin interactions: inhibitor and mutant studies’, J Mol Biol, 297: 933–45.

Jayaraman, G, A Sivakumar, T Panneerselvam, S Hemalatha, and JS Emmanuel. 2010. ‘A comparative study on the potentials of calcineurin inhibitors by docking’, Drug Invention Today, 2.

Kicka, S., V. Trofimov, C. Harrison, H. Ouertatani-Sakouhi, J. McKinney, L. Scapozza, H. Hilbi, P. Cosson, and T. Soldati. 2014. ‘Establishment and validation of whole-cell based fluorescence assays to identify anti-mycobacterial compounds using the Acanthamoeba castellanii-Mycobacterium marinum host-pathogen system’, PLoS One, 9: e87834.

Kirchmair, J., S. Distinto, P. Markt, D. Schuster, G. M. Spitzer, K. R. Liedl, and G. Wolber. 2009. ‘How to optimize shape-based virtual screening: choosing the right query and including chemical information’, J Chem Inf Model, 49: 678–92.

Komurov, K., S. Dursun, S. Erdin, and P. T. Ram. 2012. ‘NetWalker: a contextual network analysis tool for functional genomics’, BMC Genomics, 13: 282.

Lipinski, C. A. 2016. ‘Rule of five in 2015 and beyond: Target and ligand structural limitations, ligand chemistry structure and drug discovery project decisions’, Adv Drug Deliv Rev, 101: 34–41.

Lipinski, Christopher A, Franco Lombardo, Beryl W Dominy, and Paul J Feeney. 1997. ‘Experimental and computational approaches to estimate solubility and permeability in drug discovery and development settings’, Adv Drug Deliv Rev, 23: 3–25.

Lipinski, Christopher, and Andrew Hopkins. 2004. ‘Navigating chemical space for biology and medicine’, Nature, 432: 855.

Long, S. B., E. B. Campbell, and R. Mackinnon. 2005. ‘Crystal structure of a mammalian voltage-dependent Shaker family K+ channel’, Science, 309: 897–903.

Loomis, W. F. 2015. ‘Genetic control of morphogenesis in Dictyostelium’, Dev Biol, 402: 146–61.

Loregian, A., and G. Palu. 2013. ‘How academic labs can approach the drug discovery process as a way to synergize with big pharma’, Trends Microbiol, 21: 261–4.

Lu, X., K. Huang, and Q. You. 2011. ‘Enoyl acyl carrier protein reductase inhibitors: a patent review (2006 - 2010)’, Expert Opin Ther Pat, 21: 1007–22.

Macarron, R., M. N. Banks, D. Bojanic, D. J. Burns, D. A. Cirovic, T. Garyantes, D. V. Green, R. P. Hertzberg, W. P. Janzen, J. W. Paslay, U. Schopfer, and G. S. Sittampalam. 2011. ‘Impact of high-throughput screening in biomedical research’, Nat Rev Drug Discov, 10: 188–95.

Molmeret, M., M. Horn, M. Wagner, M. Santic, and Y. Abu Kwaik. 2005. ‘Amoebae as training grounds for intracellular bacterial pathogens’, Appl Environ Microbiol, 71: 20–8.

Nichols, J. M., D. Veltman, and R. R. Kay. 2015. ‘Chemotaxis of a model organism: progress with Dictyostelium’, Curr Opin Cell Biol, 36: 7–12.

Nilsson, M. T., W. W. Krajewski, S. Yellagunda, S. Prabhumurthy, G. N. Chamarahally, C. Siddamadappa, B. R. Srinivasa, S. Yahiaoui, M. Larhed, A. Karlen, T. A. Jones, and S. L. Mowbray. 2009. ‘Structural basis for the inhibition of Mycobacterium tuberculosis glutamine synthetase by novel ATP-competitive inhibitors’, J Mol Biol, 393: 504–13.

Ouertatani-Sakouhi, H., S. Kicka, G. Chiriano, C. F. Harrison, H. Hilbi, L. Scapozza, T. Soldati, and P. Cosson. 2017. ‘Inhibitors of Mycobacterium marinum virulence identified in a Dictyostelium discoideum host model’, PLoS One, 12: e0181121.

Payandeh, J., T. Scheuer, N. Zheng, and W. A. Catterall. 2011. ‘The crystal structure of a voltage-gated sodium channel’, Nature, 475: 353–8.

Pedicord, D. L., M. J. Flynn, C. Fanslau, M. Miranda, L. Hunihan, B. J. Robertson, B. C. Pearce, X. C. Yu, R. S. Westphal, and Y. Blat. 2011. ‘Molecular characterization and identification of surrogate substrates for diacylglycerol lipase alpha’, Biochem Biophys Res Commun, 411: 809–14.

Perez, Freddy, Bertha Gomez, Giovanni Ravasi, and Massimo Ghidinelli. 2015. ‘Progress and challenges in implementing HIV care and treatment policies in Latin America following the treatment 2.0 initiative’, BMC public health, 15: 1260.

Pethe, Kevin, Patricia C Sequeira, Sanjay Agarwalla, Kyu Rhee, Kelli Kuhen, Wai Yee Phong, Viral Patel, David Beer, John R Walker, and Jeyaraj Duraiswamy. 2010. ‘A chemical genetic screen in Mycobacterium tuberculosis identifies carbon-source-dependent growth inhibitors devoid of in vivo efficacy’, Nature communications, 1: 57.

Prashar, A., and M. R. Terebiznik. 2015. ‘Legionella pneumophila: homeward bound away from the phagosome’, Curr Opin Microbiol, 23: 86–93.

Sakowicz, Roman, Michael S Berdelis, Krishanu Ray, Christine L Blackburn, Cordula Hopmann, D John Faulkner, and Lawrence SB Goldstein. 1998. ‘A marine natural product inhibitor of kinesin motors’, Science, 280: 292–95.

Salsi, E., A. S. Bayden, F. Spyrakis, A. Amadasi, B. Campanini, S. Bettati, T. Dodatko, P. Cozzini, G. E. Kellogg, P. F. Cook, S. L. Roderick, and A. Mozzarelli. 2010. ‘Design of O-acetylserine sulfhydrylase inhibitors by mimicking nature’, J Med Chem, 53: 345–56.

Sarkar, Sovan, Ethan O Perlstein, Sara Imarisio, Sandra Pineau, Axelle Cordenier, Rebecca L Maglathlin, John A Webster, Timothy A Lewis, Cahir J O’Kane, and Stuart L Schreiber. 2007. ‘Small molecules enhance autophagy and reduce toxicity in Huntington’s disease models’, Nature chemical biology, 3: 331.

Schalk-Hihi, C., C. Schubert, R. Alexander, S. Bayoumy, J. C. Clemente, I. Deckman, R. L. DesJarlais, K. C. Dzordzorme, C. M. Flores, B. Grasberger, J. K. Kranz, F. Lewandowski, L. Liu, H. Ma, D. Maguire, M. J. Macielag, M. E. McDonnell, T. Mezzasalma Haarlander, R. Miller, C. Milligan, C. Reynolds, and L. C. Kuo. 2011. ‘Crystal structure of a soluble form of human monoglyceride lipase in complex with an inhibitor at 1.35 A resolution’, Protein Sci, 20: 670–83.

Scheid, P. 2014. ‘Relevance of free-living amoebae as hosts for phylogenetically diverse microorganisms’, Parasitol Res, 113: 2407–14.

Sharom, F. J. 2008. ‘ABC multidrug transporters: structure, function and role in chemoresistance’, Pharmacogenomics, 9: 105–27.

Sharp, Swee Y, Kathy Boxall, Martin Rowlands, Chrisostomos Prodromou, S Mark Roe, Alison Maloney, Marissa Powers, Paul A Clarke, Gary Box, and Sharon Sanderson. 2007. ‘In vitro biological characterization of a novel, synthetic diaryl pyrazole resorcinol class of heat shock protein 90 inhibitors’, Cancer research, 67: 2206–16.

Shelat, A. A., and R. K. Guy. 2007. ‘Scaffold composition and biological relevance of screening libraries’, Nat Chem Biol, 3: 442–6.

Shen, H., F. Wang, Y. Zhang, Q. Huang, S. Xu, H. Hu, J. Yue, and H. Wang. 2009. ‘A novel inhibitor of indole-3-glycerol phosphate synthase with activity against multidrug-resistant Mycobacterium tuberculosis’, Febs j, 276: 144–54.

Sieber, Matthias, and Ria Baumgrass. 2009. ‘Novel inhibitors of the calcineurin/NFATc hub - alternatives to CsA and FK506?’, Cell Communication and Signaling : CCS, 7: 25–25.

Simon, S., and H. Hilbi. 2015. ‘Subversion of Cell-Autonomous Immunity and Cell Migration by Legionella pneumophila Effectors’, Front Immunol, 6: 447.

Sindhikara, D., and K. Borrelli. 2018. ‘High throughput evaluation of macrocyclization strategies for conformer stabilization’, Sci Rep, 8: 6585.

Slepikas, L., G. Chiriano, R. Perozzo, S. Tardy, A. Kranjc, O. Patthey-Vuadens, H. Ouertatani-Sakouhi, S. Kicka, C. F. Harrison, T. Scrignari, K. Perron, H. Hilbi, T. Soldati, P. Cosson, E. Tarasevicius, and L. Scapozza. 2016. ‘In Silico Driven Design and Synthesis of Rhodanine Derivatives as Novel Antibacterials Targeting the Enoyl Reductase InhA’, J Med Chem, 59: 10917–28.

Solomon, J. M., G. S. Leung, and R. R. Isberg. 2003. ‘Intracellular replication of Mycobacterium marinum within Dictyostelium discoideum: efficient replication in the absence of host coronin’, Infect Immun, 71: 3578–86.

Songane, M., J. Kleinnijenhuis, B. Alisjahbana, E. Sahiratmadja, I. Parwati, M. Oosting, T. S. Plantinga, L. A. Joosten, M. G. Netea, T. H. Ottenhoff, E. van de Vosse, and R. van Crevel. 2012. ‘Polymorphisms in autophagy genes and susceptibility to tuberculosis’, PLoS One, 7: e41618.

Steinert, M., and K. Heuner. 2005. ‘Dictyostelium as host model for pathogenesis’, Cell Microbiol, 7: 307–14.

Sterling, T., and J. J. Irwin. 2015. ‘ZINC 15--Ligand Discovery for Everyone’, J Chem Inf Model, 55: 2324–37.

Sundaramurthy, V., R. Barsacchi, M. Chernykh, M. Stoter, N. Tomschke, M. Bickle, Y. Kalaidzidis, and M. Zerial. 2014. ‘Deducing the mechanism of action of compounds identified in phenotypic screens by integrating their multiparametric profiles with a reference genetic screen’, Nat Protoc, 9: 474–90.

Supino, Rosanna, Giovanna Petrangolini, Graziella Pratesi, Monica Tortoreto, Enrica Favini, Laura Dal Bo, Patrizia Casalini, Enrico Radaelli, Anna Cleta Croce, and Giovanni Bottiroli. 2008. ‘Antimetastatic effect of a small-molecule vacuolar H+-ATPase inhibitor in in vitro and in vivo preclinical studies’, Journal of Pharmacology and Experimental Therapeutics, 324: 15–22.

Swann, S. L., S. P. Brown, S. W. Muchmore, H. Patel, P. Merta, J. Locklear, and P. J. Hajduk. 2011. ‘A unified, probabilistic framework for structure- and ligand-based virtual screening’, J Med Chem, 54: 1223–32.

Tardy, S., A. Orsato, L. Mologni, W. H. Bisson, C. Donadoni, C. Gambacorti-Passerini, L. Scapozza, D. Gueyrard, and P. G. Goekjian. 2014. ‘Synthesis and biological evaluation of benzo[4,5]imidazo[1,2-c]pyrimidine and benzo[4,5]imidazo[1,2-a]pyrazine derivatives as anaplastic lymphoma kinase inhibitors’, Bioorg Med Chem, 22: 1303–12.

Tosetti, N., A. Croxatto, and G. Greub. 2014. ‘Amoebae as a tool to isolate new bacterial species, to discover new virulence factors and to study the host-pathogen interactions’, Microb Pathog, 77: 125–30.

Trofimov, V., S. Kicka, S. Mucaria, N. Hanna, F. Ramon-Olayo, L. V. Del Peral, J. Lelievre, L. Ballell, L. Scapozza, G. S. Besra, J. A. G. Cox, and T. Soldati. 2018. ‘Antimycobacterial drug discovery using Mycobacteria-infected amoebae identifies anti-infectives and new molecular targets’, Sci Rep, 8: 3939.

VanderVen, B. C., R. J. Fahey, W. Lee, Y. Liu, R. B. Abramovitch, C. Memmott, A. M. Crowe, L. D. Eltis, E. Perola, D. D. Deininger, T. Wang, C. P. Locher, and D. G. Russell. 2015. ‘Novel inhibitors of cholesterol degradation in Mycobacterium tuberculosis reveal how the bacterium’s metabolism is constrained by the intracellular environment’, PLoS Pathog, 11: e1004679.

Vidal, D., M. Thormann, and M. Pons. 2005. ‘LINGO, an efficient holographic text based method to calculate biophysical properties and intermolecular similarities’, J Chem Inf Model, 45: 386–93.

Walker, Edward H, Michael E Pacold, Olga Perisic, Len Stephens, Philip T Hawkins, Matthias P Wymann, and Roger L Williams. 2000. ‘Structural determinants of phosphoinositide 3-kinase inhibition by wortmannin, LY294002, quercetin, myricetin, and staurosporine’, Molecular cell, 6: 909–19.

Wambaugh, M. A., V. P. S. Shakya, A. J. Lewis, M. A. Mulvey, and J. C. S. Brown. 2017. ‘High-throughput identification and rational design of synergistic small-molecule pairs for combating and bypassing antibiotic resistance’, PLoS Biol, 15: e2001644.

Westermaier, Y., X. Barril, and L. Scapozza. 2015. ‘Virtual screening: an in silico tool for interlacing the chemical universe with the proteome’, Methods, 71: 44–57.

Wu, Y., Y. Yang, S. Ye, and Y. Jiang. 2010. ‘Structure of the gating ring from the human large-conductance Ca(2+)-gated K(+) channel’, Nature, 466: 393–7.

Yang, L., S. Wang, B. Sung, G. Lim, and J. Mao. 2008. ‘Morphine induces ubiquitin-proteasome activity and glutamate transporter degradation’, J Biol Chem, 283: 21703–13.

Yang, Yili, Robert L Ludwig, Jane P Jensen, Shervon A Pierre, Maxine V Medaglia, Ilia V Davydov, Yassamin J Safiran, Pankaj Oberoi, John H Kenten, and Andrew C Phillips. 2005. ‘Small molecule inhibitors of HDM2 ubiquitin ligase activity stabilize and activate p53 in cells’, Cancer cell, 7: 547–59.

Yoshida, K., K. Yamaguchi, A. Mizuno, Y. Unno, A. Asai, T. Sone, H. Yokosawa, A. Matsuda, M. Arisawa, and S. Shuto. 2009. ‘Three-dimensional structure-activity relationship study of belactosin A and its stereo- and regioisomers: development of potent proteasome inhibitors by a stereochemical diversity-oriented strategy’, Org Biomol Chem, 7: 1868–77.

Zoraghi, R., L. Worrall, R. H. See, W. Strangman, W. L. Popplewell, H. Gong, T. Samaai, R. D. Swayze, S. Kaur, M. Vuckovic, B. B. Finlay, R. C. Brunham, W. R. McMaster, M. T. Davies-Coleman, N. C. Strynadka, R. J. Andersen, and N. E. Reiner. 2011. ‘Methicillin-resistant Staphylococcus aureus (MRSA) pyruvate kinase as a target for bis-indole alkaloids with antibacterial activities’, J Biol Chem, 286: 44716–25.

